# Effect of nuclear electrostatic potential on the nucleoplasmic distribution of histones and nucleosome stability

**DOI:** 10.64898/2026.07.23.738035

**Authors:** Yuqi Guo, Fei He, Xin Dai, Jiaying Yu, Yixuan Li, Fenfang Li, Cheng-han Yu, Ying Wai Chan, Andrew W. Holle, Esteban Hoijman, Artem K. Efremov

## Abstract

Differences in the density of heterochromatic and euchromatic regions are often assumed to play an important role in determining the transcriptional state of chromatin by influencing its accessibility to transcription factors. Yet, experiments show that even the most densely packed chromatin domains are readily accessible to transcription factors. Thus, the molecular mechanisms underlying differences in the transcriptional states of hetero- and euchromatin remain poorly understood. In this study, using electrically charged mEGFP probes, we demonstrated that heterochromatin and euchromatin differ not only in density but also in the magnitude of the local electrostatic potential. Furthermore, the nucleoplasmic distribution of electrically charged proteins, such as histone-chaperone complexes, was found to correlate with the local electrostatic potential. Estimates based on experimental data have also shown that the difference in electrostatic potentials between heterochromatic and euchromatic regions could lead to unequal nucleosome stability in them, which was successfully confirmed experimentally. Subsequent theoretical calculations showed that this could shift the balance in the DNA-binding competition between transcription factors and nucleosomes, thereby explaining experimental observations that heterochromatin is less transcriptionally active than euchromatin. Overall, our study suggests the existence of a nuclear electrostatic potential-mediated pathway that may be involved in the regulation of gene transcription.

## INTRODUCTION

Chromatin organization plays an important role in the regulation of gene transcription [1]. It has been previously shown that numerous components of the cell nucleus, including chromatin-modifying and -reading enzymes [2], histone-binding chaperones [3, 4], RNA and RNA-binding proteins [5, 6], as well as proteins associated with the formation of heterochromatin [7, 8] and topologically associated domains [9], engage into complex interactions with each other, which, in combination with the macromolecular crowding effect [10–12], lead to the formation of multiple membraneless nuclear compartments, including nucleoli, heterochromatin, and euchromatin [2, 5–12]. On a larger scale, this results in a characteristic chromatin pattern in terms of chromosomal territories and compartments, which are separated by interchromatin space enriched in RNA and transcriptional machinery [1, 6, 13–15]. However, due to the extremely complex network of interactions between the aforementioned key factors, there is still no clear understanding of how the phase-separation of chromatin into hetero- and euchromatic domains leads to the experimentally observed difference in their transcriptional states.

Although it is often assumed that genes located in heterochromatic regions are generally more transcriptionally silent due to the limited accessibility of transcription factors (TFs) to densely packed chromatin, experiments have shown that dense heterochromatin has only a modest effect on the diffusion coefficients of proteins or their complexes, as well as on their accessibility to the heterochromatin volume, compared to more sparse euchromatic regions [12]. Indeed, fluorescence recovery after photobleaching (FRAP) experiments have shown that even highly condensed mitotic chromosomes can be readily accessible to TFs [16]. Furthermore, phase-separation processes involved in the formation and maintenance of heterochromatic regions, such as those mediated by heterochromatin protein 1 (HP1), were also found to have little effect on the accessibility of heterochromatin volume to core TFs, such as TFIIB [7]. Although it has been previously proposed that more accessible DNA surfaces in euchromatin regions may be scanned more efficiently by TFs than in heterochromatic regions [12], this argument does not explain why, when TFs find a specific binding site in the promoter regions of genes, they will bind DNA more efficiently in euchromatic regions compared to heterochromatic regions.

On the other hand, many experimental studies show that TFs are, in fact, involved in DNA-binding competition with nucleosomes [17–21], and indeed, the stability of nucleosomes associated with promoter regions has been experimentally demonstrated to have a profound effect on the transcriptional activity of downstream genes [22]. Thus, it can be speculated that nucleosome stability, rather than the chromatin packaging density itself, is one of the main elements responsible for the difference between the transcriptional states of genes in heterochromatic and euchromatic regions. However, the precise molecular mechanisms that could potentially mediate the differential stabilization of nucleosomes in heterochromatic and euchromatic regions remain largely unknown [23].

Previous theoretical studies have predicted that electrostatic forces could potentially play an important role in chromatin organization and in the regulation of gene transcription by influencing nucleosome stability [24, 25], since most DNA-binding proteins and chromatin-shaping biomolecules, such as histones and TFs, carry high electrical charges. In particular, it has been theoretically predicted that the Donnan electrostatic potential between the cytosol and the nucleoplasm (i.e., the nuclear electrostatic potential), arising from the confinement of highly negatively charged DNA in the nucleus [see Figure 1(a), Appendix A and ref. [26–28]], may have a significant impact on nucleosome stability by changing the binding free energy of histone octamers to DNA by ∼ 4 − 10 k_B_T [25]. Such a strong change in nucleosome stability may be significant enough to tip the balance in the DNA-binding competition between nucleosomes and TFs, as suggested by previous studies [29].

**FIG. 1.**
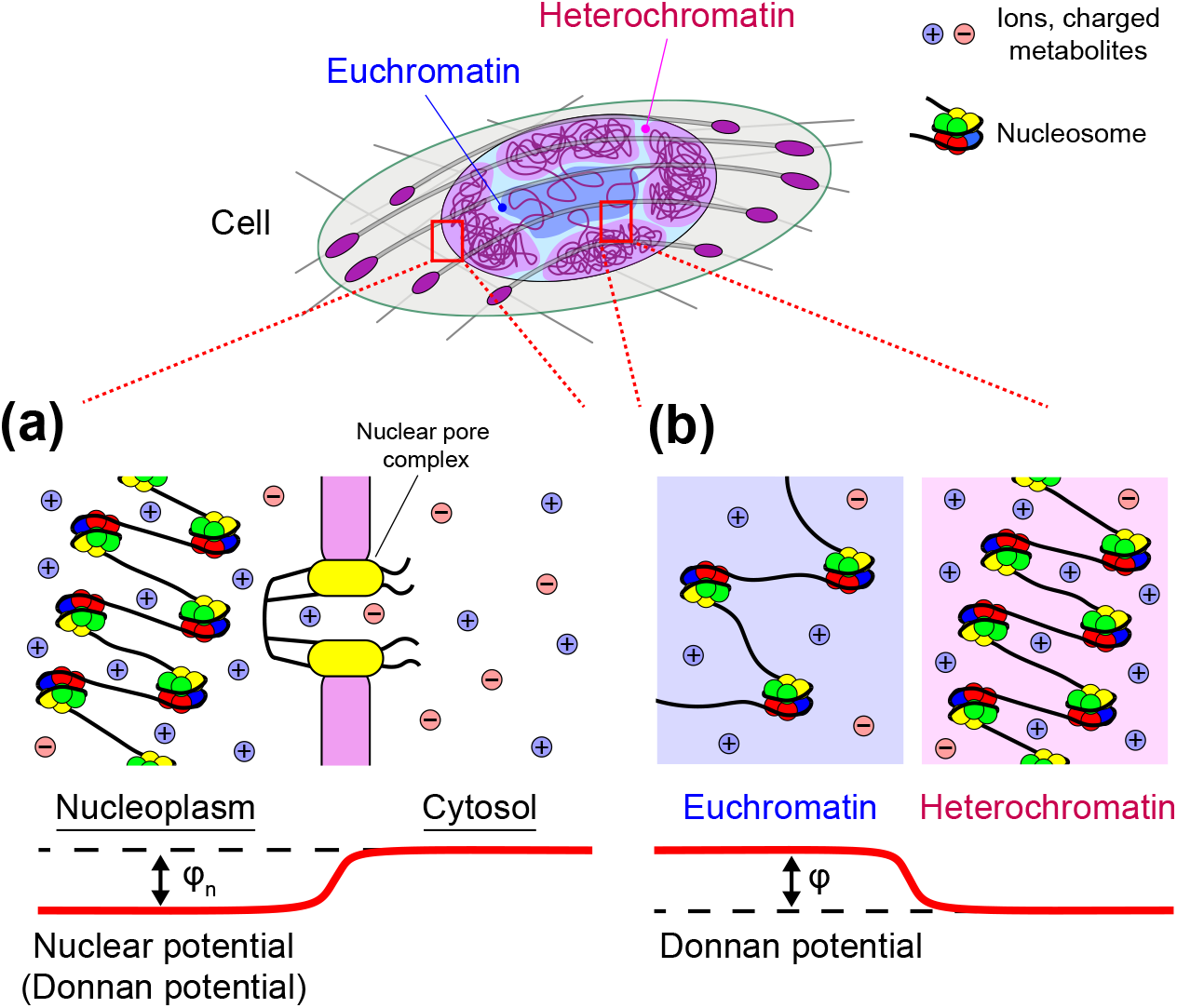
Nuclear electrostatic potential as a manifestation of the Donnan effect. **(a)** Negatively electrically charged DNA confined to the nucleus results in the appearance of a Donnan electrostatic potential between the nucleus and the cytosol (i.e., nuclear electrostatic potential). **(b)** Phase-separation of chromatin into dense heterochromatic regions and sparse euchromatic regions can potentially lead to spatial heterogeneity of the nuclear electrostatic potential.

Following this logic, it is tempting to speculate that the nuclear electrostatic potential may also be one of the key factors responsible for the different nucleosome stability in heterochromatic and euchromatic regions, since a very similar Donnan electrostatic potential should arise between these regions, which have very different chromatin densities, see Figure 1(b). Indeed, although the short Debye screening length (∼ 0.7 nm [30]) suggests that the electrostatic potential near electrically charged nuclear components (DNA, nucleosomes, etc.) should drop to low values at distances of a few nanometers, the high chromatin density in living cell nuclei and strong thermal fluctuations affecting chromatin conformation give rise to an electrostatic field, which fluctuates around a non-zero level determined by the average local electric charge density. This field may potentially be strong enough to influence the stability of electrically charged nuclear complexes, such as nucleosomes. Yet, there is currently a lack of experimental evidence to support this hypothesis, although recent experimental studies suggest that electrostatic interactions may indeed play an important role in chromatin organization, for example in mitotic chromosomes [31].

To experimentally test it and measure the spatial heterogeneity of the nuclear electrostatic potential, we developed an experimental method to calibrate the energies of electrostatic and volume-exclusion interactions between electrically charged mEGFP-based fluorescent probes and the nuclear microenvironment. Using these molecular probes, it was found that the nuclear electrostatic potential strongly correlates with the nucleoplasmic distribution of DNA in cells grown on rigid substrates. Moreover, growing cells on substrates with different elastic properties, or subjecting them to geometric constraints or hypoosmotic shock, showed that the mean nuclear electrostatic potential behaves in a mechanosensitive manner, demonstrating a strong correlation with the size of the cell nucleus. In addition, FRAP experiments performed on cells expressing fluorescently-labelled histones showed that the local nucleoplasmic density of the freely diffusing fraction of histones correlates with the local strength of the nuclear electrostatic potential, suggesting a significant effect of the latter on electrically charged nucleoplasmic proteins. Further evaluation based on the experimentally measured spatial distribution of the nuclear electrostatic potential revealed that the difference between the electrostatic potentials of heterochromatic and euchromatic regions is large enough to strongly affect the stability of nucleosomes and to a lesser extent the stability of tetrasomes, which likely leads to differential stabilization of these nucleoprotein complexes. This prediction was successfully confirmed in chromatin digestion experiments using MNase, as well as by directly measuring the fluorescent signal ratio of fluorescently-labelled H2B and H3 histones in cell nuclei. Finally, theoretical calculations based on the above experiments further suggested that differential stabilization of nucleosomes in hetero- and euchromatic regions by nuclear electrostatic potential is one of the key elements influencing the binding of TFs to DNA, providing new insights into potential molecular mechanisms underlying the experimentally observed difference in the transcriptional states of these chromatin regions.

## RESULTS

### mEGFP probes for measuring nuclear electrostatic potential

In the crowded microenvironment of the cell nucleus, chromatin undergoes a wide range of interactions with surrounding nuclear components, leading to phase-separation of chromatin into hetero- and euchromatin domains and the formation of chromosomal territories, the spatial organization of which is dictated by the geometric constraints imposed on long DNA molecules by the tight nuclear space, as well as the anchoring of various parts of chromatin to the nuclear envelope. In this context, non-specific interactions, such as electrostatic and volume-exclusion interactions, are of particular interest, since they affect all nuclear components and, therefore, impact numerous molecular processes taking place in the nucleus.

To experimentally assess the relative strength of such nonspecific interactions influencing the distribution of small charged proteins in the nucleoplasm, we employed electrically charged fluorescent probes based on mEGFP, which is known to be an inert and purely diffusive protein, interacting with the surrounding intracellular microenvironment in a non-specific manner [32], drawing upon findings from previous studies demonstrating that such probes can be used to evaluate the strength of electrostatic interactions and the electrostatic potential of the cell membrane and intracellular organelles [33–35]. In particular, in addition to the negatively charged wildtype mEGFP (mEGFP WT, *q* = − 7.3*e*), we created mEGFP probes with nearly net neutral electrical charge (mEGFP+3RK, *q* = − 1.3*e*) and net positive electrical charge (mEGFP+6RK, *q* = +4.7*e*) by adding positively charged aminoacids (R and K) to the C-terminus of mEGFP WT, as schematically shown in Figure 2(a). Since the addition of such short peptides to mEGFP has little effect on the overall mEGFP volume (*<* 7% volume increase), the measured difference in the nucleoplasmic distribution of such mEGFP probes can be largely attributed to changes in their electrostatic interactions with nuclear components, allowing a comparison of the relative strengths of electrostatic and volume-exclusion interactions.

**FIG. 2.**
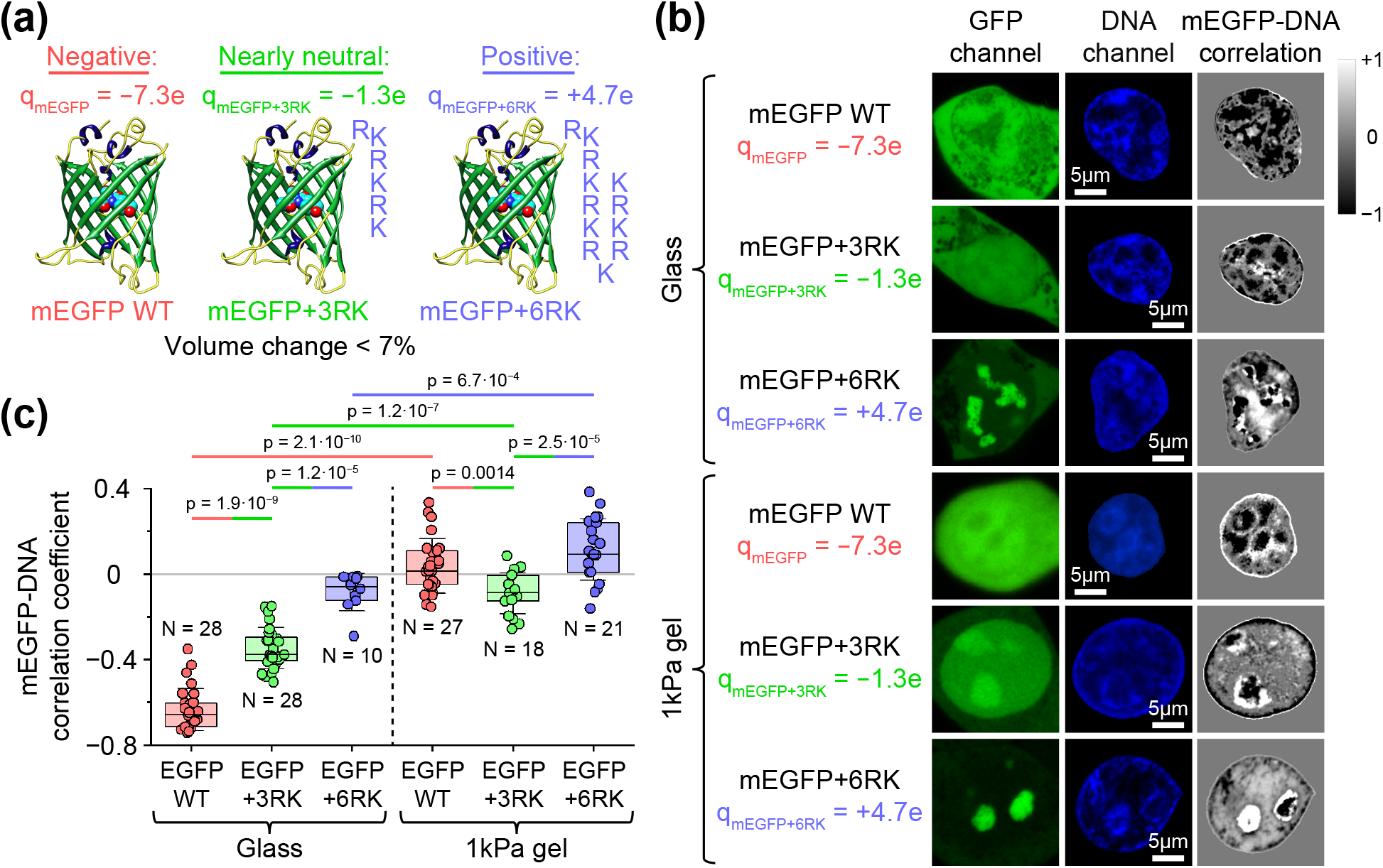
Effect of net electrical charge of mEGFP probes on their nucleoplasmic distribution. **(a)** mEGFP probes with a net negative, neutral and positive electrical charge tested in experiments. The net electrical charge of the mEGFP probes was estimated based on the measured pH level in HEK293T cell nuclei using the pHrodo Red indicator, see Figure S1, SI. **(b)** Representative images of nucleoplasmic distribution of mEGFP probes and DNA in the nuclei of HEK293T cells grown on fibronectin-coated glass slides and 1 kPa gel (see Figure S2, SI, for large-scale images of the same cells). Grayscale images of cell nuclei show the correlation between the GFP and DNA channels. In the case of cells grown on fibronectin-coated glass slides, the nucleoplasmic distribution of the negatively charged mEGFP WT probes was strongly anticorrelated with DNA, whereas the nearly neutral mEGFP+3RK probes showed significantly reduced anticorrelation with DNA. **(c)** Average mEGFP-DNA correlation coefficient measured for different types of mEGFP probes. The graph shows that electrostatic interactions between chromatin and mEGFP probes play an important role in governing the nucleoplasmic distribution of the latter. *N* indicates the number of cells processed under the corresponding experimental conditions. Boxes represent the interquartile range, error bars indicate SD. Bars above the graph show pairwise statistical differences between corresponding experimental data, calculated using Mann-Whitney test.

After transfection of HEK293T cells grown on fibronectin-coated glass slides with plasmids encoding the mEGFP probes, all three mEGFP probes were found to behave quite differently, see Figure 2(b) and S2, SI. In particular, mEGFP WT probes were found to avoid regions of high DNA density, showing a strong negative correlation with DNA distribution (mEGFP-DNA correlation coefficient: − 0.635 ± 0.019, mean ± SEM, *N* = 28 cells), see Figure 2(b,c). On the other hand, mEGFP+3RK probes demonstrated a significantly lower negative correlation with DNA (mEGFP-DNA correlation coefficient: − 0.347 ± 0.019, mean ± SEM, *N* = 28 cells), exhibiting a more uniform nucleoplasmic distribution, see Figure 2(b,c). In contrast, net positively charged mEGFP+6RK probes tended to form aggregates that showed no correlation with DNA distribution.

Measurements of the relative expression levels of the three mEGFP constructs in HEK293T cells using Western blotting, FACS, and confocal microscopy revealed that the more uniform nucleoplasmic distribution of mEGFP+3RK and the aggregation tendency of mEGFP+6RK may not have been due to their higher expression levels, as they were expressed at levels lower than that of mEGFP WT, see Figure S3, SI. Furthermore, by weakly permeabilizing cells using PBFT solution (see Methods) and monitoring the time-dependent change in the total nuclear fluorescent signals of the mEGFP constructs, it was found that both mEGFP WT and mEGFP+3RK rapidly exited the nuclei of permeabilized cells (Figure S4, SI), indicating their predominantly weak, non-specific interaction with the nuclear microenvironment. On the other hand, mEGFP+6RK was retained in the nuclei for a significantly longer period compared to mEGFP WT and mEGFP+3RK, suggesting a stronger interaction of mEGFP+6RK probes with nuclear components.

Overall, the above results suggest that non-specific electrostatic interactions play an important role in regulating the nucleoplasmic distribution of even weakly charged proteins such as mEGFP. In particular, a comparison of DNA-mEGFP correlation coefficients for mEGFP WT and mEGFP+3RK constructs demonstrates that electrostatic and volume-exclusion interactions play an equally important role in determining the nucleoplasmic distribution of mEGFP WT, which can therefore be used to probe the strength of these interactions in different parts of the nucleus. Indeed, fast diffusion of mEGFP WT probes throughout the cell nucleus and cytoplasm, makes it possible for them to reach equilibrium in a matter of seconds (∼ 0.08 − 15 s, see [32, 36]), which is orders of magnitude faster than large-scale chromatin reorganization processes (from tens of minutes to hours [37]). Thus, at each moment in time, mEGFP WT probes are in quasi-equilibrium with their microenvironment, which allows for precise measurements of their interaction with it based on Boltzmann’s law.

To calibrate mEGFP WT probes, we used an experimental procedure based on comparing their nucleoplasmic distribution with that of another fluorescent protein, mScarlet, which has a nearly identical size but carries a different net electrical charge (*q* = − 3.6*e*) and therefore experiences the same strength of volume-exclusion interactions but a different strength of electrostatic interactions with the surrounding microenvironment. Thus, by measuring the difference in the nucleoplasmic distributions of mEGFP WT and mScarlet expressed in the same HEK293T cells, one can obtain accurate estimates of the electrostatic and volume-exclusion interaction energies of the mEGFP WT probe with the surrounding microenvironment in the nucleus, see Appendix B for details. As a result, based on the Boltzmann’s law, it was found that these interaction energies exhibit a linear correlation with the logarithm of the local fluorescent intensity of mEGFP WT probes in the nucleus, which allows them to be used to map the spatial distribution of nuclear electrostatic potential across cell nuclei, see Figure S6(c,d), SI.

Notably, experimental measurements of the mean volume-exclusion interaction energy of mEGFP WT probes with the nuclear microenvironment based on the calibration curves revealed that 2.83 ± 0.12% (mean ± SEM, *N* = 21 cells) of the nuclear volume (excluding nucleoli) in HEK293T cells grown on glass slides should be occupied by chromatin, see Appendix C. This result is in good agreement with direct estimates of the fraction of nuclear volume occupied by nucleosomes (∼ 2.3%), which can be obtained from the approximate volume occupied by each nucleosome, the total number of nucleosomes, and the nuclear volume measured in experiments (Appendix C). Thus, it can be concluded that the calibration curves shown in Figure S6(c,d), SI, provide physically realistic values of the interaction energy of mEGFP WT probes with the surrounding microenvironment.

#### Mechanosensitive behaviour of nuclear electrostatic potential

Previous experimental studies show that the elasticity of the substrate on which cells are grown greatly influences cell morphology – on rigid substrates, cells take on a more flattened shape compared to cells grown on a soft substrate, see Figure 3(b) and ref. [38]. This change in cell morphology is accompanied by a corresponding change in nuclear volume, which positively correlates with cell volume [25, 38]. This results in a significant difference in the average chromatin density in cells grown on soft and rigid substrates, which may affect the average value of the nuclear electrostatic potential, see Appendix A.

**FIG. 3.**
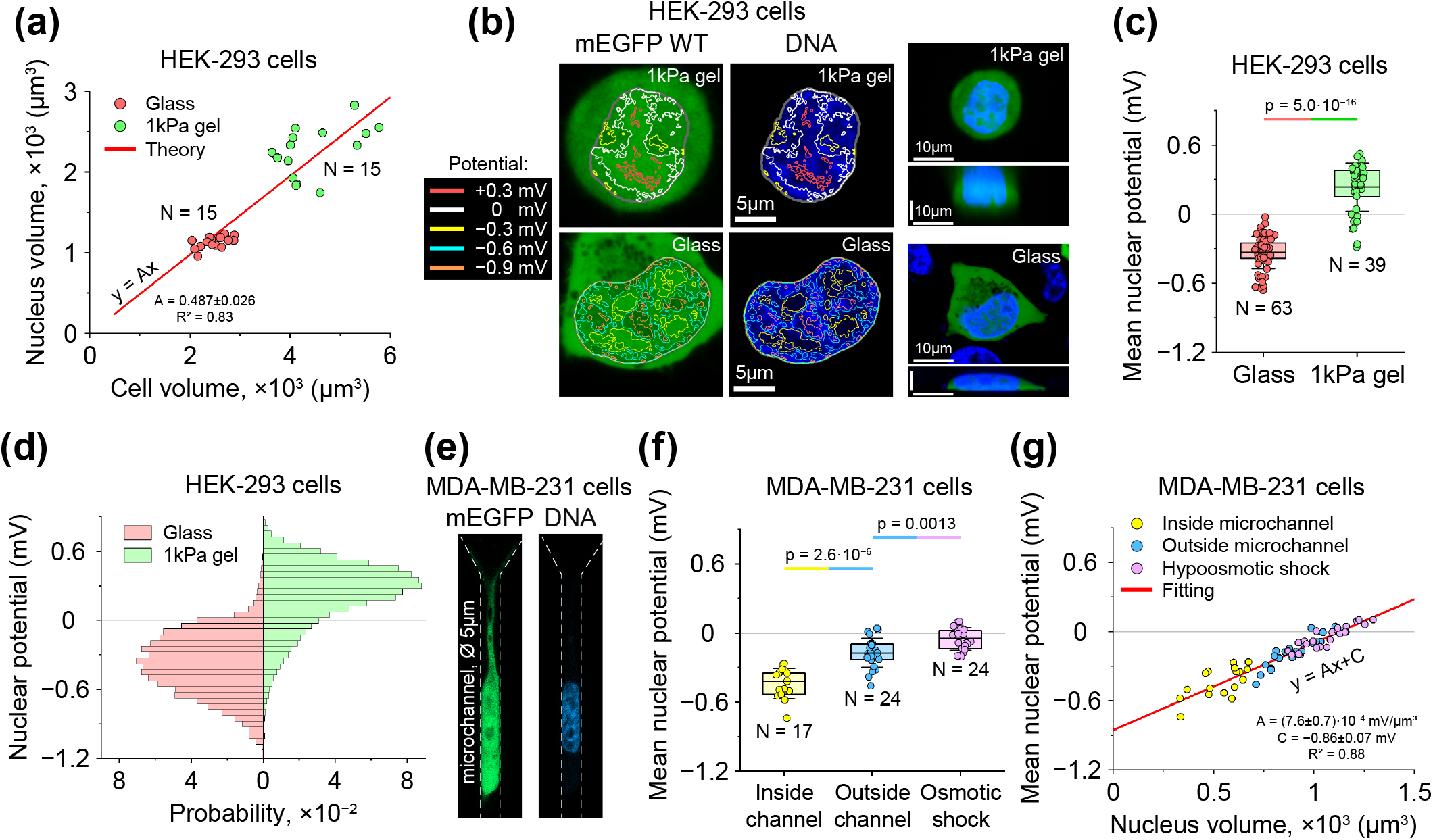
Mechanosensitive behaviour of nuclear electrostatic potential. **(a)** Correlation between cell and nuclear volumes of HEK293T cells. Cell and nuclear volumes can change by up to two times depending on the elasticity of the substrate on which cells are grown. Each data point represents the experimentally measured volumes of a HEK293T cell and its nucleus. Cells were grown on either fibronectin-coated glass or 1 kPa gel. *N* = 15 cells were measured for each experimental condition. The solid line represents the fitting of the experimental data to Eq. (D4). **(b)** Representative images of the spatial distribution of nuclear electrostatic potential measured using the mEGFP WT probe in HEK293T cells. The figure shows equipotential contours calculated based on the calibration curve of mEGFP WT [Figure S6(c), SI]. Nuclear electrostatic potential depends on the elasticity of the substrate on which the cells were grown. Moreover, the equipotential contours appeared to be strongly correlated with the local DNA density in cells grown on fibronectin-coated glass slides. The panels on the right show vertical and horizontal sections of the cells. **(c, d)** Mean nuclear electrostatic potential (c) and histogram of its probability distribution (d) measured in HEK293T cell nuclei. Measurements shown in panels (c, d) were performed in *N* = 63 and *N* = 39 cells, respectively. **(e)** MDA-MB-231 cell expressing the mEGFP WT probe and stained with Hoechest dye migrating through a 5 *µ*m wide microchannel. **(f, g)** Mean nuclear electrostatic potential (f) and its correlation with nuclear volume (g) in MDA-MB-231 cells migrating through 5 *µ*m wide microchannels and in unconfined MDA-MB-231 cells under normal conditions or hypoosmotic shock. The graph shows that there is a linear correlation between the volume of the nucleus and its mean electrostatic potential. Measurements were performed in *N* = 17, *N* = 24 and *N* = 24 cells, respectively. Boxes in panels (c, f) represent the interquartile range, error bars indicate SD. Bars above the graphs in panels (c, f) show pairwise statistical differences between corresponding experimental data, calculated using Mann-Whitney test. Fitting parameter values in panels (a, g) are presented as mean *±* 95% CI.

To experimentally test this phenomenon, we expressed electrically charged mEGFP WT probes in HEK293T cells grown on soft (1 kPa gel) and rigid (glass slides) fibronectin-coated substrates. It was found that cell and nuclear volume were indeed approximately half as large in HEK293T cells grown on glass slides compared to HEK293T cells grown on 1 kPa gel, see Figure 3(a). Furthermore, by measuring the nuclear-cytoplasmic distribution of electrically charged mEGFP WT probes, it was found that the average nuclear electrostatic potential between the nucleus and cytoplasm in HEK293T cells grown on glass (− 0.333 ± 0.018 mV, mean ± SEM, *N* = 63 cells) was significantly different compared to that in HEK293T cells grown on 1 kPa gel (+0.24 ± 0.03 mV, mean ± SEM, *N* = 39 cells), see Figure 3(b-d).

In the case of cells grown on glass, the measured mean nuclear electrostatic potential was in good agreement with theoretical estimates based on the average DNA concentration in the nucleus, see Appendix A. As for HEK293T cells grown on 1 kPa gel, the inversion of the sign of the nuclear electrostatic potential indicates that in this case it may be determined not only by negatively charged DNA, but also by some positively charged molecules located in the nucleus. Accordingly, it was found that in the case of HEK293T cells grown on 1 kPa gel, the correlation coefficient between the distributions of electrically charged mEGFP WT probes and DNA decreased to almost zero (0.036 ± 0.025, mean ± SEM, *N* = 27 cells), indicating that the nucleoplasmic distribution of mEGFP probes is governed by cellular components other than DNA, see Figure 2(b,c).

To test whether the nuclear electrostatic potential can be altered by highly positively charged nuclear proteins, we overexpressed H1.1 linker histones (*q* = +53.0*e*) conjugated to mCherry in HEK293T cells and measured the correlation between the mean nuclear electrostatic potential and the average H1.1-mCherry fluorescence intensity in the nuclei of these cells. It was found that the mean nuclear electrostatic potential indeed increased with higher density of H1.1-mCherry in cell nuclei, suggesting that supercharged nuclear components such as linker histones may have a significant impact on the nuclear electrostatic potential, see Figure S7(a), SI. Although further experiments showed that the expression level of endogenous H1 linker histones was higher in HEK293T cells grown on fibronectin-coated glass slides than in HEK293T cells grown on 1 kPa gel [Figure S7(b), SI], other positively charged nuclear components may be partially responsible for the elevated nuclear electrostatic potential in HEK293T cells grown on 1 kPa gel observed in the experiments, warranting future study.

Another key factor that may have a significant impact on the nuclear electrostatic potential is the nucleus itself. By changing its volume, the nucleus can influence the average density of DNA and other electrically charged nuclear components, which determine the nuclear electrostatic potential, see Appendix A and ref. [25]. To test whether the change in the mean value of the nuclear electrostatic potential could be caused not only by the elastic properties of the substrate but also by the application of geometric constraints to cells leading to a change in the nuclear volume, we used a microchannel cell migration assay [39]. Specifically, we generated a stable MDA-MB231 cell line expressing the electrically charged mEGFP WT probe to monitor changes in nuclear electrostatic potential during cell migration through 5 *µ*m wide microchannels, see Figure 3(e). The nuclear electrostatic potential of MDA-MB-231 cells was found to decrease from − 0.17 ± 0.03 mV (mean ± SEM, *N* = 24 cells) to − 0.44 ± 0.03 mV (mean ± SEM, *N* = 17 cells) when they entered the microchannels, see Figure 3(f). Moreover, these changes were found to be accompanied by a significant decrease in nuclear volume, which appeared to have a linear correlation with the change in the nuclear electrostatic potential, see Figure 3(g).

While the experiments described above required several hours for cells to grow on elastic substrates or migrate into microchannels, we also tested whether changes in nuclear electrostatic potential could be induced in a much shorter period of time by applying hypoosmotic shock to unconfined MDA-MB-231 cells grown on collagen-coated glass slides. Both cell and nuclear volumes were found to increase within 150 s of the onset of hypoosmotic shock, accompanied by an increase in nuclear electrostatic potential to − 0.05 ± 0.02 mV (mean ± SEM, *N* = 24 cells), demonstrating the same linear correlation with nuclear volume as for cells migrating through microchannels, see Figure 3(g).

Overall, the above results suggest that the nuclear electrostatic potential is sensitive to the mechanical properties of the surrounding microenvironment and forces / geometric constraints experienced by cells, demonstrating a strong correlation with the nuclear volume, which is in good agreement with the piezoelectric function of the cell nucleus predicted by previous theoretical studies [25]. This may provide a potential mechanism by which cells transduce mechanical forces they experience when spreading on a substrate or migrating though a narrow space into other types of physicochemical and intracellular signals.

Indeed, mirochannel cell-migration experiments showed that the nuclear-to-cytoplasmic ratio (N/C ratio) of the mEGFP WT probe strongly correlates with the mean nuclear electrostatic potential and nuclear volume, see Figure S8(c,d), SI. By multiplying the reciprocal of the slope of the fitting curve shown in Figure S8(c), SI, by the net electrical charge of mEGFP WT (*q* = − 7.3*e*) and dividing by *k*_B_*T*, it can be found that the interaction of mEGFP WT with the nuclear electrostatic potential contributes ∼ 60% to its N/C ratio in MDA-MB-231 cells, with the remaining ∼ 40% due to the volume-exclusion effect.

For comparison, similar measurements performed for the mEGFP+3RK probe, which carries a nearly neutral net electrical charge, showed that its N/C ratio was higher than that of mEGFP WT under all experimental conditions, see Figure S8, SI. This indicates that the C-terminal 3RK peptide of this construct is likely recognized by importins involved in active nuclear transport, as it has high homology to the monopartite nuclear localization signal [40–42]. Thus, the mechanosensitivity of the N/C ratio of mEGFP+3RK observed in our experiments reflects the recently described mechanosensitive behaviour of active nuclear transport [43]. Notably, a direct comparison of the slopes of the N/C ratio curves versus nuclear volume for the mEGFP WT and EGFP+3RK probes shown in Figure S8(d,e), SI, demonstrates that the contribution of electrostatic and volume-exclusion interactions to the mechanosensitive behaviour of mEGFP WT is comparable (albeit smaller) to the contribution of active nuclear transport to the mechanosensitive behaviour of mEGFP+3RK. This result suggests that nuclear electrostatic potential may function alongside active nuclear transport and volume-exclusion interactions in regulating the N/C ratio of proteins and other macromolecules, particularly those carrying a high electrical charge.

#### Effect of spatial heterogeneity of nuclear electrostatic potential on the nucleoplasmic distribution of histone-chaperone complexes

In addition to the mechanosensitive behaviour of the nuclear electrostatic potential, the above experiments also showed that it has a highly non-uniform distribution throughout the nucleus [Figure 3(b,d)], indicating that phase-separation processes, such as the formation of euchromatic and heterochromatic regions, play an important role in shaping the nuclear electrostatic potential landscape, see Appendix A.

Although the average magnitude and spatial heterogeneity of the nuclear electrostatic potential are only on the order of ∼ 1 mV [Figure 3], such an electrical field may still have a potentially significant effect on the nucleoplasmic distribution of highly electrically charged biomolecules, such as long-noncoding RNAs or freely diffusing histones, as well as on the stability of nucleosomes. For example, the electrical charge of H2A · H2B and H3 · H4 histone dimers is on the order of +37*e*, so the average difference in nuclear electrostatic potential between euchromatic and heterochromatic regions measured in the experiments (0.617 ± 0.026 mV, mean ± SEM, *N* = 21 cells) would result in a ∼ 0.85 k_B_T difference (1 k_B_T ≈ 26.7 *e* · mV) in the free energy of histone dimers located in the corresponding regions, which could affect the nucleoplasmic distribution of freely diffusing histones and the stability of nucleosomes (by ∼ 3.4 k_B_T). In fact, as shown below, this difference in the nucleosome stability between euchromatic and heterochromatic regions appears to be even higher [∼ 7.1 k_B_T, see Figure 5(b) and Appendix C]. Thus, although the experimentally measured spatial heterogeneity of the nuclear electrostatic potential may have little effect on monovalent ions and electrically charged cellular metabolites (the free energy change is *<* 0.04 k_B_T), it may significantly affect the behaviour of protein complexes carrying a strong electrical charge.

To experimentally test the influence of spatial heterogeneity of the nuclear electrostatic potential on the nucleoplasmic distribution of freely diffusing histone dimers, we performed FRAP experiments in HEK293T cells grown on glass slides that coexpressed mEGFP WT probes and H2B-mCherry or H3.1-mCherry histones. In the experiment, a fairly large area of the nucleus, encompassing both euchromatic and heterochromatic regions, was photobleached with a laser, and the recovery of the histone fluorescent signal was measured as a function of time in euchromatic and heterochromatic regions and correlated with the nuclear electrostatic potential difference measured between these regions, see Figure 4 and Figure S9, SI.

**FIG. 4.**
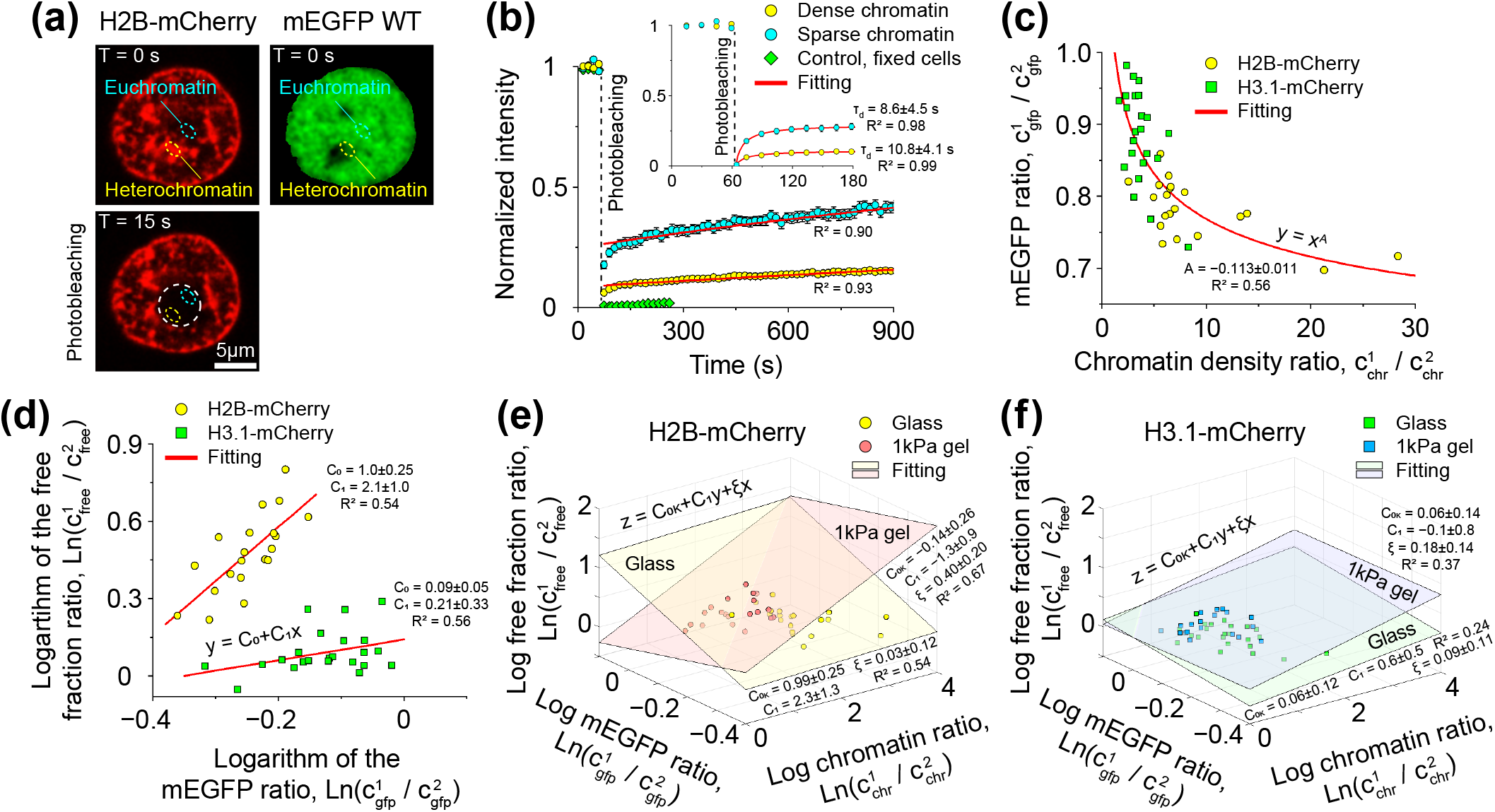
Effect of chromatin on the nucleoplasmic distribution of histone-chaperone complexes. **(a)** Photobleaching of fluorescently labelled histones in HEK293T cells coexpressing H2B-mCherry and mEGFP WT. The H2B-mCherry FRAP signal was measured in euchromatic (sparse) and heterochromatic (dense) regions corresponding to low and high nuclear electrostatic potential (i.e., low and high mEGFP signal). The white dashed circle indicates the photobleached area. Cells were grown on fibronectin-coated glass slides. **(b)** H2B-mCherry FRAP signals measured in dense (yellow data points) and sparse (cyan data points) chromatin regions of HEK293T living cells. Data fitting was performed as described in Appendix E, resulting in an estimate of the average densities of freely diffusing H2A H2B histone-chaperone complexes and chromatin in the corresponding regions. A similar analysis was also performed in HEK293T cells coexpressing H3.1-mCherry histones and mEGFP WT, see Figure S9, SI. The graph shows the average over *N* = 21 live cells (cyan and yellow data points) and, for comparison, *N* = 8 fixed HEK293T cells (green data points). Error bars indicate SEM. The inset displays an enlarged region of the graph before and immediately after the photobleaching. **(c)** Correlation between the mEGFP signal ratio 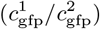 measured for dense (1) and sparse (2) chromatin regions and chromatin density ratio 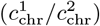 in the same regions estimated from the fitting of the H2B-mCherry and H3.1-mCherry FRAP signals shown in panel (b) and Figure S9, SI. Each data point represents a single FRAP experiment. The red curve shows the data fitting to the power function. **(d-f)** Correlation between the logarithm of the mEGFP signal ratio 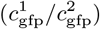 measured for dense (1) and sparse (2) chromatin regions and the logarithms of the chromatin density ratio 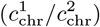 and the density ratio of freely diffusing histone-chaperone complexes 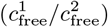 in the same regions, see also Figure S10, SI. The slopes of the fitting functions determine the strength of volume-exclusion and electrostatic interactions between histone-chaperone complexes and the surrounding microenvironment, see Appendix F for details. Non-zero intercepts of the fitting functions with the y-axis [panels (d, e)] indicate that histone-chaperone complexes may engage in weak transient binding interactions with the surrounding microenvironment. Fitting parameter values in panels (b-f) are presented as mean *±* 95% CI.

**FIG. 5.**
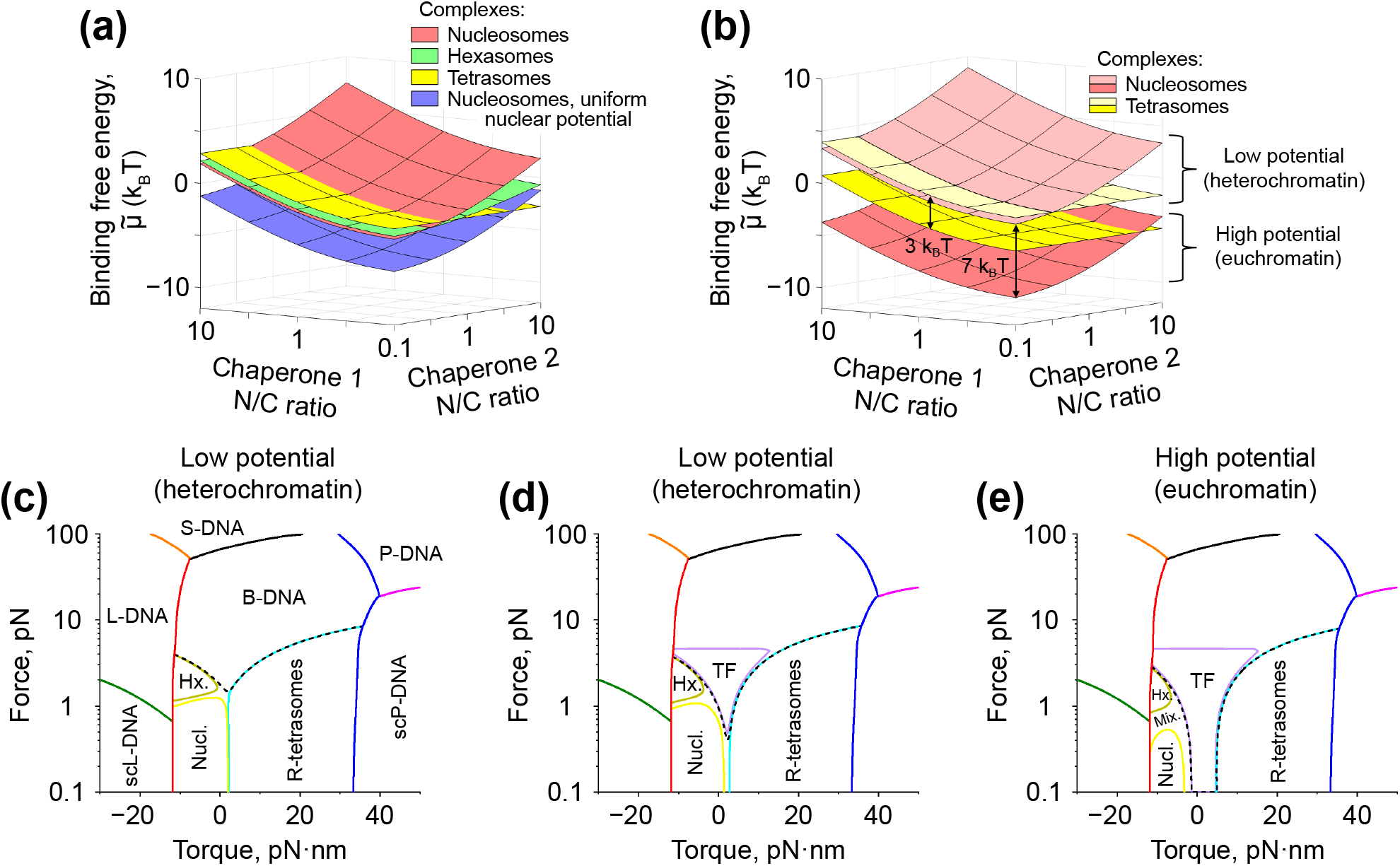
Effect of nuclear electrostatic potential on the stability of histone-based nucleoprotein complexes. **(a)** Combined contribution of the nuclear electrostatic potential, weak transient interactions and active nuclear transport to the binding free energy of histone octamers, hexamers and tetramers to DNA as a function of the nuclear-to-cytoplasmic (N/C) ratio of chaperones involved in the transportation of H2A *·* H2B and H3 *·* H4 histone dimers (chaperones 1 and 2, respectively), see Appendix G for details. On average, nucleosomes are more strongly stabilized by electrostatic interactions compared to hexasomes and tetrasomes. Furthermore, comparison of the results obtained for histone octamers (red) to the calculations performed for a hypothetical case of a uniform nuclear electrostatic potential (purple) of the same strength shows that heterogeneity of the nuclear electrostatic potential may play a significant role in nucleosome stabilization. **(b)** Sensitivity of the binding free energy of histone octamers and tetramers to DNA to heterogeneity of nuclear electrostatic potential. The difference between the binding free energies of histone octamers to DNA in regions of high and low nuclear electrostatic potential, which are often associated with euchromatin and heterochromatin, respectively, is approximately twice that of histone tetramers. As a result, tetrasomes are likely to be more abundant in euchromatin compared to heterochromatin. **(c)** Phase diagram of DNA located in a region of low electrostatic potential (heterochromatin) in the presence of histone-based nucleoprotein complexes [for euchromatin, see Figure S18(b), SI]. The solid curves represent the transition boundaries between extended (B, L and P) and supercoiled (scL and scP) states of DNA, as well as between different DNA-protein conformations, such as: right-handed tetrasomes (R-tetrasomes), hexasomes (Hx.), nucleosomes (Nucl.) or a mixture of the last two with L-tetrasomes (Mix.), see Appendix H. The dashed curve indicates the boundary of DNA state in which *>* 50% of DNA is occupied by histone-based nucleoprotein complexes. **(d, e)** Phase diagrams for DNA located in regions of low (d) and high (e) electrostatic potential in the presence of DNA-binding competition between histones and DNA-bending transcription factors (TFs). Low electrostatic potential of heterochromatic domains stabilizes nucleosomes and tetrasomes, preventing strong binding of TFs to DNA at low forces (*≥* 0.5 pN), whereas euchromatic domains with high electrostatic potential are unable to resist it. In the calculations, the binding free energy of the TF to DNA was 2.3 k_B_T, which corresponds to the experimentally measured concentration of TFs *∼* 10 times higher than their equilibrium dissociation constant from DNA [219, 220]. The values of the rest of the model parameters can be found in Tables T1-T3, SI.

It was found that the histone fluorescent signal in both euchromatic and heterochromatic regions recovers in two stages: an initial rapid partial recovery within ∼ 60 s after photobleaching is typically followed by a slower recovery over a longer period of time, see Figure 4(b) and Figure S9, SI. These results suggest that there are likely two distinct molecular processes that contribute to the recovery of the histone fluorescent signal: 1) by diffusion of a freely moving fraction of histones into the photobleached area, and 2) by exchange of fluorescently-labelled histones between histone-chaperone complexes and nucleoprotein complexes formed on DNA. Indeed, FRAP experiments in HEK293T cells coexpressing H2B-mCherry histones with the main chaperones involved in their transport, Nap1L1 or Nap1L4 (conjugated to EGFP) [4], revealed that the initial recovery of the histone signal was accompanied by a rapid recovery of the chaperone fluorescent signal, indicating that it is indeed mainly due to the diffusion of histone-chaperone complexes into the photobleached region, see Figure S11(a,b), SI. Over longer periods of time, the fluorescent signal of chaperones tended to flatten out, often reaching a plateau, suggesting that the recovery of the histone signal on this time scale is likely due in part to the exchange of fluorescently-labelled histones between histone-chaperone complexes and nucleoprotein complexes formed on DNA, rather than diffusion of histone-chaperone complexes into the photobleached region. Similar results were also obtained in HEK293T cells coexpressing H3.1-mCherry histones and one of their main chaperones, ASF1a (conjugated to EGFP) [4], see Figure S11(c), SI.

Interestingly, FRAP experiments showed that the characteristic diffusion time of chaperones into the photo-bleached region, *τ*_d_ (*τ*_d_ = 6 − 11 s), is largely independent of local chromatin density, suggesting that the volume-exclusion effect does not significantly influence chaperone diffusion, see Figure S11, SI. More importantly, these experiments demonstrated that *τ*_d_ is more than two orders of magnitude shorter than the characteristic histone turnover time in nucleoprotein complexes (*>* 1000 s), see Figure 4(b) and Figure S9, SI. Thus, these two processes are well separated in time, allowing experimental determination of the local density of the freely diffusing histone fraction by measuring the histone signal approximately ∼ 60 s after photobleaching.

FRAP experiments showed that the ratio of mEGFP WT fluorescent signals measured in photobleached euchromatic and heterochromatic regions has a strong non-linear correlation with the heterochromatin / euchromatin density ratio [Figure 4(c)], highlighting the fact that in HEK293T cells grown on glass, the nuclear electrostatic potential is mainly determined by the local chromatin density. Furthermore, the rapid recovery of the histone fluorescent signal in the euchromatic and heterochromatic regions 60 s after photobleaching, due to diffusion of the freely moving fraction of histones into the photobleached area, appeared to correlate with the strength of the local nuclear electrostatic potential measured with mEGFP WT probes. In particular, the ratio between the initial histone fluorescence recovery signals measured in euchromatic and heterochromatic regions showed a significant correlation with the ratio of mEGFP WT signals measured in the same regions in the case of H2A · H2B histone dimers and a much smaller correlation in the case of H3 · H4 histone dimers [Figure 4(d)], indicating that the nucleoplasmic distribution of freely diffusing histone dimers transported by chaperones is sensitive to the local nuclear electrostatic potential in the case of H2A · H2B histone dimers and to a lesser extent in the case of H3 · H4 histone dimers.

Furthermore, estimates based on the slopes (parameter *C*_1_) of the plots presented in Figure 4(d-f) showed that the effective net electrical charge of the chaperone complexes transporting fluorescently-labelled H2A·H2B histone dimers is of the order of −19*e* ± 6*e*, whereas for H3·H4 histone-chaperone complexes it is +3*e* ± 3*e* see Appendix F for details. Although the calculated charges include the net electrical charge of mChrerry (*q* = − 6*e*) used in the experiments to label histones, this result suggests that the chaperone complexes largely shield the positive charges of the histone dimers (*q* = +37*e*), which is in good agreement with previous experimental studies showing that histone-binding chaperones tend to have a strong negative charge [44–46].

In addition, it can be seen from Figure 4(d-f) that the linear fitting functions have a non-zero intercept with the y-axis (*C*_0K_ = 0.99 ± 0.25 in the case of H2A·H2B histone dimers and *C*_0K_ = 0.06 ± 0.12 in the case of H3 · H4 histone dimers, mean ± 95% CI), suggesting that in addition to electrostatic interactions, freely diffusing histones transported by chaperones also engage in other types of weak attractive / binding interactions with the surrounding microenvironment in the nucleus, which are stronger for H2A · H2B histone dimers compared to H3 · H4 histone dimers, see details in Appendix F. In particular, the Y-intercept value for H2A · H2B histone dimers indicates that the equilibrium dissociation constant of H2A·H2B histone-chaperone complexes from such weak binding sites in sparse chromatin regions is 2.7-fold higher than that in dense chromatin regions, suggesting that freely diffusing H2A · H2B histone-chaperone complexes bind more strongly to sites located in dense chromatin regions. Thus, weak binding sites exert an effect on histone-chaperone complexes that is opposite to the effect of nuclear electrostatic potential and volume-exclusion interactions, attracting these complexes to dense regions of chromatin. Interestingly, fitting of the experimental data revealed that the density of such binding sites exhibits negative cooperativity [*ξ* = 0.03 − 0.4, Figure 4(e,f)] depending on the local chromatin density, suggesting that these sites could potentially be formed by nuclear components other than chromatin itself.

In contrast to HEK293T cells grown on glass slides, H2A·H2B histone-chaperone complexes exhibited a completely different behaviour in HEK293T cells grown on 1 kPa gel, indicating a possible change in the magnitude of their effective net electrical charge, accompanied by a decrease in the strength of their weak binding interactions with the surrounding microenvironment, see Figure S10, SI. This result demonstrates that substrate elasticity may affect not only the nuclear electrostatic potential but also the function of histone-chaperone complexes.

Overall, FRAP experiments suggest that nuclear electrostatic potential and weak transient binding interactions of histone-chaperone complexes with the surrounding microenvironment may play an important role in determining their nucleoplasmic distribution throughout the nucleus.

#### Effect of nuclear electrostatic potential on nucleosome stability

The stability of nucleoprotein complexes, such as tetrasomes, hexasomes and nucleosomes, is generally determined by the binding free energy of the corresponding protein complexes (histone tetramers, hexamers and octamers) to DNA, see Appendix G. Experimental measurements of the nucleoplasmic distribution of chromatin and the nuclear electrostatic potential allow one to estimate the effect of volume-exclusion and electrostatic interactions on the binding free energy of histones to DNA, which makes it possible to assess the influence of such interactions on the stability of the corresponding nucleoprotein complexes, see Appendices C, G.

As a result, it was found that nuclear electrostatic potential exerts a strong stabilizing effect on nucleosomes, enhancing the binding of histone octamers to DNA up to 7 k_B_T in the case of HEK293T cells grown on fibronectin-coated glass slides, see Figure 5(a). On the other hand, hexasomes are stabilized by it to a lesser extent (up to 4.5 k_B_T), while the stability of tetrasomes changes even less (up to 3 k_B_T). Interestingly, similar estimates obtained from experimental data collected on HEK293T cells grown on 1 kPa gel showed that histone-based nucleoprotein complexes should be stabilized by the nuclear electrostatic potential to a lesser extent compared to the case of HEK293T cells grown on glass slides [Figure S17(a), SI], suggesting that the stabilization effect may be sensitive to the elastic properties of the substrate on which cells are grown, which is in good agreement with previous theoretical studies [25].

More importantly, it was found that spatial heterogeneity of the nuclear electrostatic potential may play an important role in the stabilization of histone-based nucleoprotein complexes, as they demonstrated that in the hypothetical case of a uniform nuclear electrostatic potential, the binding free energy of histone octamers to DNA should not change significantly, see red and purple 2D surfaces in Figure 5(a). I.e., the spatial heterogeneity of the nuclear electrostatic potential leads to the emergence of ‘electrostatic traps’, which, in the case of cells grown on glass slides, are created by regions of high chromatin density (heterochromatin), significantly stabilizing nucleosomes localized in them.

To further demonstrate this point, we estimated, based on experimental data, the combined contribution of the nuclear electrostatic potential, weak binding interactions between histone-chaperone complexes and the nuclear microenvironment, and active nuclear transport to the binding free energy of histone-based protein complexes located in regions of low (heterochromatin) and high (euchromatin) nuclear electrostatic potential, see Figure 5(b). The calculations showed that while histone-based nucleoprotein complexes are stabilized in heterochromatic regions, they experience the opposite destabilizing effect in euchromatic regions, see Figure 5(b). This destabilization appears to result from the fact that the interaction energy between the histone-based nucleoprotein complexes and the local nuclear electrostatic potential in euchromatic regions is not large enough to compensate for the change in electrochemical potentials of the protein complexes that participate in nucleosome assembly (Figure S16, SI).

The difference in the binding free energies of histone octamers to DNA in euchromatic and heterochromatic regions, which is mainly due to the electrostatic interaction between histone octamers and the local nuclear electrostatic potential, was found to be of the order of ∼ 7 k_B_T. On the other hand, histone tetramers were found to be less affected by the local nuclear electrostatic potential, with the difference between the binding free energies of histone tetramers to DNA in euchromatic and heterochromatic domains being ∼ 3 k_B_T [Figure 5(b)]. Thus, euchromatic and heterochromatic regions may exert very different stabilizing effects on nucleosomes and tetrasomes via the nuclear electrostatic potential.

This may have a significant influence on the DNA-binding competition between histones and other proteins, such as TFs. Indeed, previous theoretical studies show that even fairly small changes of a few k_B_T in the binding free energy of different types of proteins competing for DNA binding can significantly shift the balance between them in terms of the fraction of DNA occupied by these proteins [29]. In particular, the above results suggest that euchromatic regions should be generally more accessible to TFs binding and thus are likely associated with transcriptionally active regions of chromatin, which is in good agreement with existing studies [6, 13].

Previous single-molecule studies suggest that one of the most direct experimental ways to quantify the stability of tetrasomes and nucleosomes is to apply a mechanical load to DNA and measure the characteristic force that causes dissociation of such complexes [47–50]. Indeed, the mere formation of nucleoprotein complexes on DNA reveals little about their stability. However, by subjecting them to destabilizing stimuli, such as stretching force, their stability can be directly quantified.

To compare the binding free energy estimates obtained in our study to previously published single-molecule experimental measurements, we employed a recently developed transfer-matrix framework that not only allows for an accurate description of the transition boundaries between different states of DNA depending on the applied mechanical constraints [51, 52], but can also be used to assess the mechanical stability of nucleoprotein complexes [25, 53].

Stability diagrams calculated using the transfer-matrix method for heterochromatin and euchromatin based on the estimates of the binding free energies of histone-based nucleoprotein complexes [see Appendix G and Table T3, SI, for details] showed that, while the characteristic stretching force leading to the destabilization of such complexes in heterochromatin at low torques is ∼ 1.5 − 2 pN, for euchromatin it is ∼ 0.8 − 1.2 pN, see Figure 5(c) and Figure S18, SI. These values are in very good agreement with the characteristic chromatin unfolding force measured in previous single-molecule studies in *Xenopus* egg extracts (∼ 1.2 − 1.7 pN in the presence of ATP [48]), indicating that transfer-matrix calculations accurately describe the stability of histone-based nucleoprotein complexes and that the binding free energy estimates obtained in our study are consistent with those obtained previously in single-molecule studies.

Notably, the above results indicate that histone-based nucleoprotein complexes are approximately two-fold more stable in heterochromatin compared to euchromatin, suggesting that they may behave differently in competing with TFs for DNA binding. Indeed, stability diagrams calculated for DNA in the presence of DNA-binding competition between histones and TFs showed that histone-based nucleoprotein complexes located in heterochromatic regions are able to almost completely prevent the binding of TFs to DNA when no force or a small force (*f* ≤ 0.5 pN) is applied to it, see Figure 5(d). On the other hand, the destabilizing effect of euchromatin on histone-based nucleoprotein complexes makes it difficult for them to efficiently compete with TFs, allowing the latter to bind to DNA when the torque (*τ*) applied to it is small enough (− 1.5 pN · nm ≥ *τ* ≥ 5 pN · nm), see Figure 5(e).

Interestingly, Figure 5(d) also shows that by destabilizing nucleosomes and tetrasomes, a stretching force of the order of a few piconewtons applied to DNA can shift the DNA-binding competition between histones and TFs in favour of the latter, resulting in TFs binding to DNA under conditions similar to those found in heterochromatin, potentially explaining the force-induced activation of gene transcription previously observed in experimental cell studies [54]. Indeed, previous single-molecule experiments have shown that DNA-binding competition between histones and gene transcription repressors can indeed be regulated by mechanical constraints applied to DNA and potentially exploited by archaeal cells to regulate gene transcription [55]. Thus, it can be concluded that the local nuclear electrostatic potential, as well as mechanical constraints applied to DNA, may have a significant impact on the DNA-binding competition between different types of histone-based nucleoprotein complexes and TFs, differentially regulating DNA accessibility to such proteins in euchromatic and heterochromatic regions.

Taken together, the above results indicate that nucleosomes located in heterochromatin should be more stable than nucleosomes located in euchromatin. Furthermore, the smaller difference in the binding free energy of tetrasomes between heterochromatic and euchromatic regions [3 k_B_T, Figure 5(b)] compared to nucleosomes [7 k_B_T, Figure 5(b)] suggests that the fraction of DNA occupied by tetrasomes should generally be higher in euchromatin compared to heterochromatin. Thus, it can be expected that H2A · H2B histone dimers, which are incorporated into nucleosomes at the final stages of their assembly (Figure S16, SI), should potentially exhibit very different behaviour in heterochromatic and euchromatic regions. Indeed, previous experimental studies have shown that H2A · H2B histone dimers, but not H3 · H4 histone dimers, have increased mobility in transcriptionally active regions [56–58] that are typically associated with euchromatin, which is consistent with the above predictions.

Furthermore, experimental measurements of the fluorescent signal ratio of H3.1-mCherry and H2B-EGFP coexpressed in HEK293T cells conducted in our study showed that it significantly correlated with local chromatin density, – the H3/H2B fluorescent signal ratio was found to be approximately two-fold higher in sparse (euchromatic) regions compared to dense (heterochromatic) regions, see Figure 6(a). On one hand, this result may indicate that nucleosomes are generally less stable in euchromatic regions compared to heterochromatic regions, leading to a higher frequency of partially assembled histone-based nucleoprotein complexes, such as tetrasomes and hexasomes. On the other hand, it may also be due to the tendency of H3.1 histones to localize to transcriptionally silent regions of chromatin, including heterochromatin, which contrasts with the behaviour of H3.3 histones, which localize to transcriptionally active euchromatic regions and telomeres [59–63].

**FIG. 6.**
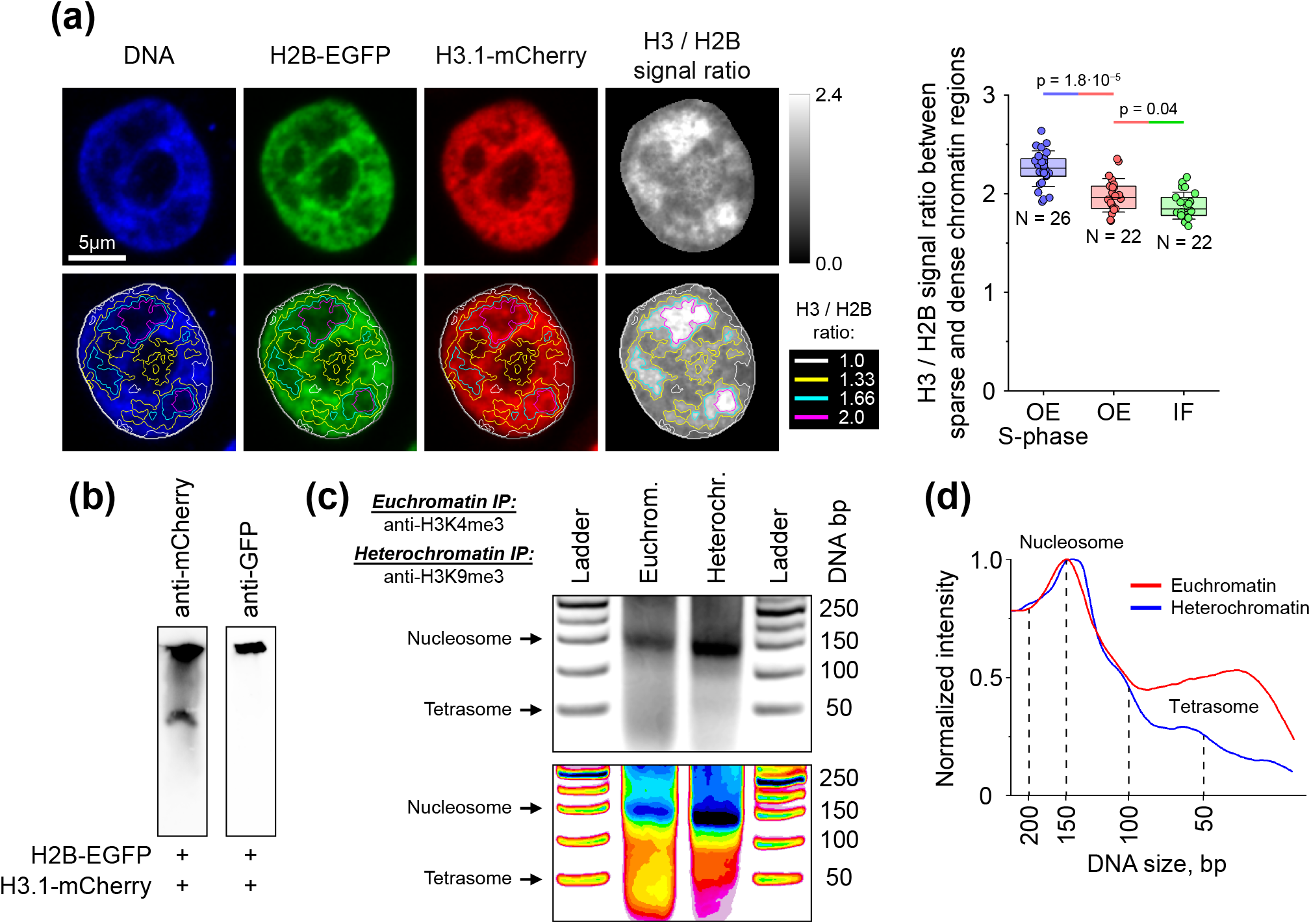
Relative distribution of H2B-EGFP and H3.1-mCherry histones in the nucleus. **(a)** Experimentally measured H2B-EGFP and H3.1-mCherry fluorescence signals and their ratio (*R* = *I*_H3_*/I*_H2B_) in HEK293T cells. As can be seen from the boxplot on the right, the ratio of H3.1-mCherry and H2B-EGFP fluorescent signals is approximately twice as high in sparse chromatin regions compared to dense chromatin regions (*R*_sparse_*/R*_dense_ = 1.98 *±* 0.04, mean *±* SEM, see OE data). Very similar results were also obtained in cells arrested in the S-phase of the cell cycle (OE S-phase data, *R*_sparse_*/R*_dense_ = 2.25 *±* 0.04, mean *±* SEM) and in wild-type HEK293T cells immunostained with antibodies to H2B and H3 histones (IF data, *R*_sparse_*/R*_dense_ = 1.88 *±* 0.03, mean *±* SEM). These results indicate that in sparse chromatin regions, chromatin may contain a high percentage of hexasomes / tetrasomes, whereas in dense chromatin regions it is composed mainly of nucleosomes. In total, *N* = 26, 22 and 22 cells were measured in the respective cases. In the graph, boxes represent the interquartile range, error bars indicate SD. The bar above the graph shows the pairwise statistical difference between the corresponding experimental data, calculated using Mann-Whitney test. **(b)** Western blot of a native PAGE gel of MNase-digested chromatin isolated from HEK293T cells coexpressing H3.1-mCherry and H2B-EGFP, stained with antibodies to mCherry and GFP. The image shows that the digested chromatin contains tetrasomes (lower band, visible only when stained with antibodies to mCherry) and nucleosomes (upper band, visible when stained with antibodies to mCherry or GFP). **(c)** DNA gel of MNase-digested chromatin immunoprecipitated from HEK293T cells using antibodies to H3K4me3 and H3K9me3 histones, which are typically found in euchromatin and heterochromatin, respectively. The top panel shows the original DNA gel, the bottom image – the same gel in a 16-color scale for improved contrast. **(d)** Intensity profiles of euchromatin and heterochromatin lanes from panel (c) normalized to the nucleosome peak. It can be seen that tetrasomes are enriched in MNase-digested euchromatin compared to heterochromatin, indicating a stronger destabilizing effect of euchromatin on nucleosomes than on tetrasomes, which is consistent with the model predictions shown in Figure 5.

To exclude the second scenario, we performed an additional experiment of co-immunostaining permeabilized HEK293T cells with antibodies to H2B and H3 histones, which, among other things, recognize both H3.1 and H3.3 histones. This experiment showed that, as in the case of live HEK293T cells coexpressing H2B-EGFP and H3.1-mCherry, there is a significant correlation between the ratio of H2B and H3 fluorescent signals and the spatial heterogeneity of chromatin [Figure 6(a)], suggesting that this phenomenon is not related to the histone H3.1 isoform used in the live HEK293T cell experiments. In addition, FRAP experiments showed that freely diffusing histone-chaperone complexes were absent from the nuclei of permeabilized cells and therefore did not contribute to the measured fluorescent signal ratio [Figure 4(b)].

To further test the potential contribution of the freely diffusing fraction of histones to the H3/H2B fluorescent signal ratio, we arrested HEK293T cells coexpressing H3.1-mCherry and H2B-EGFP in the S-phase of the cell cycle by treating them with hydroxyurea for 24 h. Indeed, expression levels of endogenous core histones are known to increase in the S-phase [64], and by competing with fluorescently-labelled histones for DNA binding, they force the latter to shift more towards the freely diffusing fraction. Direct measurements performed in FRAP experiments confirmed this, showing that HEK293T cells arrested in the S-phase have a higher fraction of freely diffusing fluorescently-labelled histones compared to untreated cells, see Figure 4(b) and Figures S9, S12(a,b), SI. Although the level of freely diffusing H2B-EGFP histones increased only modestly in these experiments, the fraction of freely diffusing H3.1-mCherry histones increased approximately two-fold, – from ∼ 0.2 − 0.4 [Figure S9, SI] to ∼ 0.5 − 0.7 [Figure S12(b), SI]. Yet, this large change in the fraction of freely diffusing H3.1-mCherry histones resulted in only a small increase in the H3/H2B fluorescent signal ratio between sparse and dense chromatin regions by ∼14%, as shown in Figure 6(a), suggesting that this is a very robust observable parameter primarily determined by DNA-bound nucleoprotein complexes.

To test whether H3.1-mCherry and H2B-EGFP histones were indeed incorporated into nucleosomes and tetrasomes (in the case of H3.1-mCherry) in the above experiments, we isolated chromatin from HEK293T cells and digested it with MNase. Western blotting of the native PAGE gel of MNase digestion products using antibodies to mCherry and GFP revealed the presence of two types of nucleoprotein complexes: 1) containing only H3.1-mCherry histones, and 2) containing both H3.1-mCherry and H2B-EGFP histones, which likely correspond to tetrasomes and nucleosomes, respectively [Figure 6(b)], indicating efficient incorporation of H3.1-mCherry and H2B-EGFP histones into nucleoprotein complexes.

To confirm that the above complexes are indeed tetrasomes and nucleosomes, and to investigate whether tetrasomes are enriched in the euchromatic or heterochromatic fractions of chromatin isolated from HEK293T cells, we performed immunoprecipitation of euchromatin and heterochromatin fragments using antibodies to H3K4me3 and H3K9me3 histones, which are typically found in euchromatin and heterochromatin, respectively, followed by MNase digestion and DNA gel electrophoresis. As a result, it was revealed that in the heterochromatin there was only one band in the range of 0 − 250 bp, corresponding to nucleosomes (∼ 150 bp). On the other hand, in the euchromatin sample, in addition to the presence of the same band corresponding to nucleosomes, an enrichment of DNA fragments of ∼ 50 bp in size, approximately corresponding to the size of tetrasomes, was also observed, see Figure 6(c,d). These results suggest that the fraction of tetrasomes is higher in isolated euchromatin compared to isolated heterochromatin, indicating a destabilizing effect of euchromatin on nucleosomes, which is consistent with the binding free energy estimates presented in Figure 5(b).

Overall, the above results imply that the enrichment of heterochromatic regions in H2A · H2B histone dimers is likely due to the difference in the stability of nucleosomes located in heterochromatic and euchromatic regions, indicating the possible importance of local chromatin density and hence local nuclear electrostatic potential in regulating nucleosome stability.

## DISCUSSION

In this study, we developed an experimental method for mapping the nuclear electrostatic potential and the strength of volume-exclusion interactions in the nuclei of living cells. Using this method, it was found that the mean nuclear electrostatic potential is sensitive to the expression level of supercharged nuclear proteins, such as H1 linker histones (Figure S7), and the size of the cell nucleus (Figure 3), which is in good agreement with previous theoretical predictions [25]. Moreover, the nuclear electrostatic potential was found to exhibit strong spatial heterogeneity, which in the case of cells grown on rigid substrates appears to correlate with the nucleoplasmic distribution of chromosomal DNA / chromatin, see Figure 2. On the other hand, in cells grown on soft 1 kPa gel, the nuclear electrostatic potential, although remaining quite non-uniform, showed no correlation with DNA / chromatin density, indicating that in this case it is probably determined by other electrically charged components of the nucleus. Since previous studies have shown that the Young’s modulus of various living tissues varies in the range from 200 Pa to 100 kPa [65], these experiments suggest that both of these mechanisms could potentially have physiological significance in various living tissues.

Furthermore, our experiments demonstrated that the nucleoplasmic distribution of electrically charged proteins and their complexes, such as freely diffusing histone-chaperone complexes, correlates with the nuclear electrostatic potential (Figure 4), suggesting that the stability of nucleoprotein complexes assembled from histones, such as tetrasomes, hexasomes and nucleosomes, may depend on the spatial heterogeneity of the nuclear electrostatic potential. Indeed, theoretical estimates based on experimentally measured nucleoplasmic distributions of chromatin and nuclear electrostatic potential have shown that spatial heterogeneity of nuclear electrostatic potential may lead to a ∼ 7 k_B_T difference in the binding free energy of histone octamers to DNA in euchromatic regions compared to heterochromatic regions [Figure 5(b)]. This roughly corresponds to the standard Gibbs free energy of nucleosome assembly under normal conditions in *in vitro* bulk solution experiments (∼ 7 k_B_T [66]), indicating that the nuclear electrostatic potential may significantly influence nucleosome stability in living cells. Subsequent measurements of the H3/H2B fluorescent signal ratio in HEK293T cells coexpressing H3.1-mCherry and H2B-EGFP constructs, as well as MNase chromatin digestion experiments, confirmed these estimates, showing that nucleosome stability correlates with local chromatin density and, consequently, with nuclear electrostatic potential, see Figure 6. Overall, these results suggest that nuclear electrostatic potential may be one of the key factors influencing nucleosome stability in response to the surrounding chromatin microenvironment.

Further theoretical calculations showed that such strong stabilization of nucleosomes by heterochromatin leads to the fact that nucleosomes and tetrasomes, like Scylla and Charybdis from Greek mythology, jointly block a narrow region on the DNA stability diagram corresponding to the state of DNA bound by TFs, see Figure 5(d,e). This Scylla-Charybdis mechanism (or SC-mechanism for short) may provide one of the previously missing links to explain the heterochromatin-mediated gene transcriptional silencing effect [23]. In contrast to heterochromatin, similar calculations performed for euchromatic regions showed that due to the lower stability of nucleosomes in euchromatic regions, the SC-mechanism losses its efficiency and becomes unable to provide significant resistance to DNA binding by TFs, suggesting that genes located in euchromatic regions are more accessible to the gene transcription machinery.

Based on the obtained results, it is tempting to speculate a possible role of some post-translational modifications (PTMs) of histones in nucleosomes in the regulation of gene transcription. In particular, previous experimental studies indicate that many specific histone PTMs are associated with different transcriptional states of chromatin. For example, lysine acetylation on the tails of histones H3 and H4 is usually associated with transcriptionally active chromatin, whereas trimethylation of lysine 9 and 27 of histones H3 (H3K9me3, H3K27me3) is a typical feature of heterochromatin, which is associated with a repressed transcriptional state [67]. It has previously been shown that one of the main proteins involved in the formation of heterochromatin, HP1, is capable of recognizing and binding to H3K9me3 histones, forming cross-links between a pair of nucleosomes containing such histones [68–70]. However, the experimentally measured average residence time of HP1 protein on chromatin was found to be on the order of seconds [71], suggesting that HP1-mediated cross-linking between nucleosomes is highly dynamic and unlikely to directly contribute to nucleosome stabilization. On the other hand, it has been proposed that these dynamic HP1-mediated cross-links lead to chromatin condensation and heterochromatin formation via a liquid phase-separation process [7, 8]. According to our results, such chromatin condensation may result in a significant change in the local nuclear electrostatic potential, followed by nucleosome stabilization, rendering heterochromatin into a closed state via the SC-mechanism and making it less accessible to TFs. Thus, one of the potential roles of heterochromatin-associated histone PTMs in repressing the transcriptional state of chromatin may be the formation of regions of low electrostatic potential (i.e., ‘electrostatic traps’) due to chromatin condensation that trap nucleosomes in a more stable state.

In addition to HP1, it has been suggested that linker histone H1 may also be involved in nucleosome stabilization and chromatin condensation by cross-linking incoming and outgoing DNA in nucleosomes [72–76]. Although the exact role of the linker histone H1 *in vivo* is not fully understood, it has been found to accumulate in heterochromatic regions [77] and is generally associated with a transcriptionally silent chromatin state [78]. Surprisingly, *in vitro* bulk experimental studies have shown that the standard Gibbs free energy of linker histone H1 binding to nucleosomes is only ∼ 1 k_B_T [66], indicating its weak affinity for nucleosomes. On the other hand, the high electrical charge of H1 (+53.0*e*) suggests that, like the core histones H2A, H2B, H3 and H4, it may be very sensitive to the local nuclear electrostatic potential. Indeed, estimates based on experimental measurements of the nuclear electrostatic potential carried out in our study indicate that when histone H1 binds to nucleosomes located in heterochromatic regions, it may stabilize them by an additional ∼ 1 − 2 k_B_T compared to the case of H1 binding to nucleosomes in euchromatic regions. Thus, linker histone H1, working in conjunction with HP1 proteins, could potentially be used by cells to further fine-tune nucleosome stability and chromatin condensation in different parts of the nucleus.

What about histone PTMs typically found in euchromatin, such as H3K9ac? Acetylation of the lysine 9 of histone H3 (H3K9ac) requires its prior demethylation, which reduces the affinity of HP1 proteins for nucleosomes [69]. The reduced affinity of acetylated nucleosomes for the heterochromatin-associated protein HP1, coupled with the stronger volume-exclusion effect and electrostatic repulsion experienced by nucleosomes in heterochromatin compared to euchromatin (∼ 4 *k*_B_*T* energy difference, Appendix C), may ultimately result in the displacement of acetylated nucleosomes from heterochromatic regions due to local chromatin mobility [1] to less dense adjacent euchromatic regions with a more positive electrostatic potential. According to our findings, this should lead to a more open chromatin state because of the weakening of the SC-mechanism, which allows the recruitment of TFs and increases the accessibility of DNA to transcription factories present in the interchromatin space / euchromatic regions [6, 13]. In addition to the processes mentioned above, previous studies have shown that histone PTMs may also be involved in interplay with other molecular processes that influence gene transcription, which may also depend on the SC-mechanism described in our work. In particular, previous experimental studies show that promoters of all genes can be divided into two large groups: those that contain CpG islands (∼ 70%) and those that do not (∼ 30%) [79–82]. CpG-island (CGI) promoters, which are associated with ‘housekeeping’ genes and a subset of developmental regulator genes [83, 84], have been shown to be nucleosome-deficient compared to other chromatin regions [85–88], including non-CpG-island (non-CGI) promoters [81], and therefore more accessible to TFs.

Based on the sensitivity of nucleosome stability to DNA sequence observed in *in vitro* experiments [89– 92], it has been suggested that nucleosome deficiency in CGI promoters may be sequence-dependent [81]. Indeed, bioinformatic analysis showed that nucleosomes in living cells are somewhat sensitive to DNA sequence, which may partially explain the nucleosome deficiency at yeast promoters [93–100]. However, unlike in yeasts, DNA sequences of CGI promoters in human cells were found to encode a strong tendency for nucleosome formation, which is completely opposite to the nucleosome deficiency in CGI promoters observed *in vivo* [88]. This highlights the limitations of DNA sequence in determining nucleosome stability in promoter regions and, consequently, promoter activity in higher eukaryotes. Instead, it was found that promoter activity is regulated by the PRC1 and PRC2 polycomb-group complexes through H3K27me3 nucleosome PTM and associated chromatin compaction, leading to the formation of so-called ‘bivalent’ CGI promoters, which account for ∼ 20% of CGI promoters in embryonic stem cells [82, 101, 102]. Therefore, it can be concluded that histone PTMs and the level of chromatin compaction play a much more important role in regulating CGI promoter activity in higher eukaryotes, possibly partly through the SC-mechanism, than nucleosome sensitivity to DNA sequence.

Furthermore, other experimental studies showed that promoters of actively transcribed genes are often enriched in nucleosomes that, instead of the usual H2A and H3 histones, contain their isoforms H2A.Z and H3.3 [103– 105], both of which are normally present in such nucleosomes [104, 106, 107]. Additionally, nucleosomes containing both H2A.Z and H3.3 were found to be typically localized to ‘nucleosome-free regions’ of active promoters [106], which appear to be nucleosome-deficient likely because such nucleosomes are less stable than normal ones [104, 105, 108]. Although the influence of H2A.Z histone on nucleosome stability has been debated in many studies showing that H2A.Z histones can have both stabilizing and destabilizing effects on nucleosomes [104, 105, 108– 112], potentially depending on the co-presence of H3.3 histones in the same nucleosomes [104], the overall conclusion appears to be that both H2A.Z and H3.3 used by living cells to fine-tune gene expression through the regulation of RNA polymerase pausing at transcription-poised promoters [105, 110–115]. Notably, H2A.Z and H3.3 histones in nucleosomes associated with promoters of active genes were found to be enriched in acetylation and H3K4me3 PTMs, typical of euchromatin [103, 116]. Yet, *in vitro* experiments have shown that the acetylation state itself has little effect on the stability of nucleosomes containing H2A.Z and H3.3 histones [104, 117]. Combined with the observation that H2A.Z and H3.3 histones are deposited in chromatin after assembly of the preinitiation complex [118], these results suggest that H2A.Z and H3.3 histones may be used by cells in conjunction with / downstream of molecular mechanisms involved in regulating chromatin condensation and accessibility to TFs. Such a multi-level regulation of gene transcription may be important for synchronization and reduction of cell-to-cell variability in gene expression during developmental processes [113, 119–121]. In the future, it will be interesting to explore how the SC-mechanism described in our study may be involved in interplay with H2A.Z and H3.3 histone isoforms to fine-tune gene transcription.

As for non-CGI promoters, unlike CGI promoters, they are typically associated with stable nucleosomes and rely on chromatin remodelling complexes, such as SWI/SNF, for their activation [81]. Indeed, previous studies show that by binding to the promoters of transcriptionally silent genes, pioneering TFs can, among other things, recruit chromatin remodelling complexes that use the energy of ATP hydrolysis (∼ 20 k_B_T [122]) to displace or remove nucleosomes, clearing the way for DNA binding to downstream TFs required for transcription activation [23, 123]. This mechanism allows for selective activation and tight regulation of the corresponding genes, minimizing their basal transcription and thereby preventing synthesis of gene products that may be harmful to cells if constitutively present in low concentrations [81].

Interestingly, although at some promoters the SWI/SNF remodelling complex catalyses nucleosome sliding, allowing pre-initiation complex assembly and gene transcription [124], at other promoters SWI/SNF-dependent remodelling coincides with increased accessibility of promoter DNA to TFs, but nucleosomes do not appear to be displaced or evicted [125, 126]. These results suggest that the primary function of SWI/SNF remodelling complexes may not be limited to nucleosome eviction / displacement. Indeed, recent experimental studies have shown that in addition to sliding and disassembling nucleosomes, SWI/SNF remodelling complexes can act as molecular ‘stir bars’, disrupting interactions between nucleosomes and locally decondensing chromatin [127]. According to our study, this may result in nucleosome destabilization via the SC-mechanism, providing a potential explanation for how SWI/SNF remodelling complexes can activate promoters without displacing or evicting nucleosomes. Interestingly, a very similar mechanism may also be responsible for the Top 2*β* topoisomerase-dependent gene regulation observed during neuronal differentiation, which has been suggested to be related to changes in the local chromatin condensation level caused by the activity of promoter-associated topoisomerases [128]. Thus, such an interplay between the activity of chromatin motor proteins, the level of chromatin condensation and the SC-mechanism in the regulation of gene transcription may be quite universal, warranting future study.

Finally, it should be noted that previous studies indicate that transcription of some genes may be further regulated via the formation of DNA loops by CTCF and cohesin protein complexes, which either help bring gene promoters into proximity with enhancers for gene transcription activation or create isolated regulatory chromatin regions that protect genes within the loop domain from the influence of enhancers outside it [9, 129–133]. Indeed, enhancers are short regions of chromatin enriched in nucleosomes containing histones with PTMs that are typically found in transcriptionally active chromatin, as well as chromatin-modifying enzymes required for the introduction of such PTMs, and, in addition, include chromatin remodelling complexes and TFs needed to activate gene transcription [134]. Thus, by bringing enhances closer to promoters via the DNA loop extrusion mechanism, cells can create a local microenvironment near them, similar to that found in euchromatin, which is additionally enriched in TFs. As a result, by engaging chromatin remodelling complexes and, possibly, the SC-mechanism, transcription of the corresponding genes is activated. Although only ∼ 0.5 − 1% of genes have been found to be regulated by the promoter-enhancer looping mechanism [9, 130, 131], it may still pay an important role, as it has been shown to promote the activation of genes involved in vital cell regulatory pathways [135], leading to downstream changes in the transcriptional activity of other genes through mechanisms other than promoter-enhancer looping. Indeed, the number of differentially expressed genes following CTCF depletion-induced disruption of DNA loops was found to increase more than 10-fold from day 1 to day 4 after CTCF depletion [130], suggesting that enhancer-controlled genes can have a significant impact on cell behaviour.

In particular, recent experimental studies indicate that the promoter-enhancer mechanism may play an important role in shaping the mechanosensitive behaviour of cells in response to the elastic properties of the surrounding microenvironment, for example, by altering, among other genes, the transcription level of myosin IIA heavy chain (MYH9) in a TEAD-YAP-dependent manner [136, 137]. Additionally, previous studies indicate that nuclear and chromatin organization may also play an important role in mediating mechanotransduction processes in living cells. In particular, it has previously been shown that mechanical forces and geometric constraints experienced by cells can influence the nucleocytoplasmic distribution of many TFs and chromatin-modifying enzymes, leading to changes in histone PTMs and rearrangement of heterochromatin domains, which collectively cause a shift in gene expression patterns [14, 43, 54, 65, 138–150]. Yet, the molecular mechanisms underlying these mechanotransduction processes remain largely unclear.

Our results suggest that there may be several mechanisms involved in these mechanotransduction processes. First, theoretical calculations based on the experimental data collected in our study indicate that mechanical force applied to DNA may be directly involved in regulating the balance between histone-based nucleoprotein complexes and other proteins, such as TFs, that compete with each other for DNA binding. In particular, calculations showed that while TFs can efficiently compete with histone-based complexes such as nucleosomes and tetrasomes in euchromatic regions, in heterochromatic regions DNA is largely inaccessible to TFs unless it is stretched by mechanical force, which has been found to significantly enhance TF binding to DNA by destabilizing nucleosomes / tetrasomes. This result is in good agreement with single-molecule experiments showing that mechanical forces can directly influence DNA-binding competition between histone-based nucleoprotein complexes and transcription-regulating factors [55], as well as with experimental studies showing that mechanical forces applied to a cell can be transmitted through the actin cytoskeleton, nesprins, and LINC protein complexes directly to the lamina network inlaying the nuclear envelope and associated with chromatin, influencing gene transcription [14, 54, 138, 151–154].

In addition, our results suggest that mechanical forces and geometric constraints experienced by cells may potentially influence nuclear organization and gene transcription by inducing changes in the nuclear electrostatic potential. Indeed, calculations based on experimentally measured changes in the mean nuclear electrostatic potential as a function of the elasticity of the substrate on which cells are grown predicted that the nuclear electrostatic potential could influence nucleosome stability by ∼ 3 k_B_T [compare Figure 5(b) to Figure S17(b), SI], making it easier for TFs to compete with histone-based nucleoprotein complexes for DNA binding in cells grown on soft substrates (1 kPa gel) compared to the case of rigid substrates (glass slides). This effect may be further enhanced by the reduced expression levels of H1 linker histones observed in cells grown on soft substrates, which may lead to further nucleosome destabilization [Figure S7(b)].

More importantly, in the case of soft substrates, the nuclear electrostatic potential was found to lose its correlation with the nucleoplasmic distribution of DNA, suggesting that other electrically charged molecules may play a key role in shaping it. For example, our experiments showed that the nuclear electrostatic potential is sensitive to the expression level of H1 linker histones, although several other electrically charged nuclear components likely also contribute to the change in the spatial distribution of nuclear electrostatic potential observed between cells grown on soft and rigid substrates. Combined with the findings regarding the role of spatial heterogeneity of the nuclear electrostatic potential in governing the nucleoplasmic distribution of electrically charged nuclear components and in regulating nucleosome stability and chromatin accessibility for TFs (via the SC-mechanism), it can be concluded that this mechanism may also significantly contribute to the mechano-dependent changes in gene transcription observed in previous studies. Since the elasticity of many living tissues is on the order of a few kPa or less [65], these results suggest that such changes in the spatial distribution of nuclear electrostatic potential may synergistically cooperate with other transcriptional regulatory mechanisms mentioned above to shape the behaviour of living cells in these tissues.

Finally, it should be noted that the role of nuclear electrostatic potential in nucleosome stabilization may also provide clues to the so-called C-value paradox. Specifically, previous studies have shown that C-values (i.e. genome sizes) vary greatly across species and that this value has no relation to the estimated number of genes and/or complexity of the organism [155, 156]. In particular, it has been shown that genome size does not reflect the number of genes in higher eukaryotes, as the majority of DNA appears to be ‘junk’ DNA, the biological function of which remains poorly understood because this DNA does not encode either protein or functional RNA [156]. Yet, the proportion of this DNA can reach *>* 95% of the genome size [156]. Moreover, the sequence of ‘junk’ DNA is not evolutionary conserved and is subject to frequent mutations [156–158] and rapid turnover [159–162]. However, despite this, the genome size remains relatively stable over long periods of evolution [160, 162], suggesting that it is the bulk amount of noncoding DNA, rather than its sequence, that is important [155].

Although the functional role of ‘junk’ DNA remains poorly understood, our study suggests that this DNA, which constitutes the largest part of the genome, may be one of the key elements involved in the formation and shaping of nuclear electrostatic potential. Indeed, previous observations of the preferential localization of HP1 proteins to simple DNA repeats and transposable elements [163], which are well-known typical elements of ‘junk’ DNA [164–171], as well as extensive methylation of these DNA regions and heterochromatin-like PTMs of nucleosomes associated with them [170, 171], suggest that ‘junk’ DNA serves as a scaffold for heterochromatin formation, thereby significantly contributing to the nuclear electrostatic potential landscape. Thus, ‘junk’ DNA may be one of the key factors involved in regulating the transcriptional state of nearby chromatin regions (via the SC-mechanism), working in conjunction with other transcriptional regulatory mechanisms.

Indeed, it has previously been shown that noncoding regions of DNA located near exons are less likely to be lost during evolution, suggesting that they may serve a specific biological function [159]. Moreover, the possible role of ‘junk’ DNA in the formation of nuclear electrostatic potential may explain the insensitivity of cells to its frequent mutations and rapid turnover, as it is the bulk amount of DNA, and not its sequence, that is important for the generation of nuclear electrostatic potential, since each base-pair carries the same electrical charge. In the future, comparison of the evolutionary rates of different regions of ‘junk’ DNA, as well as their nuclear localization in different species and changes in the transcription of closely located genes, may provide additional information about the potential biological function of ‘junk’ DNA.

In conclusion, our study suggests that nuclear electrophysiology may potentially play an important role in regulating gene transcription in combination with other previously established molecular mechanisms, thereby serving as an additional regulatory layer based on physical principles.

## AUTHOR CONTRIBUTIONS

Y.G., F.H., X.D. and J.Y. performed experiments. Y.G., F.H., X.D., J.Y., Y.L., F.L., C.H.Y., Y.W.C., A.W.H., E.H. and A.K.E. analysed the data. Y.G., F.H., X.D. and A.K.E. wrote the paper. A.K.E. designed the research, performed theoretical calculations and supervised the study.

## ACKNOWLEDGEMENT

We greatly appreciate the encouraging and insightful discussions with Dr. Jie Yan (MBI, Singapore), Dr. Alexander Bershadsky (MBI, Singapore) and Dr. Fazly Ataullakhanov (University of Pennsylvania, USA). We would like to thank Dr. Gang Li (Shenzhen Bay Laboratory, China) and Dr. Mao Li (Shenzhen Bay Laboratory, China) for providing HEK293T and MDA-MB-231 cell lines. Computation and data analysis were carried out at Shenzhen Bay Laboratory High Performance Computing and Informatics Core Facility. Also, we would like to thank Bioimaging Core of Shenzhen Bay Laboratory for providing cell imaging support. This work was supported by the start-up funds from Shenzhen Bay Laboratory (A.K.E.).

## METHODS

### Cell culture, constructs and transfection

Human embryonic kidney cells (HEK293T, ATCC CRL-3216) and human breast cancer cells (MDA-MB-231, ATCC HTB-26) were obtained from the laboratories of Dr. Gang Li (Shenzhen Bay Laboratory, China) and Dr. Mao Li (Shenzhen Bay Laboratory, China). All cell lines were cultured in Dulbecco’s modified Eagle medium (DMEM, LONZA) supplemented with 10% fetal bovine serum (FBS, Gibco), 1% GlutaMAX (Gibco), and penicillin/streptomycin (100 U/ml penicillin and 0.1 mg/ml streptomycin) at 37^◦^C, 5% CO_2_. Cells were tested for mycoplasma contamination. Cell transfection was conducted using jetPRIME reagent (Polyplus) in 35 mm glass-bottom Petri dishes (Cellvis, USA) coated with fibronectin according to the manufacturer’s protocol.

Negatively charged mEGFP construct (mEGFP WT) was created by amplifying the relevant part of pcDNA3.1-Zeo-mEGFP plasmid (Addgene, #214145) using PCR primers encoding BamHI and EcoRI restriction sites and subcloning the PCR product into pcDNA3.1(+) vector backbone (Invitrogen). Net neutrally charged mEGFP (mEGFP+3RK) and positively charged mEGFP (mEGFP+6RK) constructs were generated first by adding 3-fold and 6-fold amounts of lysine and arginine aminoacids (3 × RK and 6 × RK) to the C-terminus of mEGFP in pcDNA3.1-Zeo-mEGFP plasmid, respectively, and then amplified using PCR primers encoding BamHI and EcoRI restriction sites, followed by subcloning the PCR products into pcDNA3.1(+) vector backbone. Similarly, pcDNA3.1-H1.1-mCherry, pcDNA3.1-mCherry, pcDNA3.1-mScarlet, and pcDNA3.1-mRuby2 expression vectors were constructed by amplifying histone H1.1, mCherry, mScarlet, and mRuby2 cDNAs using PCR primers containing HindIII and XbaI restriction sites, followed by subcloning of the PCR products into pcDNA3.1(+) vector backbone.

pEGFP-N1-H2B, pEGFP-N1-H1.1, pEGFP-N1-NAP1L1, pEGFP-N1-NAP1L4, and pEGFP-N1-ASF1a expression vectors were generated by amplifying cDNAs of histone H2B, histone H1.1, and histone-binding chaperones NAP1L1, NAP1L4 and ASF1A using PCR primers containing BglII and BamHI restriction sites and by subcloning the obtained PCR products into the pEGFP-N1 vector backbone (Clontech). pmCherry-N1-H2B and pmCherry-N1-H3.1 expression vectors were constructed by amplifying histone H2B and histone H3.1 cDNAs using PCR primers containing BglII and NotI restriction sites, followed by subcloning the PCR products into the pmCherry-N1 vector backbone (Addgene, #87327). The lentiviral vector system, pLenti-CMV-MCS-EGFP-SV-Puro, encoding mEGFP WT and mEGFP+3RK, was constructed by amplifying the corresponding fragments of the above plasmids containing these constructs using PCR primers with XbaI and BamHI restriction sites and subcloning the resulting PCR products into the pLenti-humanized vector backbone (Addgene, #73582).

### Lentiviral transduction

A stable transfected MDA-MB-231 cell line was created via lentiviral-based plasmid transduction. Briefly, 3 · 10^6^ MDA-MB-231 cells were co-transfected with a lentiviral plasmid encoding either mEGFP WT or mEGFP+3RK construct, along with psPax2 and pMD2.G vectors at a ratio of 4 *µ*g : 2 *µ*g : 1 *µ*g using Lipofectamine 3000 (ThermoFisher Scientific) as the transfection reagent. After 24 h, the culture medium was replaced and virus collection was performed after another 36 h / 72 h. Viral aliquots were pooled and filtered using a 0.22 *µ*m filter. Then, 2 · 10^5^ MDA-MB-231 cells were transduced using a mixture of 1 ml freshly harvested virus and 1 ml DMEM with 5 *µ*g/ml Polybrene (Sigma-Aldrich), after which the transduced cells were seeded into 6-well plates.

### Preparation of polyacrylamide (PA) gel substrate

PA substrates for cell culturing were prepared following the protocol described in ref. [172]. Namely, the gel was prepared by mixing 40% AA (Sigma-Aldrich, #A9099) and 1% bis-AA (Sigma-Aldrich, #146072) water solutions with 20% NHS-AA ester (Tokyo Chemical Industry Co., Ltd, #H0623) DMSO solution. The NHS-AA mixture was incubated for 5 min at room temperature. The gel solutions were degassed for 30 min in vacuum. To activate polymerization, 10 *µ*l of 10% ammonium persulfate (APS, Beyotime, #ST005) and 1 *µ*l of tetramethylethylenediamine (TEMED, Beyotime, #ST728) were added to the solution (1% and 0.1% of total volume, respectively) and briefly mixed. Then, 25*/*30*/*69 *µ*l of the mixture were placed in 35 mm glass-bottom Petri dishes/12-well plates/6-well plates and covered with a glass coverslip. The sandwiched gels were incubated for 1 h at room temperature, after which the glass coverslip was removed. The polymerized gels were washed three times for 5 min with 2 ml of 1 × PBS to remove unreacted AA. Finally, gels were coated with 10 *µ*g/ml fibronectin overnight at 4^◦^C.

### Photolithography and microfluidic device fabrication

Preparation of microfluidic channels for cell migration assay using stably transfected MDA-MB-231 cells expressing mEGFP WT or mEGFP+3RK constructs was performed according to the protocol published in ref. [39].

### Live cell imaging and confocal microscopy

Following jetPRIME transfection, HEK293T cells were seeded at a density of 10^5^ cells/ml onto 35 mm glass-bottom Petri dishes or 1 kPa PA gel substrate coated with 10 *µ*g/ml fibronectin and cultured for 24 h. Before the experiment, to stain the nuclei, cells were treated with Hoechst 33258 dye (Thermo Fisher Scientific, #H3569) at a final concentration of 1 *µ*g/ml. After 15 min incubation, cells were imaged in Leibovitz’s L-15 medium without Phenol Red containing 10% FBS. Snapshots or timelapse images were acquired with a laser scanning confocal microscopy system (Nikon A1 HD25) based on an Nikon Eclipse Ti-2 inverted microscope, equipped with 60 × /1.4 Oil immersion objective and 37^◦^C cell incubator. Fluorescence excitation was carried out using 15 mW 405 nm, 100 mW 488 nm, 50 mW 561 nm and 100 mW 642 nm solid state lasers. In 3D confocal imaging experiments, the Z-stack step was 1 *µ*m.

### FRAP experiments

For FRAP experiments, HEK293T cells co-transfected with plasmids encoding mEGFP WT and H2B-mCherry or H3.1-mCherry constructs, or with Nap1L1-EGFP / Nap1L4-EGFP + H2BmCherry and ASF1a-GFP + H3.1-mCherry plasmid pairs, were cultured on a glass-bottom Petri dish for 24 h. In the case of HEK293T cells arrested in the Sphase of the cell cycle, cell cycle arrest was achieved by adding 2 mM hydroxyurea (final concentration) to the culture medium. FRAP experiments were performed using a confocal laser scanning microscope (LSM980, ZEISS) with a 60 × /1.4 Oil immersion objective. Fluorescence images of the GFP and mCherry channels were acquired every 15 s for 20 min using 488 nm and 561 nm lasers, respectively. Photobleaching of a circular region of the cell nucleus with a radius of 2 *µ*m was performed using a 488 nm laser for a time sufficient to reduce the average fluorescence intensity of the photobleached region to less than 10% of its original value. The power of the photobleaching laser was set low enough to avoid damage to other cellular components, such as stress fibers.

### Measurement of the pH level in the nuclei of living cells

To measure the pH level in HEK293T cells, the pHrodo™ Red AM Intracellular pH Indicator (ThermoFisher Scientific, #P35372) was used according to the manufacturer’s protocol. Briefly, cells seeded in a 96-well plate were loaded with 5 *µ*M pHrodo™ Red AM indicator for 30 min. The cells were then washed with Live Cell Imaging Solution™ and incubated with pHrodo™ AM staining solution for 30 min, followed by washing with Live Cell Imaging Solution™, after which the pH level in cells was measured using a confocal microscope. To calibrate pHrodo™ Red AM indicator, the cells used in the experiment were washed with Live Cell Imaging Solution™ and then incubated with Cell Loading Solution containing 10 *µ*M nigericin and 10 *µ*M valinomycin, as well as Cellular pH Calibration buffer to fix the intracellular pH at 4.5, 5.5, 6.5, or 7.5. By measuring cellular fluorescence in three independent experiments, a standard curve was generated for the pHrodo™ Red AM indicator in HEK293T cells, which demonstrated a linear relationship between intracellular pH and the fluorescence intensity of the indicator in the cells.

### Immunofluorescence experiments

HEK293T cells cultured in a glass-bottom Petri dish coated with fibronectin were fixed with 3.7% paraformaldehyde (PFA) at room temperature, then permeabilized and blocked with a mixture of 0.1% Triton X-100 and 2.5% bovine serum albumin (BSA) in 1 × PBS at room temperature for 30 min. The permeabilized cells were then treated with primary and secondary antibodies dissolved in a mixture of 1% BSA and 0.1% Tween-20 in 1 × PBS (PBST buffer). Specifically, the cells were incubated with primary antibodies to histone H3 (Abcam, #ab176842, 1/1000 dilution, the antibody recognizes the same peptide sequence in histones H3.1 and H3.3) and histone H2B (Abcam, #ab52484, 1/1000 dilution) overnight at 4^◦^C and then washed three times with 0.1% PBST. Secondary antibodies Goat Anti-Rabbit IgG H&L (Abcam, #ab175471, 1/1000 dilution) and Goat Anti-Mouse IgG (H+L), F(ab’)2 Fragment (CST, #4408, 1/1000 dilution) were incubated with the cells for 1 h at room temperature.

### Flow cytometry

HEK293T cells transfected with a plasmid encoding either mEGFP WT, mEGFP+3RK, or mEGFP+6RK using jetPRIME transfection agent (Polyplus) were cultured in a 6-well plate for 36 h, then harvested and washed with 1 × PBS before the experiment. Following passage through a 40 *µ*m cell strainer, the GFP signal was analysed using BD FACSAria™ III Cell Sorter flow cytometer. For each sample, 10000 events were recorded, with cell doublets and cellular debris (∼ 10%–20% of events) excluded from the analysis.

### Cell permeabilization experiments

HEK293T cells transfected with a plasmid encoding either mEGFP WT, mEGFP+3RK, or mEGFP+6RK using jetPRIME transfection agent (Polyplus) were cultured in glass-bottom Petri dishes for 36 h. Before the experiment, cells were washed with 1 × PBS. The dish was then filled with fresh culture medium and imaged using IXplore Spin SR (Evident) super-resolution spinning disk confocal microscope. For weak permeabilization of the cell membrane, the culture medium was replaced with permeabilization solution (100 mM CH_3_CO_2_K, 30 mM KCl, 10 mM Na_2_HPO_4_, mM DTT, 1 mM MgCl_2_, 5% w/v Ficoll 400, 1 mM ATP, and 1% Triton X-100), and cell images were acquired every 10 s to measure the diffusion rate of mEGFP probes out of the nuclei of permeabilized cells.

### Hypoosmotic shock experiments

Stably transfected MDA-MB-231 cells expressing mEGFP WT were cultured in glass-bottom flow chambers, which were precoated with rat tail collagen I (Solarbio) dissolved in 6 mM acetic acid at a concentration of 0.3 mg/ml, for 24 h. Before the experiment, cells were washed with 1 × PBS. The dish was then filled with fresh culture medium containing 1 *µ*g/ml Hoechst 33258 dye (Thermo Fisher Scientific, #H3569) and imaged using IXplore Spin SR (Evident) super-resolution spinning disk confocal microscope. To induce a hypoosmotic shock, the culture medium was replaced with a hypoosmotic solution consisting of 10% fetal bovine serum (Gibco), 1% penicillin-streptomycin (Biosharp), 44% Milli-Q water, and 45% DMEM (Gibco). Cells were imaged before and after hypoosmotic shock.

### Cell extracts and Western blotting

HEK293T cells transfected with a plasmid encoding either mEGFP WT, mEGFP+3RK, mEGFP+6RK, or H1.1-EGFP using jetPRIME transfection agent (Polyplus) were cultured in a 6-well plate for 36 h. Cell lysates were prepared by adding RIPA buffer (FansBio) to the 6-well plate and harvesting the cells, after which the lysates were incubated on ice for 30 min and centrifuged at 15000 rpm for 30 min at 4^◦^C using a benchtop centrifuge. The total protein concentration in the collected supernatant was determined using the Pierce™ Bradford Protein Assay Kit (ThermoFisher Scientific), and an equal amount of protein was loaded into each lane of SDS-PAGE gel (FansBio). After separation of proteins by electrophoresis, they were transferred to a PVDF membrane, followed by blocking it with 5% non-fat dry milk in PBST buffer (1 × PBS supplemented with 0.1% Tween 20) for 1 h at room temperature. The PVDF membrane was then incubated with primary antibodies to *β*-tubulin (Abmart, #M20005S, 1/5000 dilution) and to GFP (Abmart #M20004S, 1/1000 dilution) or histone H1 (Santa Cruz Biotechnology, #sc-393358, 1/1000 dilution, the antibody recognizes all histone H1 isoforms) at 4^◦^C overnight. The next day, after washing 3 times with PBST buffer, the membrane was incubated with Goat Anti-Mouse IgG HRP (Abmart, #M21001S, 1/1000 dilution) secondary antibodies for 1 h, then washed 3 times and visualized using SuperFemto ECL Chemiluminescence Kit (Vazyme, #E423-01) and ChemiDoc MP Imaging System (BioRad).

### Chromatin immunoprecipitation and digestion assay

HEK293T cells were cultured in 15 cm Petri dishes and fixed with 1% formaldehyde for 10 min. The reaction was stopped by adding 125 mM glycine solution, after which the cells were collected and lysed with lysis buffer (50 mM HEPES-KOH, 140 mM sodium chloride, 1 mM EDTA, 1% Triton X-100, 0.1% sodium deoxycholate, 0.1% SDS and protease inhibitor). The cell lysates were then sonicated to fragment genomic DNA into smaller parts using S220 Focused-ultrasonicator with a peak power of 105 W, a duty factor of 5%, 200 cycles, and a processing time of 40 s. After this, the samples were centrifuged at 8000 × *g* for 10 min, and the supernatant was collected. Then, antibodies to H3K4me3 (Proteintech, #91264) or H3K9me3 (CST, #13969T) were added to the supernatant and incubated in a rotator at 4^◦^C for 1 h to isolate the euchromatic or heterochromatic fraction, respectively. In the next step, chromatin samples were immunoprecipitated using protein A/G magnetic beads (Beyotime, #P2083S) by incubation overnight in a rotator at 4^◦^C. Finally, after washing with wash buffer (0.1% SDS, 1% Triton X-100, mM EDTA, 20 mM Tris-HCl, 150 mM NaCl), the chromatin-bound magnetic beads were treated with micrococcal nuclease (MNase, Beyotime, #D7201S) at a final concentration of 2 U/ml for 30 min. The reaction was stopped by adding 25 mM EDTA (final concentration), after which the beads were eluted with elution buffer (1% SDS, 100 mM NaHCO_3_). The eluate was incubated with 5 M NaCl and proteinase K (Beyotime, #D0063-6) to reverse cross-linking and release residual DNA fragments from DNA-binding proteins. DNA fragment size determination was performed by electrophoresis of the resulting DNA samples in a 3% agarose gel in TBE buffer.

### Chromatin native PAGE gel and Western blot assay

HEK293T cells were transfected with plasmids encoding H3.1-mCherry and H2B-EGFP constructs using jetPRIME transfection agent (Polyplus). After 72 h of culture, the cells were harvested, lysed, and then treated with 2 U/ml MNase (final concentration, Beyotime, #D7201S) in the same manner as described in the ‘Chromatin digestion assay’ section. After stopping the MNase digestion reaction with EDTA, the products were separated by electrophoresis in a 5% native PAGE gel in TBE buffer, then transferred to a PVDF membrane, after which the membrane was blocked with 5% non-fat dry milk in PBST buffer (1 × PBS supplemented with 0.1% Tween 20) for 1 h at room temperature. The PVDF membrane was then incubated with primary antibodies to mCherry (Proteintech, #26765-1-AP, 1/1000 dilution) and to GFP (Abmart, #M20004S, 1/1000 dilution) at 4^◦^C overnight. The next day, after washing 3 times with PBST buffer, the membrane was incubated with Goat Anti-Rabbit IgG HRP (CST, #7074, 1/10000 dilution) and Goat Anti-Mouse IgG HRP (Abmart, #M21001S, 1/10000 dilution) secondary antibodies for 1 h, then washed 3 times and visualized using SuperFemto ECL Chemiluminescence Kit (Vazyme, #E423-01) and ChemiDoc MP Imaging System (BioRad).

### Image analysis

Collected cell images were analysed with the help of ImageJ-win64 and Matlab software. In 3D confocal imaging experiments, the nuclear volume was calculated by measuring the nucleus projected nuclear area on each Z-stack slice, then summing all the resulting areas and multiplying the obtained sum by the Z-stack step (1 *µ*m).

In live cell imaging experiments, the correlation between two different fluorescent channels of a cell nucleus image was calculated for each pixel, *j*, as follows:

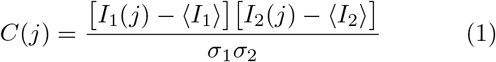

Where *C*(*j*) is the correlation coefficient between channels 1 and 2 at the *j*^th^ pixel (*j* = 1…*N*_pix_, *N*_pix_ is the total number of pixels in the image of the cell nucleus). *I*_1_(*j*) and *I*_2_(*j*) are the *j*^th^ pixel brightness in channels 1 and 2, correspondingly. ⟨*I*_1_⟩ and ⟨*I*_2_⟩ are the average intensities of the nucleus in channels 1 and 2. *σ*_1_ and *σ*_2_ are the standard deviations of the fluorescent signals in channels 1 and 2.

The local nuclear electrostatic potential in cells expressing mEGFP WT probes was calculated for each pixel of the cell nucleus image by measuring the mEGFP WT fluorescence intensity and then using an experimentally obtained calibration curve as described in Appendix B.

The FRAP experimental data were fitted using the diffusion model described in ref. [173] and a compartmental model similar to that described in ref. [174], see Appendix E for details.

All cell and nuclear images presented in the article are either original images or slices of 3D stacks. Z-stack averaging was not used to create the images.

## DATA AVAILABILITY

All data supporting the findings of this study is presented in the main text and Supplementary Information.

## CODE AVAILABILITY

The main program codes can be downloaded from websites artem-efremov.org or artemefremovlab.com. Other programs are available upon request.

## SUPPLEMENTARY INFORMATION

## Appendix A: Gibbs-Donnan effect and nuclear electrostatic potential

The nuclear envelope (NE) serves as a semipermeable membrane that allows ions and small cellular metabolites and proteins to pass through nuclear pore complexes [28, 175] while preventing large biomolecules, such as DNA, from leaving the cell nucleus. Since DNA carries a strong electrical charge, this leads to establishment of the Gibbs-Donnan equilibrium, which is characterized by different cytosolic and nucleoplasmic concentrations of electrically charged molecules that can move freely through the nuclear pore complexes, resulting in an electrostatic Donnan potential between the nucleus and the cytosol, see Figure 1(a). The value of this electrostatic potential can be found from the condition of electroneutrality, which must be satisfied for both the nucleus and the cytosol:

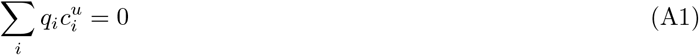

Where *q*_*i*_ and 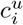 are the electrical charge and concentration of the i^th^ type of molecule in compartment *u* = n or c (nucleus or cytosol, respectively).

In the case of the cytosol, the sum in Eq. (A1) is mainly determined by the contribution of monovalent ions, such as K^+^, Na^+^, Cl^*−*^, as well as by small cellular metabolites, which on average have a negative charge of 1∼ − *q*_e_ [176–179], where *q*_e_ = 1*e* = 1.6 · 10^*−*19^ C is the elementary electrical charge. Indeed, previous studies suggest that the total cytosolic concentrations of positive and negative monovalent ions and small metabolites are maintained at a relatively stable level of 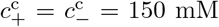 in living cells [176–179]. In contrast, the average protein concentration in living cells was found to be only ∼ 0.3 − 2.5 mM [180], i.e., two orders of magnitude lower than the total concentrations of positive and negative monovalent ions and small cellular metabolites. Thus, to a first approximation, the contribution of cytosolic proteins, which typically carry a fairly low electrical charge, can be neglected in Eq. (A1).

As for the nucleus, in addition to monovalent ions and electrically charged cellular metabolites, the main contribution to the average electrical charge density is made by negatively charged DNA and positively charged chromatin proteins, such as histones. Therefore, Eq. (A1) can be rewritten in the following form:

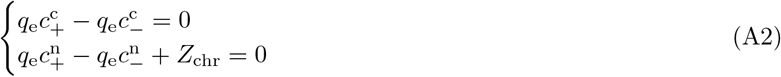

Where 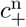 and 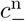 are the total concentrations of positively and negatively charged monovalent ions and small cellular metabolites in the nucleus. *Z*_chr_ is the net average electrical charge density of chromatin in the nucleus, which includes the electrical charges of DNA and histones. For example, in the case of HEK293T cells grown on a glass surface coated with fibronectin, the average nuclear volume is *V*_nucl_ = 1130 ± 20 *µ*m^3^ [mean ± SEM, *N* = 15 cells, Figure 3(a)]. The total length of the chromosomal DNA is ∼ 6.2 Gbp [181], with each basepair carrying an electrical charge of − 2*e*. Thus, the average electrical charge density of DNA in the nucleus is ∼ − 18 *e* mM. Since histone octamers that serve as protein cores of nucleosomes each carry a positive charge of ∼ +149*e* (based on the primary histone sequence), and the average nucleosome spacing on DNA is ∼ 190 bp [182], it can be found that the net average electrical charge density of chromatin in the nucleus is *Z*_chr_ ≈ − 11 *e* mM.

To close Eq. (A2), it should be supplemented by the following formulas for the Donnan potential, *φ*_n_, which arises between the nucleus and the cytosol due to the difference in the cytosolic and nucleoplasmic concentrations of electrically charged ions and small cellular metabolites that can move freely between the two compartments:

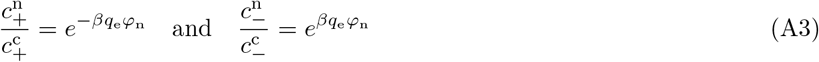

Where *β* = 1*/k*_B_*T* is the reciprocal of the thermodynamic temperature.

Combining Eq. (A2) and Eq. (A3), it is easy to obtain the following formula for the nuclear electrostatic potential:

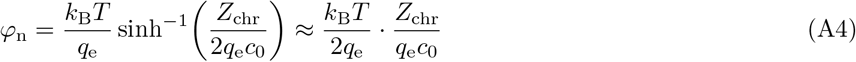

Here 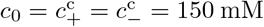 is the total concentration of monovalent ions and electrically charged cellular metabolites in the cytosol. sinh is the inverse of the hyperbolic sine function (i.e., arcsinh). In the right-hand side of the above equation, we used sinh *x* ≈ *x* approximation, which is valid for *x* ≪ 1, since *Z*_chr_ ≪ 2*q*_e_*c*_0_.

Given the average electrical charge density of chromatin in the nucleus (*Z*_chr_ ≈ − 11 *e* · mM), it is easy to find from Eq. (A4) that the average electrostatic potential of the nuclei of HEK293T cells grown on fibronectin-coated glass slides should be of the order of *φ*_n_ ≈ − 1.0 mV, which is in good agreement with the theoretical predictions obtained by solving the Poisson-Boltzmann equation (*φ*_n_ ≈ − 1.0 mV, [25]) and is comparable to the experimental measurements performed in the present study [*φ*_n_ = − 0.333 ± 0.018 mV, mean ± SEM, *N* = 63 cells, Figure 3(c,d)]. The difference between the theoretical and experimentally measured values of the nuclear electrostatic potential may be due to the fact that, in addition to the core histones, other positively charged DNA-binding proteins can also contribute to the net average electrical charge density of chromatin in the cell nucleus, for example, the positively charged linker histone H1 (*q* = +53.0*e*).

Indeed, previous experimental studies show that the expression level of H1 linker histones in mammalian cells is comparable to the expression level of core histones H2A, H2B, H3 and H4 [183]. Direct calculations show that mammalian cell nuclei contain, on average, *N*_n_ = 6.2 Gbp*/*190 bp ≈ 3.3 · 10^7^ nucleosomes, which corresponds to ∼ 100 *µ*M average concentration of each core histone in the nucleus. Assuming that the nuclear concentration of H1 linker histones is the same, it can be estimated that the average electrical charge density of H1 linker histones should be of the order of +5 *e* mM. Thus, H1 linker histones compensate for a significant part of the average electrical charge density of chromatin *Z*_chr_, resulting in *Z*_chr_ ≈ −6 *e* · mM. Using Eq. (A4), it can be found that the latter corresponds to an average nuclear electrostatic potential of the order of *φ*_n_ ≈ −0.5 mV, which is in better agreement with the experimentally measured value of *φ*_n_ = − 0.333 ± 0.018 mV.

Notably, the above approach can be used not only to predict the average value of the nuclear electrostatic potential, but also to describe its spatial heterogeneity on a finer scale, caused by the phase-separation of chromatin into hetero- and euchromatin domains, see Figure 1(b). Using an electroneutrality condition similar to that in Eq. (A2), it can be shown that the difference in the electrical charge densities in dense heterochromatin domains and sparse euchromatin domains results in a Donnan electrostatic potential 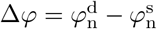 between these domains, where 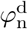 and 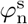 are the electrostatic potentials of densely and sparsely packed chromatin domains, characterized by the following relationship:

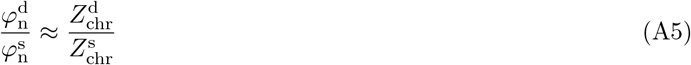

Here 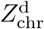 and 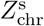 are the average electrical charge densities of chromatin in the corresponding domains (‘d’ – densely packed heterochromatin, ‘s’ – sparsely packed euchromatin).

## Appendix B: Calibration of the mEGFP WT probe

To map the electrostatic potential in the nuclei of living cells and assess the relative effect of electrostatic and volume-exclusion interactions on the nucleoplasmic distribution of mEGFP WT, we calibrated the mEGFP WT probe using another fluorescent protein, mScarlet, which has the same size as mEGFP WT but carries a different net electrical charge (mEGFP WT *q* = − 7.3*e*, mScarlet *q* = − 3.6*e*). Specifically, the FRAP experiments performed in our study show that the fluorescent signal of mEGFP WT is recovered very quickly in the photobleached area, – in less than 15 s [Figure S15]. This result suggests that fluorescent probes quickly reach thermodynamic equilibrium in the nuclei of living cells, and therefore their nucleoplasmic distribution should obey the well-known Boltzmann law:

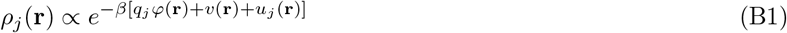

Here *ρ*_*j*_(**r**) is the nucleoplasmic distribution of the *j*^th^ probe (*j* = 1 mEGFP WT, and *j* = 2 mScarlet) as a function of the position vector **r**. *β* = 1*/k*_B_*T* is the reciprocal of the thermodynamic temperature. *q*_*j*_ is the net electrical charge of the *j*^th^ probe. *φ*(**r**) is the nuclear electrostatic potential at the point corresponding to the position vector **r**. *v*(**r**) is the strength of volume-exclusion interactions of the probes with the surrounding microenvironment, which is assumed to be the same for both probes since have the same size. In particular, *v*(**r**) describes the local volume-excluded fraction of space (*F*_ex_) due to the interaction of probes with chromatin and other nuclear components: *F*_ex_(**r**) = 1 − *e*^*−βv*(**r**)^ ≈ *βv*(**r**) if *v*(**r**) ≪ *k*_B_*T* . *u*_*j*_(**r**) is the energy term describing other types of interactions of the *j*^th^ probe with the surrounding microenvironment, except for the volume-exclusion interaction and the leading term of the electrostatic interaction, which are described by the fields *φ*(**r**) and *v*(**r**). I.e., *u*_*j*_(**r**) includes multipole electrostatic interactions, as well as interactions of other nature with the surrounding microenvironment.

Assuming that the intensity of the fluorescent probes, *I*_*j*_(**r**), is proportional to their density, *ρ*_*j*_(**r**), it follows that:

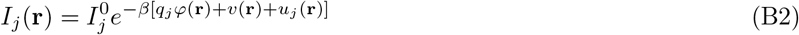

Where 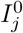 is the average intensity of the *j*^th^ probe in the cell cytoplasm, which in we use as a reference point for the electrostatic field *φ*(**r**), as well as *v*(**r**) and *u*_*j*_(**r**) energies. In the experiments, 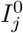 can be measured as the average intensity of the brightest regions of the cytoplasm (i.e., free of organelles blocking the fluorescent probes).

From Eq. (B2) it is easy to find that:

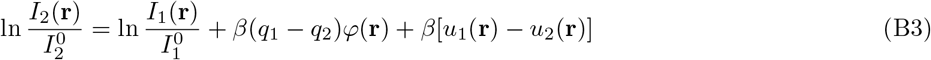

The above formula suggests that it is possible to map the nuclear electrostatic potential by measuring the logarithms of 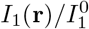 and 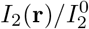 fluorescent signal ratios at each pixel of the cell nucleus image, within the error due to the energy term *β*[*u*_1_(**r**) − *u*_2_(**r**)]. The latter can be roughly estimated by measuring the offset between the logarithms of 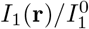 and 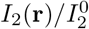 fluorescent signal ratios as schematically shown in Figure S5, or the width of the data point distribution. Measurements of this offset show that the value of the *u*_1_(**r**) − *u*_2_(**r**) term (− 0.033 ± 0.004 *k*_B_*T*, mean ± SEM, *N* = 13 cells) is on average an order of magnitude smaller than the electrostatic and volume-exclusion interaction energies of the mEGFP WT and mScarlet probes with the surrounding microenvironment [0 − 0.7 *k*_B_*T*, see Figure S6(c,d)], indicating that this protein pair is well suited for accurate mapping of the nuclear electrostatic potential and volume-exclusion interactions in the nucleus.

In comparison, experiments performed using mRuby2 instead of mScarlet showed that mRuby2 has strong interactions with the surrounding microenvironment in the nucleus that are unlikely to be due to the volume-exclusion interactions and the leading term of electrostatic interactions. This often results in a significant shift between the logarithms of 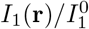 and 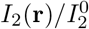 fluorescent signal ratios measured for mEGFP WT and mRuby2, which varies from one cell to another, leading to much greater scattering of the experimental data points, see Figure S6(a). Thus, unlike the case of mEGFP WT + mScarlet, mRuby2 cannot be used together with mEGFP WT to accurately map nuclear electrostatic potential due to the strong contribution of the term *β*[*u*_1_(**r**) − *u*_2_(**r**)] in Eq. (B3).

Furthermore, measurements of the logarithms of the 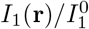 and 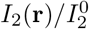 fluorescent signal ratios showed that mEGFP WT and mScarlet probes potentially interact with nuclear components outside and inside the nucleoli in different ways, see Figure S6(b). Since in this study we were interested in mapping the nuclear electrostatic potential and volume-exclusion interactions in the nucleoplasm outside the nucleoli, in all calculations the fluorescent signal ratios 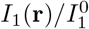 and 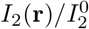 were considered only for regions outside the nucleoli.

Taking into account that for a good pair of fluorescent proteins, such as mEGFP WT and mScarlet, the term *β*[*u*_1_(**r**) − *u*_2_(**r**)] makes a small contribution to Eq. (B3), one can obtain the following formula for the nuclear electro-static potential:

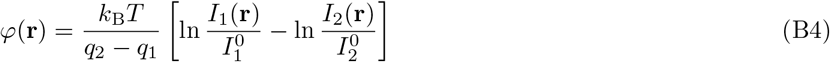

Substituting Eq. (B4) into Eq. (B2), it is easy to find that:

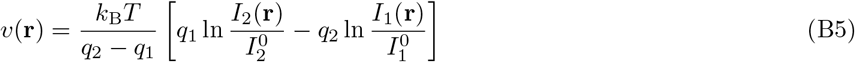

It should be noted that Eq. (B4) and Eq. (B5) are only valid if the fluorescent proteins are truly monomeric and do not form dimers or higher-order multimers. Experimental assays, such as the OSER test, show that this is essentially true for the mEGFP WT probe, which has an OSER index value of 98.8, indicating that mEGFP WT only exhibits a tendency to dimerize 1.2% of the time in the OSER test [184]. On the other hand, mScarlet has an OSER index value of 80, showing that it is prone to weak dimerization in crowded microenvironments [185].

For fluorescent proteins that can form dimers, Eq. (B4) and Eq. (B5) should be slightly modified. Specifically, it can be shown that in the case where the energies of electrostatic and volume-exclusion interactions of fluorescent probes with the surrounding microenvironment are less than 1 *k*_B_*T* [which is the case for the mEGFP WT and mScarlet probes, see Figure S6(c,d)], the following approximate formulas can be obtained:

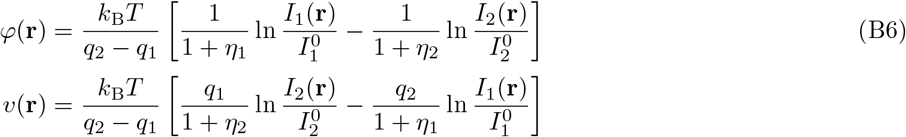

Here *η*_1_ and *η*_2_ are the fractions of probes 1 and 2 that form dimers in the nucleus. In all calculations shown in Figure S6(c,d), the OSER index value was used as a first-order approximation to the dimerization tendency of mEGFP WT and mScarlet fluorescent proteins by using *η*_*j*_ = 1 − (OSER index)_*j*_*/*100. Thus, for mEGFP WT: *η*_1_ = 0.012, while for mScarlet: *η*_2_ = 0.2.

Using Eq. (B6), it was found that the electrostatic and volume-exclusion interaction energies of mEGFP WT probes with the nuclear microenvironment in HEK293T cells grown on fibronectin-coated glass slides were linearly correlated with the logarithm of the fluorescence intensity ratio 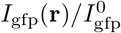 measured for mEGFP WT, see Figure S6(c,d). Thus, after performing such calibration of mEGFP WT with mScarlet, it becomes possible to use only the mEGFP WT probe to map the electrostatic potential and strength of volume-exclusion interactions in the nuclei of HEK293T cells. Very similar calibration curves using the mEGFP WT and mScarlet pair were also obtained for HEK293T cells grown on fibronectin-coated 1 kPa gel.

## Appendix C: Fraction of the nucleus occupied by chromatin and the relative contribution of electrostatic and volume-exclusion interactions to nucleosome energy

By calibrating the mEGFP WT probe as described in Appendix B, it becomes possible to experimentally measure the energy of its volume-exclusion interaction with the surrounding microenvironment. Knowing this energy, one can approximately estimate the average fraction of the nuclear volume occupied by nuclear components. Specifically, assuming that chromatin is the primary contributor to the volume-exclusion interaction energy of mEGFP WT probes, and considering that nucleosomes are the primary components of chromatin and likely make contribute the most to the volume-exclusion interaction energy, the volume occupied by chromatin can be estimated as follows.

First, crystallographic studies show that nucleosomes are barrel-shaped nucleoprotein complexes with a diameter of ∼ 12 nm and a height of ∼ 7 nm [186], the volume of which (*V*_0_ ≈ 790 nm^3^) corresponds to a sphere with a radius *R*_n_ ≈ 5.7 nm, which we denote as the characteristic radius of nucleosomes. Similarly, it was previously found that GFP has a barrel shape with a diameter of ∼ 3 nm and a height of ∼ 4 nm [187], which corresponds to a sphere with a radius *R*_gfp_ ≈ 1.9 nm (i.e., the characteristic radius of mEGFP WT probes). Knowing these characteristic radii, the excluded volume experience by mEGFP WT, which is created by a single nucleosome, can be approximated as:

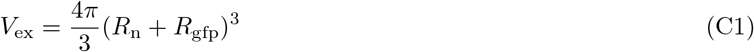

Since the volume occupied by the nucleosome itself is 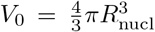, it can be seen that the excluded volume experienced by mEGFP WT is larger than the actual volume occupied by each nucleosome. The ratio of these two volumes is equal to:

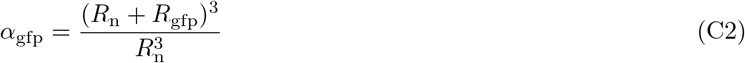

Substituting *R*_n_ ≈ 5.7 nm and *R*_gfp_ ≈ 1.9 nm into Eq. (C2), it can be found that *α*_gfp_ ≈ 2.4. Therefore, if *v*(**r**) is the local energy of volume-exclusion interaction of mEGFP WT probes with the surrounding microenvironment at the point corresponding to the position vector **r**, then the volume fraction occupied by chromatin can be estimated as:

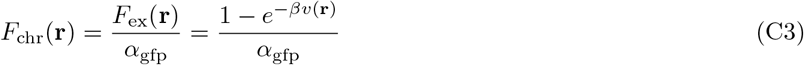

Where *F*_ex_(**r**) is the local volume-excluded fraction of space experienced by mEGFP WT probes due to their interaction with the surrounding microenvironment, see Appendix B.

By mapping the distribution of mEGFP WT probes in the nuclei of HEK293T cells grown on fibronectin-coated glass slides and using the calibration curve shown in Figure S6(d), it was estimated from Eq. (C3) that, on average, *F*_chr_ = 2.83 ± 0.12% (mean ± SEM, *N* = 21 cells) of the nuclear volume (excluding nucleoli) should be occupied by chromatin. On the other hand, knowing the total length of DNA (6.2 Gbp [181]) and the average nucleosome spacing on DNA (∼ 190 bp [182]), as well as the volume occupied by each nucleosome (*V*_0_ ≈ 790 nm^3^) and the average volume of HEK293T cell nuclei [*V*_nucl_ = 1130 ± 20 *µ*m^3^, mean ± SEM, *N* = 15 cells, Figure 3(a)], it is easy to find that nucleosomes occupy ∼ 2.3% of the nuclear volume. Thus, both methods yield comparable estimates of the fraction of excluded nuclear volume, indicating that the calibration curves shown in Figure S6(c,d) provide physically realistic values of the interaction energy of mEGFP WT probes with the surrounding microenvironment.

It is worth noting that previous studies on chromatin organization in the nuclei of living cells using electron microscopy tomography reported that the local fraction of nuclear volume occupied by chromatin ranges from ∼ 10% to 50% [188]. However, independent studies have shown that such high measured chromatin density may result from flattening of the nucleus caused by a several-fold reduction in the nucleus height occurring during cell fixation [189], a sample preparation step employed in the experiments described in ref. [188]. Furthermore, in ref. [188], the fraction of the nuclear volume occupied by chromatin was estimated by calculating the projected area of chromatin on tomographic slices, which may lead to an additional overestimation of the chromatin-occupied nuclear volume if a proper 3D reconstruction of the chromatin fiber is not performed. If both of these factors are taken into account (the reduction in the nuclear volume and the potential overestimation of the nuclear volume occupied by chromatin resulting from the measurement of the chromatin projected area), then the resulting estimates of the nuclear volume occupied by chromatin fall within the same range of values as those found in our study.

Using an approach similar to the one described above, one can also estimate the interaction energy of nucleosomes with the surrounding microenvironment caused by the volume-exclusion effect. In particular, in the case of a pair of interacting nucleosomes, Eq. (C2) becomes:

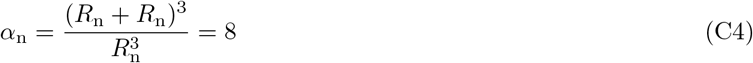

Similarly, Eq. (C3) turns into:

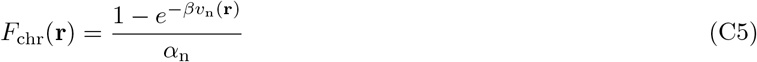

Where *v*_n_(**r**) is the volume-exclusion interaction energy of nucleosomes with the surrounding microenvironment. From Eq. (C3) and Eq. (C5) it then follows that:

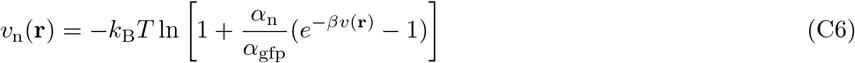

It should be noted that Eq. (C6) is based on the assumption that each nucleosome contributes the same effective excluded volume, *α*_n_*V*_0_, which is sensed by other nucleosomes. This is a valid approximation when the average inter-nucleosomal distance is greater than 4*R*_n_ (i.e., *F*_chr_(**r**) ≲ 0.1). Using Eq. (C3), it can be found that this condition corresponds to *v*(**r**) ≲ 0.3 *k*_B_*T*, which covers most of the experimentally measured data points shown in Figure S6(d). However, in the case of denser chromatin regions, a more elaborate approach is required to estimate the volume-exclusion interaction energy of nucleosomes by taking into account the overlaps between the volume-exclusion regions of adjacent nucleosomes.

Experimental measurements of volume-exclusion interactions in the nuclei of HEK293T cell grown on fibronectin-coated glass slides using the mEGFP WT probe and calculations based on Eq. (C6) show that the average difference between the volume-exclusion interaction energies of nucleosomes in chromatin regions of low and high electrostatic potential, which are often associated with heterochromatic and euchromatic domains, respectively, is 0.72 ± 0.04 *k*_B_*T* (mean ± SEM, *N* = 21 cells). On the other hand, the average difference in the nuclear electrostatic potential of the same regions is 0.617 ± 0.026 mV [mean ± SEM, *N* = 21 cells, also see Figure 3(d)], which, after multiplying by the electrical charge of the histone octamer (∼ +149*e*), corresponds to a difference of 3.44 ± 0.15 *k*_B_*T* (mean ± SEM, *N* = 21 cells) between the electrostatic interaction energies of histone octamers in sparse and dense chromatin regions. These estimates indicate that electrostatic interactions likely play a much more important role in influencing nucleosome stability compared to the volume-exclusion effect.

More careful data processing showed that if, instead of calculating the local average electrostatic potentials of dense and sparse chromatin regions, one calculates the statistical sum over the brightest and faintest parts of HEK293T cell nuclei in the GFP channel (excluding nucleoli), weighted by the local chromatin density, then the estimated difference between the energies of electrostatic interactions of nucleosomes with the surrounding microenvironment in the corresponding regions of the nuclei turns out to be even larger: 7.1 ± 0.5 *k*_B_*T* (mean ± SEM, *N* = 21 cells), see Appendix G and Table T3.

Interestingly, based on the measured average difference in the nuclear electrostatic potentials of heterochromatic and euchromatic regions, it can be found that nucleosomes with a net electrical charge of *q* = − 145*e* should have ∼ 3.4 *k*_B_*T* higher interaction energy with the local nuclear electrostatic potential in heterochromatic regions compared to euchromatic regions. Combined with the aforementioned ∼ 0.7 *k*_B_*T* difference between the energies of volume-exclusion interactions of nucleosomes in heterochromatin and euchromatin regions, these results indicate that, to condense chromatin and form heterochromatic domains, living cells must use molecular mechanisms capable of overcoming the ∼ 4 *k*_B_*T* energy difference between nucleosomes located in heterochromatic and euchromatic regions. Experimental studies suggest that one of such molecular mechanisms may be the cross-linking of nucleosomes by HP1 protein, which is associated with the formation of heterochromatin [7, 8, 69]. Another possible mechanism may be mediated by H1 linker histones, which are typically associated with heterochromatic regions and carry a very strong positive electrical charge (*q* = +53.0*e*) [77]. By binding to nucleosomes, H1 linker histones shield a large part of the nucleosome negative charge, thereby likely reducing their repulsion interaction with the local nuclear electrostatic potential and further stabilizing nucleosomes, as mentioned in the Discussion section.

## Appendix D: Correlation between cellular and nuclear volumes

To fit experimentally measured correlation between cell volume (*V*_cell_) and nuclear volume (*V*_nucl_), we used the mathematical model from ref. [25, 39], which postulates that nuclear size is determined by a mechanical equilibrium corresponding to a net zero pressure exerted on the nuclear envelope (NE). It was previously shown that in general this mechanical equilibrium includes three main pressure terms [25]:

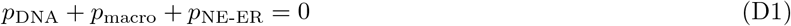

Where *p*_DNA_ is the pressure created by chromosomal DNA on the NE, *p*_macro_ is the osmotic pressure exerted by cytosolic macromolecules that cannot move through nuclear pore complexes (NPCs) into the nucleus, and *p*_NE-ER_ is the pressure resulting from the tug-of-war between the NE and the endoplasmic reticulum (ER) over the shared lipid membrane, which in higher eukaryotes arises mainly due to the polymerization of the lamina network underlying the NE.

In ref. [25, 39] it was shown that the DNA pressure term plays an important role only in the case of small nuclei (*V*_nucl_ ≥ 350 *µ*m^3^). On the other hand, in larger nuclei (*V*_nucl_ *>* 350 *µ*m^3^), such as in the case of HEK293T cells [Figure 3(a)], it plays a rather minor role. Thus, to fit experimental data shown in Figure 3(a), it is necessary to consider only *p*_macro_ and *p*_NE-ER_ terms, which have the following form [25, 39]:

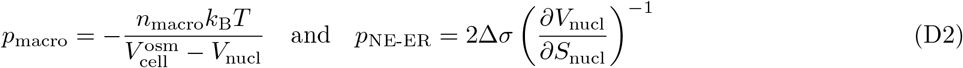

Where 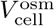 is the osmotically active volume of the cell, which usually occupies *ω* ≈ 70% of the total cell volume: 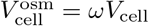 [190]. *n*_macro_ is the total number of macromolecules in the cell cytosol, which can be considered approximately proportional to the total number of proteins in a living cell: *n*_macro_ ≈ *ζc*_pr_*V*_cell_. Here *ζ* is the proportionality coefficient (0 *< ζ <* 1), and *c*_pr_ = 2.7 · 10^6^ proteins/*µ*m^3^ is the average protein concentration in mammalian cells [180]. The minus sign in the formula for *p*_macro_ indicates that pressure on the NE is exerted by macromolecules from outside of the nucleus. *S*_nucl_ is the surface area of the nucleus. 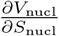 is the partial derivative of the nuclear volume with respect to the surface area of the nucleus, which, as previous experiments have shown, is constant over a wide range of nuclear sizes [39]. Δ*σ* = *σ*_ER_ − *σ*_NE_ is the difference between the surface tensions (i.e., surface free energy densities) of the NE and ER membrane.

From Eq. (D1) and Eq. (D2) it follows that the mechanical equilibrium of large nuclei is described by the following formula:

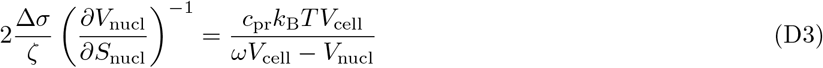

To a first approximation, Δ*σ* can be considered a constant (Δ*σ* ≈ const) in the above equation. This leads to the following linear correlation between cell and nuclear volumes, which describes the experimental data obtained on HEK293T cells quite well [see Figure 3(a)]:

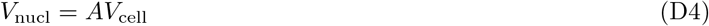

Here the proportionality coefficient *A* is equal to:

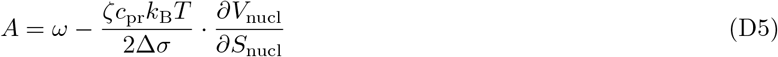

## Appendix E: Fitting experimental data collected in FRAP experiments

To fit experimental data collected in FRAP experiments, the FRAP signal measured in the nuclei of HEK293T cells was first double normalized to account for the loss of fluorescent signal due to the photobleaching pulse and bleaching during post-photobleaching imaging:

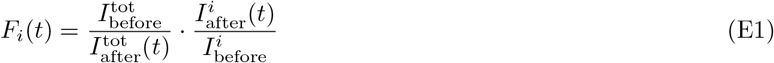

Here *F*_*i*_(*t*) is the normalized FRAP signal measured in dense (*i* = 1) and sparse (*i* = 2) chromatin regions as a function of time, *t*, see Figure 4(a,b) in the main text. 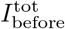 and 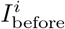 are the integral intensities of the entire nucleus and the *i*^th^ chromatin region (*i* = 1, 2) before the photobleaching pulse. 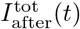 and 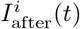 are time-dependent fluorescent intensities of the entire nucleus and the *i*^th^ chromatin region measured in FRAP experiments after the photobleaching pulse. All intensity values in Eq. (E1) are taken with background noise subtracted.

In experiments, it was found that the normalized FRAP signal recovers in two stages, see Figures 4(b) and S9. The first stage is a partial rapid recovery of the fluorescent signal due to the diffusion of freely moving histone dimers transported by chaperones, while the second stage is due to the exchange of histone dimers between the freely diffusing fraction and those associated with DNA in the form of nucleosomes, see Figure S11. To fit the initial rapid recovery of the FRAP signal during the first couple of minutes after photobleaching, we fitted the normalized FRAP signal to the following fluorescence recovery formula [173]:

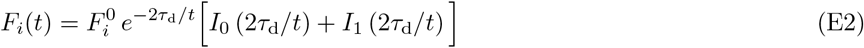

Here 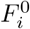 is the fraction of the FRAP signal recovered due to the diffusion of freely moving histone-chaperone complexes, and *τ*_d_ = *R*^2^*/*4*D* is the characteristic diffusion time of these complexes, where *R* is the radius of the photobleached area, and *D* is the diffusion coefficient of histone-chaperone complexes in the nucleoplasm [173]. *I*_0_ and *I*_1_ are modified Bessel functions of the first kind.

The subsequent slow recovery of the FRAP signal after initial equilibration of the freely diffusing histone-chaperone complexes between the unbleached and photobleached regions is likely due in part to the exchange of fluorescently-labelled histones between the histone-chaperone complexes and nucleoprotein complexes formed on DNA, see Figure S11. Thus, this part of the FRAP signal was fitted using a kinetic model similar to that described in ref. [174]. In particular, the model considers three distinct regions of the nucleus (Figure S13): 1) dense and 2) sparse chromatin regions selected within the photobleached nucleus area, and 3) the rest of the nucleus. The change in fluorescent signal in these three regions over the long-term (minutes to hours) due to the exchange between photobleached DNA-bound histones and freely diffusing fluorescent ones transported by chaperones is a kinetic process that can be described by the following master equation:

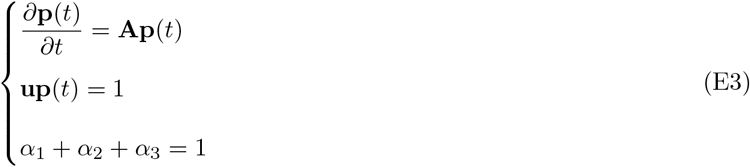

Where **p**(*t*) is the probability vector that a fluorescent histone dimer is bound to DNA in the dense chromatin region (probability *p*_1_), or in the sparse chromatin region (probability *p*_2_), or outside these two regions (probability *p*_3_), or is in the freely diffusing fraction of histones (probability *p*_f_); and **A** is a matrix that describes the kinetic transition rates between these four states:

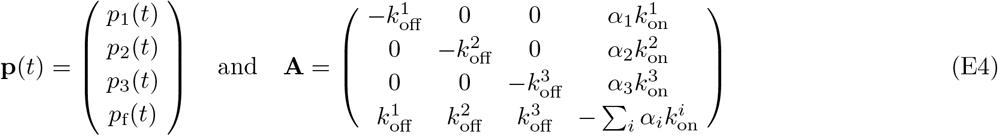

In the above Eq. (E3) and Eq. (E4), **u** = (1 1 1 1). *α*_*i*_ is the fraction of freely diffusing fluorescent histone dimers transported by chaperones in the *i*^th^ nucleus region (*i* = 1, 2, 3). 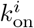 and 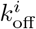 are the average binding and dissociation rates of histone dimers to / from DNA in the corresponding regions of the cell nucleus.

To fit the experimentally measured normalized FRAP signal [Eq. (E1)], the steady-state of Eq. (E3) describing the state of the nucleus before photobleaching was first obtained by solving the following linear system:

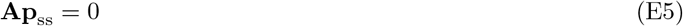

Where **p**_ss_ is the steady-state probability vector.

Knowing the vector **p**_ss_, the value of the probability vector immediately after the photobleaching pulse can be written as:

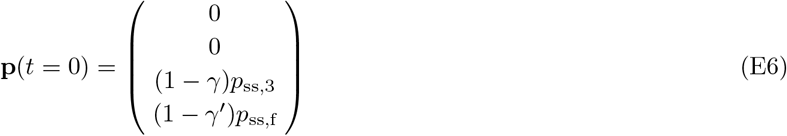

Where *p*_ss,3_ and *p*_ss,f_ are the corresponding elements of the steady-state probability vector. *γ* is the fraction of the photobleached nucleus area outside the dense and sparse chromatin regions, whereas *γ*^*′*^ is the fraction of the photobleached freely diffusing histone dimers. The above formula takes into account that DNA-bound histones in the dense and sparse chromatin regions are almost completely photobleached, as found in experiments carried out on fixed cells, see Figures 4(b) and S9.

To calculate fluorescence recovery curves as a function of time for the dense and sparse chromatin regions, Eq. (E3) was solved using the initial condition shown in Eq. (E6):

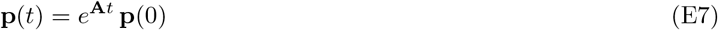

The normalized fluorescence recovery curve for each chromatin region was then calculated using the following formula:

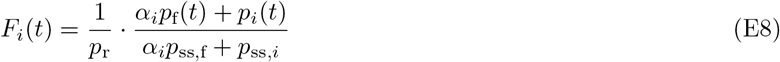

Where *p*_r_ = 1 − *p*_ss,1_ − *p*_ss,2_ − *γp*_ss,3_ − *γ*^*′*^*p*_ss,f_ is the fraction of fluorescent histone dimers remaining after the photo-bleaching.

It should be noted that the values of many of the model parameters shown in Eq. (E3), Eq. (E4) and Eq. (E6) can be found directly from experimental measurement, which significantly reduces the model degrees of freedom and makes the data fitting more robust. In particular, the fraction of freely diffusing histone dimers (*α*_*i*_) in each region of the nucleus can be estimated using the third line in Eq. (E3) in combination with the following two equations:

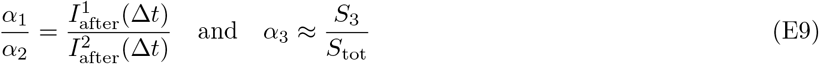

Where 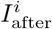 is the total fluorescence intensity of the *i*^th^ region of the nucleus (*i* = 1, 2) after the photobleaching pulse. Δ*t* is the average time for the initial establishment of quasi-equilibrium of freely diffusing fluorescent histone dimers between the unbleached and photobleached regions of the nucleus (Δ*t* ≈ 1 − 2 min, see Figures 4(b), S9 and S11). *S*_*i*_ and *S*_tot_ are the area of the *i*^th^ region of the nucleus (*i* = 1, 2, 3) and the total projected area of the nucleus (*S*_tot_ = *S*_1_ + *S*_2_ + *S*_3_), respectively (see Figure S13).

Similarly, *γ, γ*^*′*^ and *p*_ss,f_ can be determined experimentally using the following formulas:

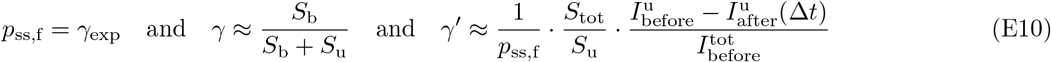

Here *γ*_exp_ is the average fraction of freely diffusing histone dimers in the nucleus, measured using the continuous photobleaching method [191], see Figure S14. *S*_b_ and *S*_u_ are the photobleached and unbleached areas outside the dense and sparse chromatin regions (*S*_b_ + *S*_u_ = *S*_3_, Figure S13). 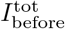 is the integral fluorescence intensity of the entire nucleus before the photobleaching. 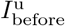 and 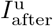 are the integral fluorescence intensities of the unbleached part of the nucleus before and after the photobleaching pulse.

Therefore, to fit the normalized FRAP signals measured in three different regions of the nucleus to Eq. (E8), it is necessary to vary only the kinetic rates 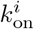 and 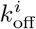 (*i* = 1, 2, 3) such that the model obeys the following three experimental constraints:

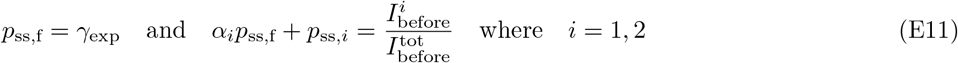

Here 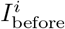 is the experimentally measured integral fluorescence intensity of the *i*^th^ region of the nucleus before the photobleaching. These constraints were imposed on the model in the form of a penalty function that had to be minimized in addition to minimizing the deviation of the theoretically predicted FRAP curves from the measured experimental data points. Minimization was performed using the Nelder-Mead simplex algorithm [192].

As a result, the model had only three degrees of freedom, making the simultaneous fitting of the normalized FRAP signals measured in the dense and sparse chromatin domains described by Eq. (E1) very robust (i.e., 1.5 degrees of freedom per fluorescence recovery curve).

## Appendix F: Correlation between the density of freely diffusing histone-chaperone complexes, nuclear electrostatic potential and chromatin density

FRAP experiments performed in our study show that the fluorescent signal of electrically charged mEGFP WT probes is recovered in the photobleached area much faster compared to the case of freely diffusing histone-chaperone complexes: the fluorescent mEGFP WT signal in the photobleached area becomes indistinguishable from the signal in the rest of the nucleus in *<* 15 s [Figure S15], whereas the signal recovery of fluorescently-labelled histones due to their diffusion takes ∼ 1 − 2 min [Figures 4(b), S9 and S15]. These results indicate that the diffusion of histone-chaperone complexes occurs significantly slower, which cannot be explained only by the difference in the size of mEGFP and histone-chaperone complexes. Thus, it can be concluded that freely diffusing histone-chaperone complexes are involved in interactions with the surrounding microenvironment.

To describe the experimentally observed correlation between the density of freely diffusing histone-chaperone complexes, nuclear electrostatic potential, and chromatin density, we used a model that considers two chromatin regions, dense and sparse, which are designated by indexes *i* = 1 and *i* = 2, respectively. All histone-chaperone complexes are divided into two groups: 1) those that move freely, and 2) those that are transiently bound to surrounding weak binding sites. In both chromatin regions (*i* = 1, 2), a quasi-equilibrium between the two groups of histone-chaperone complexes is assumed:

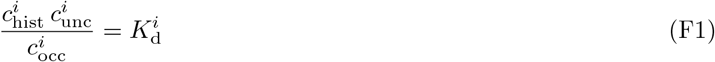

Where 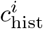 is the concentration of freely moving histone-chaperone complexes in the *i*^th^ chromatin region (*i* = 1, 2) that are not associated with weak binding sites. 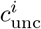 and 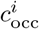 are the concentrations of unoccupied and occupied weak binding sites in the *i*^th^ chromatin region. 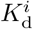 is the dissociation constant in the corresponding chromatin region.

The freely diffusing fraction of histones measured in FRAP experiments includes both freely moving histone-chaperone complexes and those transiently bound to weak binding sites: 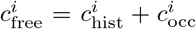, since these two groups exchange with each other over a timescale (seconds to minutes) that is shorter than the timescale associated with the exchange of histones between nucleosomes and histone-chaperone complexes (on the order of hours), see Figures 4(b), S9 and S15. Since FRAP experiments suggest that the fraction of histone-chaperone complexes bound to weak binding sites is significantly larger than the fraction of freely moving histone-chaperone complexes (otherwise, these complexes would rapidly reach equilibrium between the photobleached region and the rest of the nucleus within a few seconds, which is not observed in the experiments), it follows that 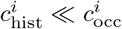 and 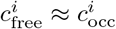.

Furthermore, since interactions between histone-chaperone complexes and binding sites are assumed to be weak, the binding free energy of such interactions (*µ*_*i*_), which is determined by 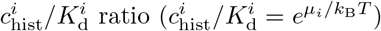, should be low (i.e., *µ*_*i*_ *<* 0 and, therefore, 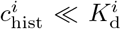). In this case, it follows from Eq. (F1) that 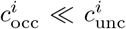 and, thus, 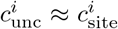, where 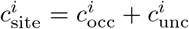 is the total concentration of weak binding sites in the corresponding chromatin region. Using Eq. (F1), one can then find that the ratio between the fluorescent signals corresponding to the freely diffusing fraction of histones measured in the dense and sparse chromatin regions in the FRAP experiments is:

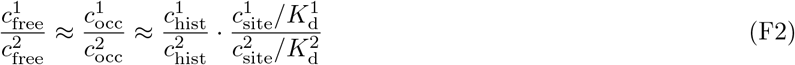

In equilibrium (or quasi-equilibrium), the densities of mEGFP WT probes and freely moving histone-chaperone complexes obey Boltzmann’s law given by Eq. (B1), from which it follows that:

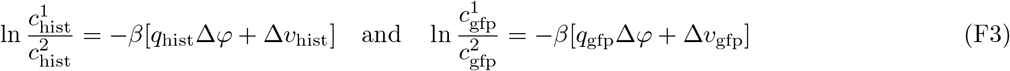

Where *q*_hist_ and *q*_gfp_ are the electrical charges of the histone-chaperone complexes and mEGFP WT probes, respectively. Δ*φ* = *φ*_1_ − *φ*_2_ is the difference in electrostatic potentials of the corresponding chromatin regions, and Δ*v*_hist_ and Δ*v*_gfp_ are the analogous differences in the volume-exclusion energies of the corresponding protein complexes between the two chromatin regions. Here we assume that freely moving protein complexes participate only in electrostatic and volume-exclusion interactions with the surrounding microenvironment in the nucleus, omitting energy terms describing other types of interactions, for details see Appendix (B).

Using a formula for histone-chaperone complexes similar to Eq. (C6), it can be shown that in the physiologically relevant case of Δ*v*_gfp_ ≲ 0.3 *k*_B_*T*, the approximate relationship Δ*v*_hist_ ≈ *α*Δ*v*_gfp_ holds, where the proportionality coefficient *α* depends on the characteristic size of the histone-chaperone complexes. Substituting this formula into Eq. (F3), it is easy to see that:

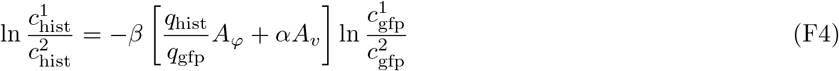

Where *A*_*φ*_ and *A*_*v*_ are the linear slopes of the calibration curves of the electrostatic and volume-exclusion interaction energies of mEGFP WT probes shown in Figure S6(c,d).

After substituting Eq. (F4) into Eq. (F2), we get:

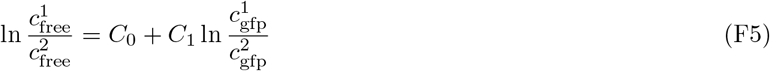

Here 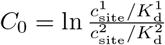 and 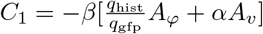.

Therefore, by experimentally measuring the correlation between the logarithms of the ratios 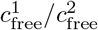 and 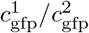 and using Eq. (F5) to fit the experimental data, it is possible to determine the net electrical charge of histone-chaperone complexes and the strength of their transient binding interactions with the surrounding microenvironment in the nuclei of living cells. I.e., mEGFP WT probes can be used as a reference for experimental measurements of key physical parameters determining the nucleoplasmic distribution of histone-chaperone complexes.

In fact, by measuring the chromatin density ratio 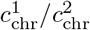 in addition to the above ratios of the freely diffusing fraction of histones and mEGFP WT probes, one can gain information about the potential correlation between the density of weak binding sites, 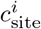, and the chromatin density, 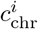. In particular, assuming that 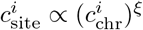, where *ξ* is the coefficient describing the cooperativity of the formation of weak binding sites by chromatin, from Eq. (F2) and Eq. (F5) it is easy to obtain the following formula:

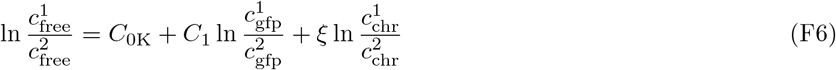

Where *C*_0K_ = ln 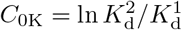.

## Appendix G: Effect of nuclear electrostatic potential on the stability of nucleosomes, hexasomes and tetrasomes

In general, the stability of nucleoprotein complexes such as nucleosomes, hexasomes and tetrasomes is determined by the binding free energy of the corresponding protein complexes to DNA (i.e., the binding free energy of histone octamers / hexamers / tetramers to DNA). To calculate this energy, it should be noted that histone-based nucleoprotein complexes consist of several H2A · H2B and H3 · H4 histone dimers, which are typically transported by two different groups of chaperones [3, 4] that we will denote as c_1_ and c_2_. As a result, the overall chemical equation describing nucleosome assembly can be schematically represented as shown in Figure S16, see also ref. [66, 193].

From Figure S16 it follows that the binding free energy of histone octamers / hexamers / tetramers to DNA generally depends on the chemical potentials of the chaperones in their histone-bound (c_1_b, c_2_b) and unloaded states (c_1_u, c_2_u), i.e., on the chemical potentials of the initial reactant complexes and the final products. These chemical potentials can be found by calculating the partition function (*Z*_*i*_) of each type of chaperone complex involved in the chemical equation shown in Figure S16:

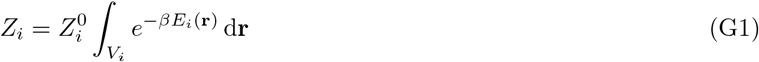

Here the subscript *i* indicates the different types of chaperone complexes: *i* = c_1_u, c_2_u, c_1_b and c_2_b are histone-binding chaperones in the unloaded (c_1_u, c_2_u) and histone dimer-bound states (c_1_b, c_2_b), respectively. 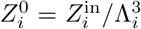, whereΛ_*i*_ and 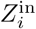 are the de Broglie thermal wavelength and the part of the partition function describing the inner degrees of freedom of the *i*^th^ type of chaperone complexes. *E*_*i*_(**r**) is the location-dependent energy of the *i*^th^ type of chaperone complexes. *β* = 1*/k*_B_*T* is the reciprocal of the thermodynamic temperature. *V*_*i*_ is the total volume available for diffusion of the *i*^th^ type of chaperone complexes.

Generally, histone-binding chaperones can shuttle between the cell cytosol and nucleoplasm, accessing the entire osmotically active volume of the cell 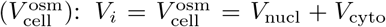, where *V*_nucl_ and *V*_cyto_ are the volumes of the nucleus and cytosol, respectively. However, due to the presence of active nuclear transport through nuclear pore complexes, the cytosolic and nucleoplasmic concentrations of chaperones usually differ. For example, experimental measurements of the average cytosolic and nucleoplasmic densities of freely diffusing histone-chaperone complexes in HEK293T cells performed in our study show that the nuclear-to-cytoplasmic ratio (N/C ratio) of freely diffusing H2A · H2B histone dimers is 4.40 ± 0.24 (mean ± SEM, *N* = 21 cells), whereas in the case of H3 · H4 dimers it is 5.10 ± 0.32 (mean ± SEM, *N* = 22 cells), which is in good agreement with previously published measurements performed in HeLa cells for H2B-GFP histones (N/C = 8.5 ± 3.3 [182]).

Given that freely diffusing histone-chaperone complexes rapidly reach equilibrium / quasi-equilibrium in both the cytosol and nucleoplasm [the characteristic fluorescence recovery time due to diffusion of histone-chaperone complexes in the nucleus is ∼ 1 min, see Figures 4(b), S9 and S15], it follows that the net effect of active nuclear transport can be simply described by the free energy offset, *η*_*i*_, gained by chaperone complexes located in the nucleus due to active transport across the NE, see [25]. As a result, for chaperones in the histone-bound and unloaded states, we have:

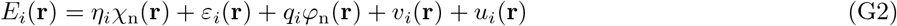

Here *q*_*i*_ is the net electrical charge of the *i*^th^ type of chaperone complexes (*i* = c_1_u, c_2_u, c_1_b and c_2_b). *η*_*i*_ is the change in the free energy of the *i*^th^ type of chaperone complexes due to their translocation into the nucleus via active nuclear transport. *χ*_n_(**r**) is the characteristic function of the nucleus: *χ*_n_(**r**) = 1 if **r** ∈ *V*_nucl_, and 0 otherwise. *ε*_*i*_(**r**) and *v*_*i*_(**r**) are the energies of weak transient binding interactions and volume-exclusion interactions of chaperone complexes with the surrounding microenvironment in the nucleus, see Appendix F. *u*_*i*_(**r**) is the energy term accounting for other types of interactions of chaperone complexes, see Appendix B.

Knowing the partition function of the chaperone complexes, one can then calculate their electrochemical potentials using the definition of Gibbs free energy:

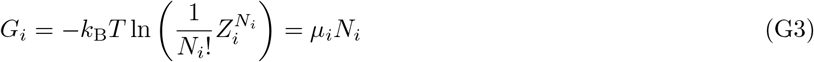

Here *G*_*i*_ is the part of the Gibbs free energy contributed by the *i*^th^ type of chaperone complexes. *µ*_*i*_ and *N*_*i*_ are the electrochemical potential and the total number of molecules of the *i*^th^ type of chaperone complexes.

Using Stirling’s formula, ln *N* ! ≈ *N* ln *N* − *N*, from Eq. (G3) it follows that:

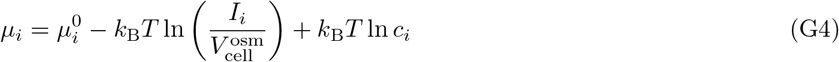

Here 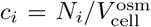 is the average concentration of the *i*^th^ type of chaperone complexes. 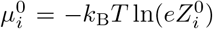 is the standard chemical potential of the *i*^th^ type of chaperone complexes, and *I*_*i*_ is the following integral:

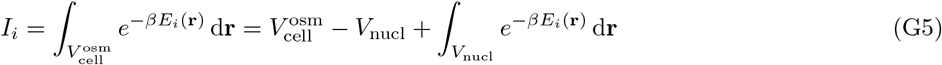

Importantly, from Eq. (G1), Eq. (G2) and Boltzmann’s law it follows that the N/C ratio of chaperone complexes is equal to:

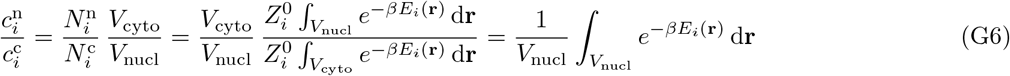

Where 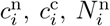 and 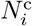 are the average concentrations and total amounts of the *i*^th^ type of chaperone complexes (*i* = c_1_u, c_2_u, c_1_b, c_2_b) in the nucleus and cytosol, respectively.

Substituting Eg. (G6) into Eq. (G5), it is easy to see that:

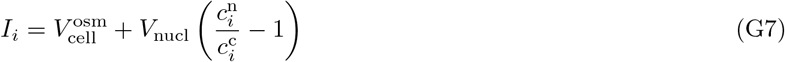

I.e., the values of the integrals *I*_*i*_ can be determined by measuring cell and nuclear volumes, as well as the N/C ratio of the corresponding chaperone complexes.

Given electrochemical potentials of the chaperone complexes, the binding free energy of a histone octamer to DNA at the location specified by the position vector **r** can be found as the difference between the sum of the electrochemical potentials of the initial reactants and the final products shown in Figure S16:

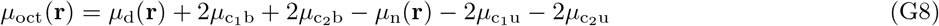

Here *µ*_oct_ is the binding free energy of histone octamers to DNA. *µ*_d_(**r**) and *µ*_n_(**r**) are the location-dependent electrochemical potentials of the binding site on DNA in the unoccupied and histone octamer-bound states, respectively:

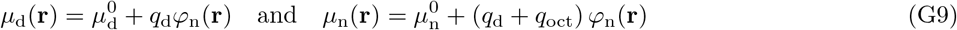

Where 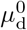 and 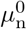 are the standard chemical potentials of the binding site in the unoccupied and histone octamer-bound states, respectively. *q*_d_ and *q*_oct_ are the net electrical charges of the binding site and the histone octamer, correspondingly. Here we neglect the energy of volume-exclusion interactions of nucleosomes with the surrounding microenvironment, since it is estimated to be several times smaller than the energy of electrostatic interactions, see Appendix C.

Substituting Eq. (G4) and Eq. (G9) into Eq. (G8), it follows that:

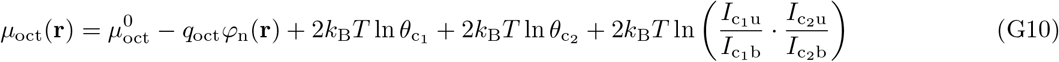

Here 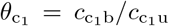 and 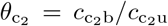 are the occupancy ratios of chaperones c_1_ and c_2_, respectively. 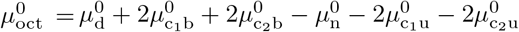 is the standard Gibbs free energy of nucleosome assembly in the absence of nuclear electrostatic potential. The average experimentally measured value of 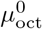 is of the order of ∼ 7 *k*_B_*T* [66, 194].

Using the approach described above, it is also easy to derive formulas similar to Eq. (G10) in the case of histone tetramers and hexamers. Namely, for the binding free energies of histone hexamers (*µ*_hex_) and tetramers (*µ*_tet_) to DNA we have:

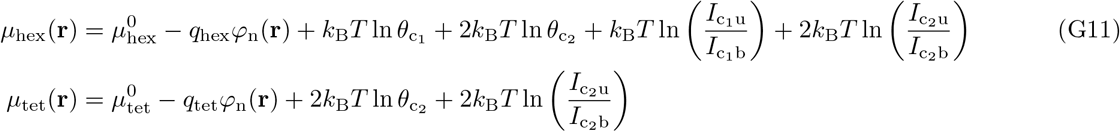

Where 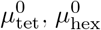, *q*_tet_ and *q*_hex_ are the standard Gibbs free energies and electrical charges of the corresponding protein complexes.

From Eq. (G10) and Eq. (G11) it follows that the combined contribution of the nuclear electrostatic potential, as well as weak transient binding interactions and active nuclear transport of chaperone complexes to the binding free energies of histone octamers / hexamers / tetramers to DNA at a given location (**r**) is described by the following formulas:

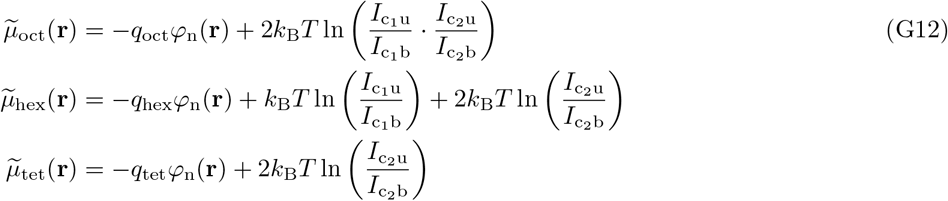

Thus, from Eq. (G12) it is clear that the nuclear electrostatic potential has not only a local effect on the binding free energy of nucleoprotein complexes [− *q*_*i*_*φ*_n_(**r**) energy terms], but also a global effect on the nuclear-cytoplasmic distributions of chaperone complexes, which leads to a shift in the binding free energies of histone octamers / hexamers / tetramers to DNA [the remaining energy terms in Eq. (G12)].

Using Eq. (G12), one can estimate statistically-weighted binding free energies of nucleosomes, hexasomes and tetrasomes in a given region of the nucleus based on experimental measurements as:

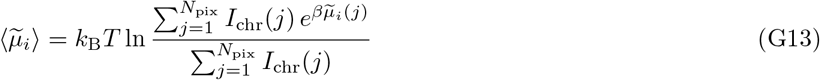

Where the sum is taken over the pixels of the given region in an experimentally obtained image of the nucleus. *N*_pix_ is the total numbers of pixels in the corresponding nuclear region. *I*_chr_(*j*) is the chromatin intensity at the *j*^th^ pixel, measured by expressing fluorescently-labelled histones, whereas *µ*_*i*_(*j*) is the binding free energy of the corresponding protein complex (*i* = oct, hex, tet) at the same pixel, estimated based on nuclear electrostatic potential measurements using mEGFP probes coexpressed in the same cell.

Using Eq. (G12) and Eq. (G13), we calculated statistically-weighted binding free energies of nucleosomes, hexasomes and tetrasomes in nuclear regions of low and high electrostatic potential, which are often associated with dense and sparse chromatin regions in HEK293T cells grown on fibronectin-coated glass slides, by selecting nuclear regions (except nucleoli) that contain the ∼ 16% of the brightest and faintest pixels of the nuclei images in GFP channel (i.e., selecting pixels with the intensities greater than the mean + SD and less than the mean − SD). Table T3 shows the calculated values of the binding free energies of corresponding nucleoprotein complexes, averaged over *N* = 21 HEK293T cells grown on fibronectin-coated glass slides.

## Appendix H: Transfer-matrix calculations

To assess the effect of the interaction of nucleoprotein complexes with the nuclear electrostatic potential on their stability, we used the transfer-matrix calculation method described previously in ref. [25, 51–53]. In particular, in the calculations we considered a small part of chromatin located either in a euchromatic or heterochromatic region in the presence of force and torque constraints applied to DNA. On the one hand, we used these mechanical constraints to characterize the stability of nucleoprotein complexes by calculating the characteristic unfolding force of nucleoprotein complexes in different microenvironments (euchromatin / heterochromatin), – i.e., more stable nucleoprotein complexes can withstand higher stretching forces before experiencing unfolding / protein dissociation from DNA. On the other hand, these mechanical constraints were also used in our study to assess the potential influence of RNA polymerases and other DNA-motors on local chromatin structure, since RNA polymerases are known to exert very high forces and torques on DNA during gene transcription [195, 196].

Following ref. [25, 51–53], the behaviour of DNA under force and torque constraints was described using a semi-flexible polymer model, in which DNA is represented by a polygonal chain consisting of straight segments whose 3D orientations in space are characterized by three Euler rotation angles. By introducing transfer-matrices defined at each of the vertices joining neighbouring DNA segments, it is then possible to calculate the DNA partition function and obtain detailed information regarding the DNA conformation and DNA structural fluctuations. Namely, the DNA partition function (*Z*_*f,τ*_) under force *f* and torque *τ* can be found as:

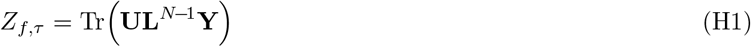

Here **L** is the DNA transfer-matrix, which describes the structural state of each DNA segment under the force and torque constraints, and the matrices **Y** and **U** describe the boundary conditions imposed on the DNA end segments. *N* is the total number of DNA segments considered in the calculations.

While detailed information on the structure and elements of the transfer-matrix **L** and the boundary condition matrices **Y** and **U** can be found in ref. [53], here we would like only to mention that in the present study we considered four possible structural states of DNA that are the most relevant to physiological ranges of forces and torques: 1) B-DNA form, which is a right-handed DNA most frequently found in living cells in the case of relaxed or slightly supercoiled DNA; 2) L-DNA form, a left-handed DNA, which is more energetically favourable at fairly strong negative torques; 3) P-DNA form, a right-handed DNA, which is energetically favourable at strong positive torques, and 4) S-DNA form, an overstretched extended DNA, which is observed when DNA subjected to large stretching forces, see ref. [51, 52] for a more detailed description of these DNA forms. The values of the model parameters used in the transfer-matrix calculations to describe different DNA structural states can be found in Table T1. In addition to these parameters, it should be noted that in all our calculations the size of all DNA segments was set to *b* = 0.5 nm (i.e., *b*_bp_ = 1.5 bp) for all DNA structures.

**TABLE T1.** Structural and elastic characteristics of different DNA forms^(1,2)^.

| DNA form | Bending persistence length | Twisting persistence length | Contour length relative to B-DNA | DNA helical repeat | Base-pairing energy relative to B-DNA | $\lambda$ |
| --- | --- | --- | --- | --- | --- | --- |
| B-DNA | 50 nm<br>[202, 203] | 95 nm<br>[204–206] | 1 | 10.4 bp [207] | 0 | 4.3 [51–53] |
| L-DNA | 7 nm [208, 209] | 15 nm<br>[208–210] | 1.35 [208, 209] | 16 bp [204, 208, 210] | 5 $k_B T$ [51–53] | 4.3 [51–53] |
| P-DNA | 15 nm [209] | 25 nm [209] | 1.7 [209, 211, 212] | 3 bp<br>[204, 209, 211–213] | 17.8 $k_B T$ [51–53] | –1.0 |
| S-DNA | 15 nm<br>[209, 214] | 20 nm [209] | 1.7 [204, 212, 215] | 35 bp [204, 212] | 5.1 $k_B T$ [52] | 4.3 [52] |

<sup>1</sup> To account for the cooperativity of the DNA structural transitions, the domain wall penalty, $J = 9.0 k_B T$ [210, 216], characterizing the DNA preference for structural uniformity was used in the transfer-matrix calculations in addition to the model parameters listed in the above table.
<sup>2</sup> For more information about the model parameters, see ref. [51–53].

Beside various structural states of DNA, the calculations also took into account the formation of nucleoprotein complexes on B-DNA, including L- and R-tetrasomes (left- and right-handed tetrasomes), hexasomes, and nucleosomes, which are formed by histone dimers. Furthermore, we also considered the formation of DNA-transcription factor complexes to assess the potential effect of the local microenvironment (euchromatin / heterochromatin) on the DNA-binding competition between histone-based nucleoprotein complexes and transcription factors. The structures of the histone-based nucleoprotein complexes used in the calculations were the same as those described in ref. [53]. As for the transcription factors, they were assumed to belong to the family of DNA-bending proteins [53], forming nucleoprotein complexes with an architecture similar to that experimentally observed for TATA-binding protein [197] or Sox2 [198] transcription factors. The values of the model parameters used in the transfer-matrix calculations to describe the architecture of nucleoprotein complexes can be found in Table T2. In addition, the binding free energies of various histone-based protein complexes to DNA in euchromatic and heterochromatic regions, estimated from experimental measurements using Eq. (G12), are listed in Table T3. As for the binding free energy of transcription factors to DNA, it was set equal to 2.3 k_B_T, regardless of the local microenvironment (i.e., it was assumed that the local concentration of transcription factors in both euchromatic and heterochromatic regions is ∼ 10 times higher than the equilibrium dissociation constant of the transcription factor from DNA, see Figure 5 captions for details).

**TABLE T2.** Values of the model parameters for different nucleoprotein complexes used in the transfer-matrix calculations^(1,4)^.

| Nucleoprotein complex | Standard Gibbs free energy of assembly<sup>(2)</sup> | DNA linking number change | Rotation matrices of the entry and exit DNA segments, $\mathbf{A}_{\text{in}}$ and $\mathbf{A}_{\text{out}}$ |
| --- | --- | --- | --- |
| L-tetrasome | 2.11 k<sub>B</sub>T [66] | −0.73 [217] | $\mathbf{A}_{\text{in}}(0, 2.26, -1.11)$ [53]<br>$\mathbf{A}_{\text{out}}(-1.11, 2.26, 0)$ [53] |
| R-tetrasome | −0.19 k<sub>B</sub>T<sup>(3)</sup> [66, 217] | +1.0 [217] | $\mathbf{A}_{\text{in}}(0, 2.26, 1.11)$ [53]<br>$\mathbf{A}_{\text{out}}(1.11, 2.26, 0)$ [53] |
| Hexasome | 4.55 k<sub>B</sub>T [66] | −0.97 | $\mathbf{A}_{\text{in}}(0, 1.07, 0.12)$<br>$\mathbf{A}_{\text{out}}(0.12, 1.07, 0)$ |
| Nucleosome | 6.98 k<sub>B</sub>T [66] | −1.2 [217, 218] | $\mathbf{A}_{\text{in}}(0, 2.12, -0.79)$ [53]<br>$\mathbf{A}_{\text{out}}(-0.79, 2.12, 0)$ [53] |
| DNA-bending transcription factor | | 0 | $\mathbf{A}_{\text{in}} = \mathbf{A}_{\text{out}} = \mathbf{A}_1 = \mathbf{I}$ [53]<br>$\mathbf{A}_2(\pi, 0.14, \pi)$ [53] |

<sup>1</sup> In addition to the model parameters listed in the table, all nucleoprotein complexes were characterized by dimensionless bending and twisting elasticities of the entry and exit DNA segments of the nucleoprotein complexes: $a_{\text{pr}} = 33$ and $c_{\text{pr}} = 33$ , see ref. [53]. Furthermore, the occupancy ratios of histone-binding chaperones were set to $\theta_{c_1} = \theta_{c_2} = 6$ so that the characteristic stretching force leading to the destabilization of histone-based nucleoprotein complexes in heterochromatin domains would be on the order of $\sim 1.5 - 2$ pN, see the main text. Notably, from Eq. (G12) it follows that the difference between the Gibbs free energies of nucleoprotein complexes ( $\Delta\tilde{\mu}_i = \tilde{\mu}_i^e - \tilde{\mu}_i^h$ ) in euchromatic ( $\tilde{\mu}_i^e$ ) and heterochromatic ( $\tilde{\mu}_i^h$ ) regions for each type of nucleoprotein complex ( $i = \text{oct, hex, tet}$ ) does not depend on the exact values of $\theta_{c_1}$ and $\theta_{c_2}$ .
<sup>2</sup> This is the standard Gibbs free energy of the assembly reaction of the corresponding nucleoprotein complexes in the presence of histone-binding chaperones, measured in experiments [66].
<sup>3</sup> The standard Gibbs free energy of R-tetrasome assembly was estimated based on experimental measurements the standard Gibbs free energy of L-tetrasome assembly [66] and the difference in free energies of L- and R-tetrasomes ( $\Delta G = 2.3$ k<sub>B</sub>T [217]).
<sup>4</sup> For more information about the model parameters, see ref. [53].

**TABLE T3.** Contribution of nuclear electrostatic potential, weak transient interactions and active nuclear transport to the Gibbs free energy of histone-based nucleoprotein complexes in nuclear regions of low (heterochromatin) and high (euchromatin) nuclear electrostatic potentials, estimated based on experimental measurements [see Eq. (G12) and Eq. (G13) in Appendix G]^(1)^.

| Nucleoprotein complex | Euchromatin,<br>contribution to the<br>Gibbs free energy, $\tilde{\mu}_i$ | Heterochromatin,<br>contribution to the<br>Gibbs free energy, $\tilde{\mu}_i$ |
| --- | --- | --- |
| L- and R-tetrasomes | -2.03 k<sub>B</sub>T | 1.15 k<sub>B</sub>T |
| Hexasome | -2.86 k<sub>B</sub>T | 2.12 k<sub>B</sub>T |
| Nucleosome | -3.68 k<sub>B</sub>T | 3.38 k<sub>B</sub>T |

<sup>1</sup> For simplicity, in the calculations it was assumed that the N/C ratios of chaperones 1 and 2 in the unloaded state is: $c_{c1u}^n/c_{c1u}^c = c_{c2u}^n/c_{c2u}^c = 2$ . Notably, from Eq. (G12) it follows that the difference between the Gibbs free energies of nucleoprotein complexes ( $\Delta\tilde{\mu}_i = \tilde{\mu}_i^e - \tilde{\mu}_i^h$ ) in euchromatic ( $\tilde{\mu}_i^e$ ) and heterochromatic ( $\tilde{\mu}_i^h$ ) regions for each type of nucleoprotein complex ( $i = \text{oct, hex, tet}$ ) does not depend on the exact values of the N/C ratios of chaperones 1 and 2.

Given the DNA partition function, *Z*_*f,τ*_, one can calculate observable quantities commonly measured in experiments, such as DNA extension (*z*) and linking number change (ΔLk), as well as the total number of DNA segments (*N*_pr,*i*_) bound to the *i*^th^ type of protein (*i* = L/R-histone tetramer, hexamer, octamer, or transcription factor) and the total number of DNA segments (*N*_dna,*j*_) in each of the alternative structural states of DNA (*j* = L-, P- or S-DNA), by differentiating *Z*_*f,τ*_ with respect to the force (*f*), torque (*τ*), protein binding free energy (*µ*_pr,*i*_), or DNA base-pairing energy (*µ*_dna,*j*_) in the corresponding state [53]:

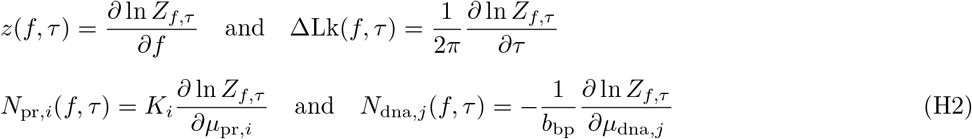

Here *K*_*i*_ is the total number of DNA segments occupied by each nucleoprotein complex of type *i*.

Using the above observables, it is then easy to find the fraction of DNA occupied by DNA-bound proteins (*O*_pr,*i*_) or the fraction of DNA in an alternative DNA structural state (*O*_dna,*j*_) as: *O*_pr,*i*_ = *N*_pr,*i*_*/N* and *O*_dna,*j*_ = *N*_dna,*j*_*/N* . This allows one to plot the stability phase diagrams of DNA / chromatin shown in Figures 5(c-e) and S18. The solid lines in these diagrams show the transition boundaries between different structural states of DNA and/or DNA-protein complexes, which were calculated as follows. The boundaries between B-, L-, P- and S-DNA states were defined as the set of points (*f, τ*) at which ∼ 50% of the DNA segments are in L-, P- or S-DNA forms, respectively. Furthermore, the boundary between the extended and supercoiled conformations of DNA in a particular structural state (scB-, scL- and scP-DNA) was defined as the set of points at which DNA extension experiences a ∼ 50% drop with respect to the value predicted by the worm-like chain model for the corresponding form of DNA in the torsionally relaxed state. As for the transition boundaries between the bare B-DNA and protein-bound DNA states, they were defined as the set of points at which half of the maximum DNA binding sites is occupied by the corresponding protein. Here we would like to point that the total number of DNA binding sites is not necessarily equivalent to the total number of DNA segments. The main reason for this is the existence of bare DNA segment gaps between nucleoprotein complexes, which correspond to DNA linkers connecting adjacent DNA-protein complexes. For example, in the case of reconstituted nucleosome arrays or tightly packed yeast chromatin, the minimal length of such DNA linkers was found to be of the order of ∼ 10 − 20 bp [199, 200]. For this reason, the minimal possible spacing between adjacent nucleosomes was set to 18 bp (i.e., 12 DNA segments) in all transfer-matrix calculations. The same minimum DNA linker length was also used in calculations of DNA interactions with histone tetramers, hexamers, and transcription factors, since previously published structural data suggest that such linkers are also likely to exist between DNA-bending nucleoprotein complexes, see, for example, ref. [201].

**FIG. S1.**
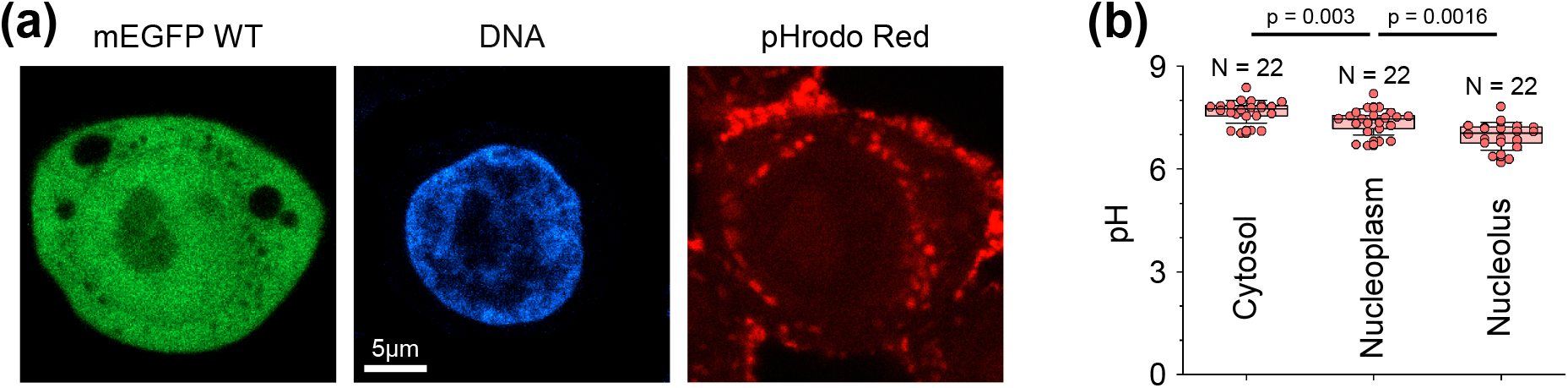
Measurement of the pH level in HEK293T cells. **(a)** Representative image of a HEK293T cell expressing mEGFP WT, additionally stained with Hoechest and pHrodo Red. **(b)** Average pH levels of the cytoplasm, nucleus and nucleoli of HEK293T cells measured using pHrodo Red. A total of *N* = 22 cells were measured. Boxes and error bars represent the interquartile range and SD, respectively. Bars above the graph show pairwise statistical differences between corresponding experimental data, calculated using Mann-Whitney test.

**FIG. S2.**
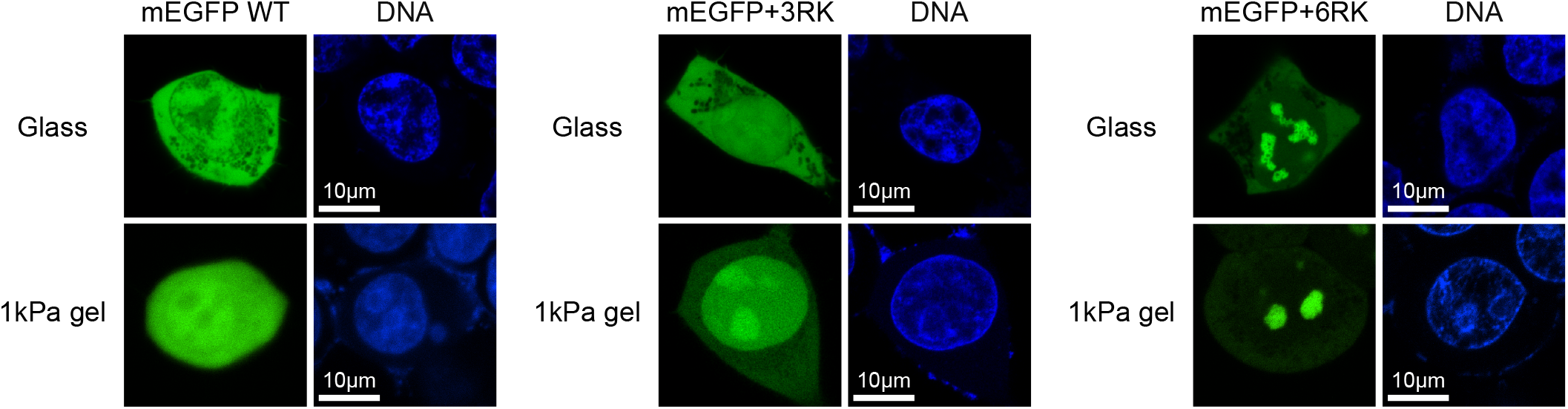
Distribution of different mEGFP probes in HEK293T cells. Representative images showing the nuclear-cytoplasmic distribution of mEGFP WT, mEGFP+3RK and mEGFP+6RK probes and DNA in HEK293T cells grown on fibronectin-coated glass slides and 1 kPa gel.

**FIG. S3.**
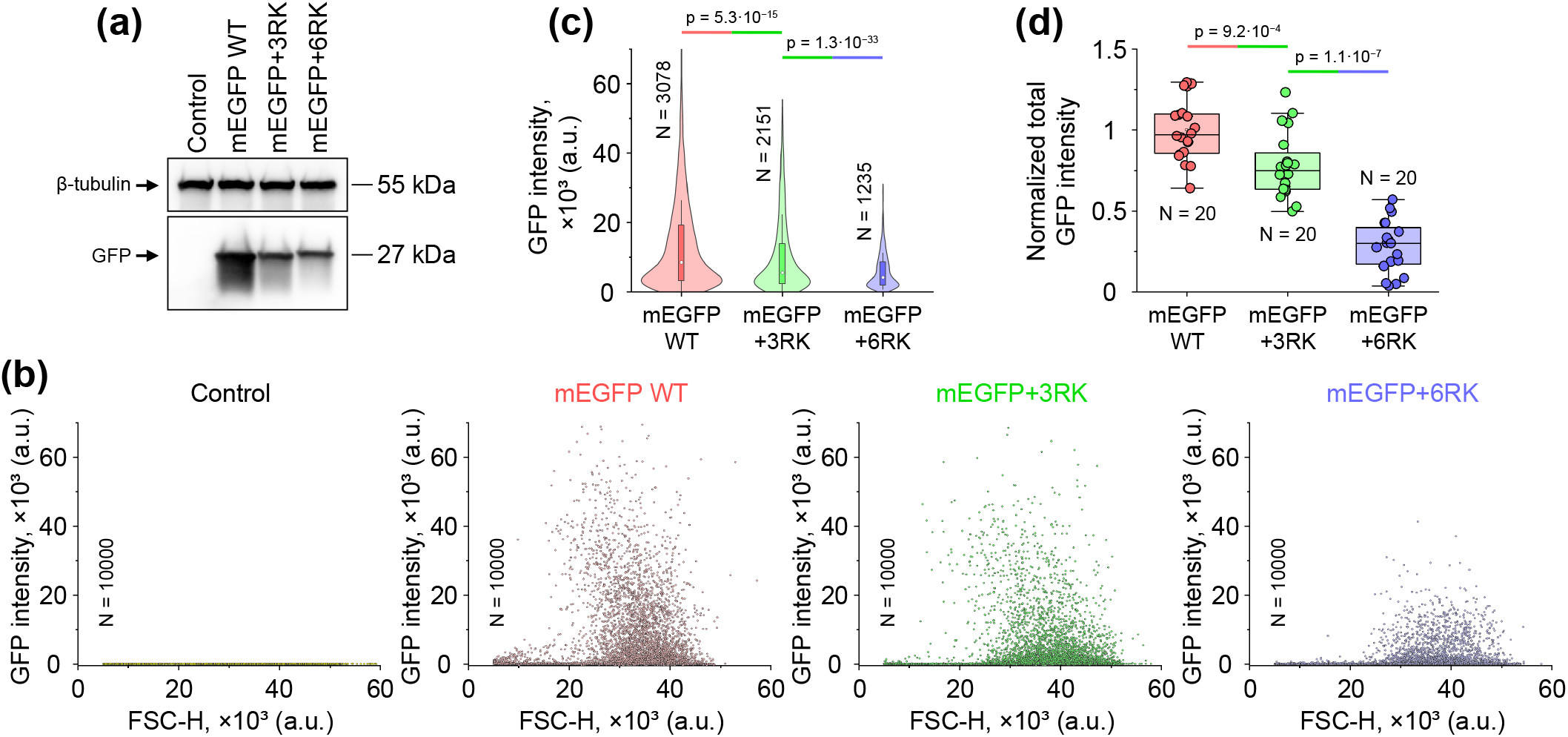
Characterization of expression levels of different mEGFP probes in HEK293T cells. **(a, b)** Western blot (a) and FACS plots (b) of HEK293T cells expressing mEGFP WT, mEGFP+3RK and mEGFP+6RK probes. The control group includes WT HEK293T cells that were not transfected with any of the mEGFP constructs. **(c)** Total fluorescence intensity of HEK293T cells expressing mEGFP WT, mEGFP+3RK and mEGFP+6RK probes in the GFP channel obtained from the FACS plots shown in panel (b). The graph shows distributions calculated considering only cells in which the GFP signal exceeds the background level measured in the control case. **(d)** Normalized total fluorescence intensity of HEK293T cells expressing mEGFP WT, mEGFP+3RK and mEGFP+6RK probes measured in independent experiments using a confocal microscope. Normalization was performed to the mean total fluorescent intensity of HEK293T cells expressing mEGFP WT. Comparison of the data presented in panels (a, c, d) shows that all methods provide consistent estimates of mEGFP probe expression levels in HEK293T cells. Overall, the results presented in panels (a, c, d) and in Figure S4 suggest that the aggregation of mEGFP+6RK probes in HEK293T cell nuclei, which can be seen in Figure S2, is unlikely to be caused by a difference in their expression level compared to mEGFP WT or mEGFP+3RK probes, but rather by a difference in their interaction with nuclear components. In panels (c, d), *N* indicates the number of cells processed under the corresponding experimental conditions. Boxes represent the interquartile range, error bars indicate SD. Bars above the graph show pairwise statistical differences between corresponding experimental data, calculated using Mann-Whitney test.

**FIG. S4.**
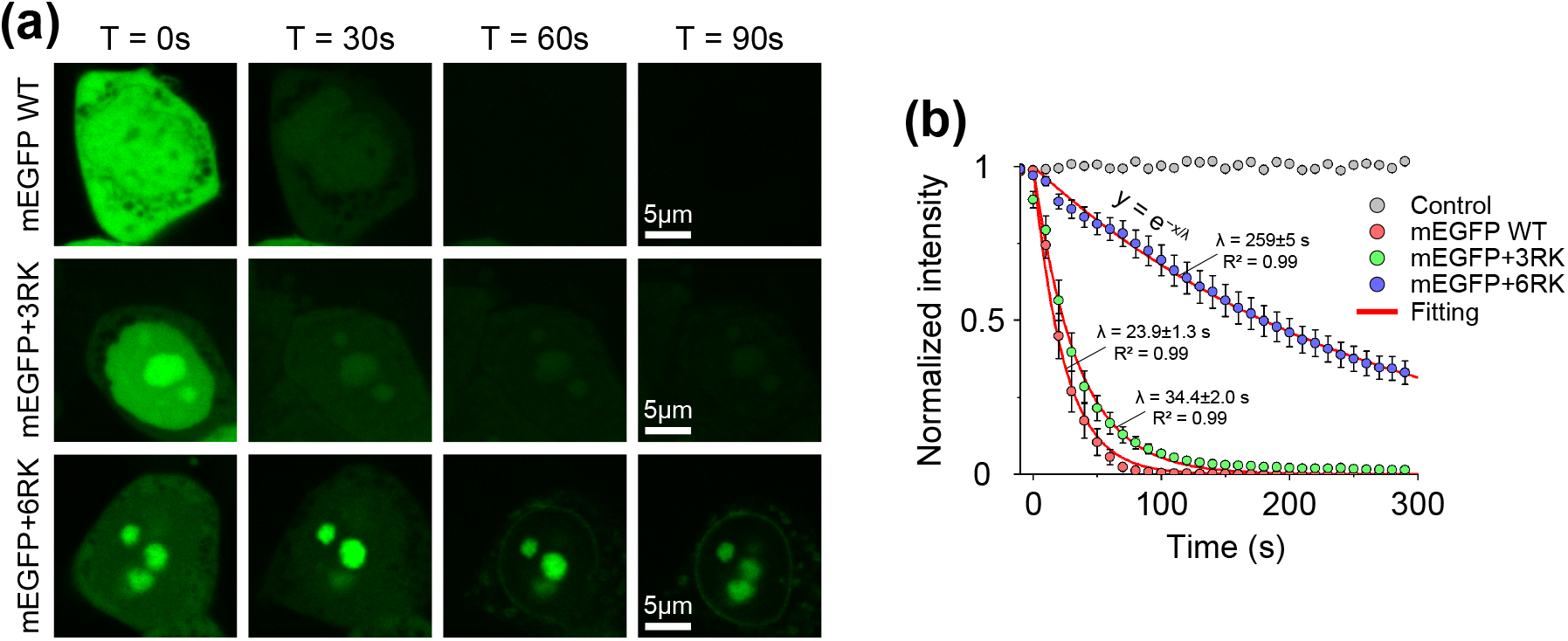
Diffusion rates of different mEGFP probes out of the nuclei of weakly permeabilized HEK293T cells. **(a)** Representative time-lapse sequence showing images of permeabilized HEK293T cells expressing mEGFP WT, mEGFP+3RK and mEGFP+6RK probes. A rapid decrease in the intensity of the mEGFP WT and mEGFP+3RK probes is observed in permeabilized cells compared to the mEGFP+6RK probes. Permeabilization of cells was performed by treating them with PBFT solution, see Methods. **(b)** Total normalized intensity of mEGFP WT, mEGFP+3RK and mEGFP+6RK in the nuclei of permeabilized HEK29T cells as a function of time. The control group included untreated HEK293T cells expressing mEGFP WT. From the experimental data fitting it can be seen that mEGFP+6RK leaves the nucleus at a significantly lower rate compared to mEGFP WT and mEGFP+3RK, which is likely due to a relatively strong binding energy of mEGFP+6RK to various nuclear components, leading to the formation of aggregates that can be seen in panel (a). On the other hand, mEGFP WT and mEGFP+3RK leave the nucleus much faster, indicating that both probes engage in rather weak non-specific interactions with nuclear components. In total, *N* = 16 (control), 22 (mEGFP WT), 15 (mEGFP+3RK) and 19 (mEGFP+6RK) cells were measured in the experiment. Error bars indicate SEM. Fitting parameter values in panel (b) are presented as mean *±* 95% CI.

**FIG. S5.**
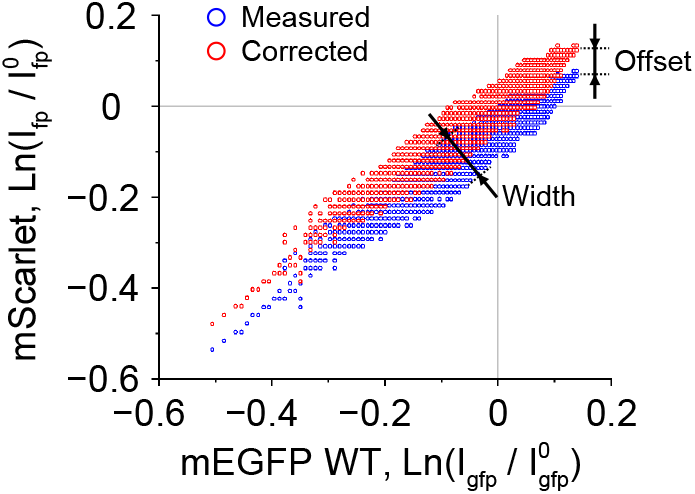
Typical offset between mEGFP WT and mScarlet fluorescent signals measured in HEK293T cell nuclei. The graph shows the logarithm of the normalized mScarlet fluorescent signal 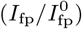 measured in the nucleus of a HEK293T cell coexpressing mEGFP WT and mScarlet plotted as a function of the logarithm of the normalized mEGFP WT fluorescent signal 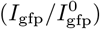. In both cases, normalization was performed to the fluorescent signals of mScarlet 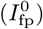 and mEGFP WT 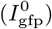 measured in the cytoplasm. The blue data points show the experimentally measured intensities of individual pixels in the image of the nucleus. The red data points represent a hypothetical ideal case where the cloud of data points passes through the origin. The offset between these two data sets and the width of the spread of the data points shown in the figure provide a rough estimate of the *β*[*u*_1_(**r**) *− u*_2_(**r**)] energy term in Eq. (B3) in Appendix B.

**FIG. S6.**
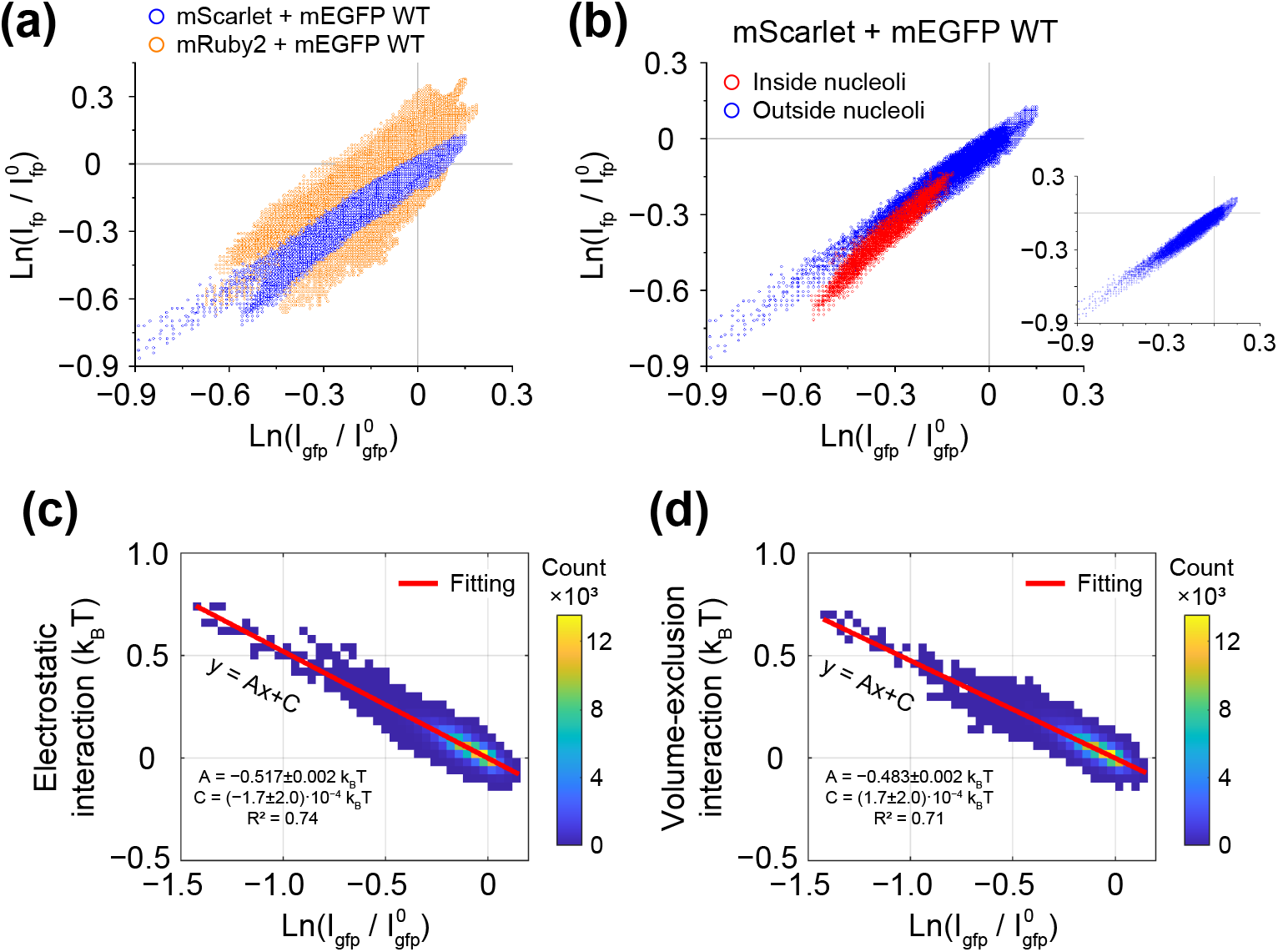
Calibration of the mEGFP WT probe. **(a)** Logarithm of the normalized mScarlet fluorescent signal 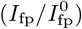 measured in the nuclei of HEK293T cells coexpressing mEGFP WT and mScarlet plotted as a function the logarithm of the normalized mEGFP WT fluorescent signal 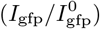. In both cases, normalization was performed to the fluorescent signals of mScarlet 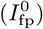 and mEGFP WT 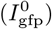 measured in the cytoplasm. Data points represent individual pixels of the nuclei images. For comparison, the graph also shows experimental data obtained on HEK293T cells coexpressing mEGFP WT and mRuby2. In the experiments, cells were grown on fibronectin-coated glass slides. In total, the intensities of *N* = 164550 pixels (mScarlet + mEGFP WT) and *N* = 347009 pixels (mRuby2 + mEGFP WT) were measured in the nuclei of *n* = 13 and *n* = 27 cells, respectively. **(b)** Subdivision of the mScarlet-mEGFP WT data points shown in panel (a) into two groups: those corresponding to pixels measured outside the nucleoli (group 1, blue) and inside the nucleoli (group 2, red) in the cell nuclei. These two groups exhibit distinct behaviours, indicating that mScarlet and/or mEGFP WT engage in different modes of interaction with nuclear components outside and inside the nucleoli. The inset on the right shows only the data points corresponding to group 1; the graph shows a strong linear correlation between the fluorescent signals of mScarlet and mEGFP WT. **(c, d)** Calibration curves showing the average electrostatic and volume-exclusion interaction energies of mEGFP WT with the surrounding nucleoplasmic microenvironment, calculated based from the experimental data presented in the inset of panel (b), as described in Appendix B. A total of *N* = 148880 pixel intensities were measured in the nuclei of *n* = 13 cells. Fitting parameter values in panels (c, d) are presented as mean *±* 95% CI.

**FIG. S7.**
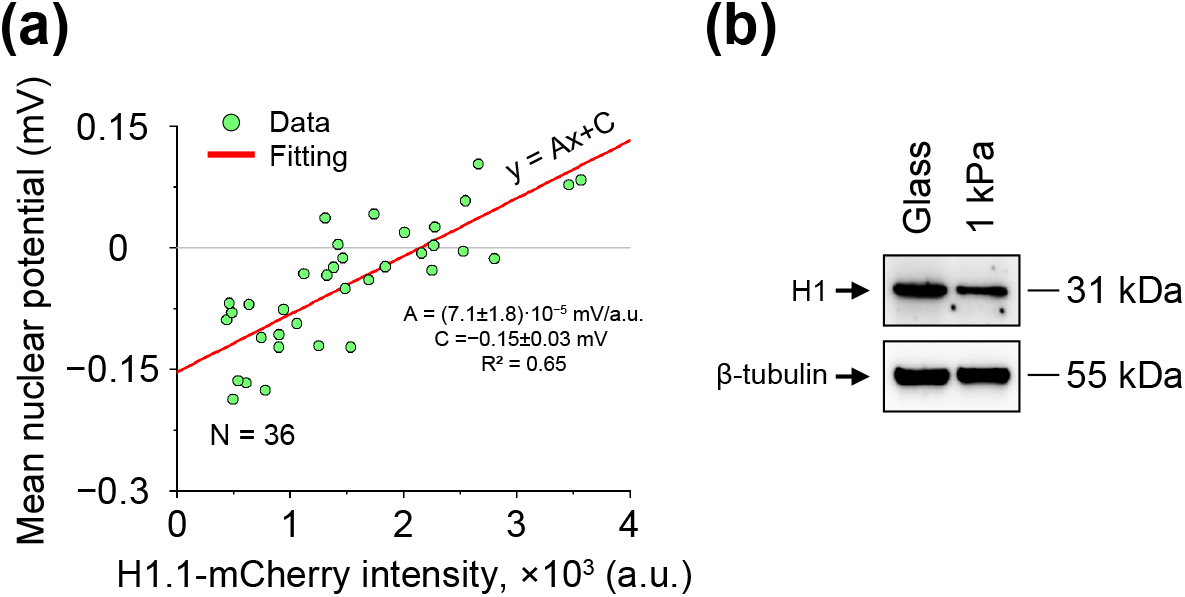
Mechano-dependent expression of H1 linker histones and their effect on nuclear electrostatic potential. Nuclear electrostatic potential of HEK293T cells transfected with a plasmid encoding H1.1-mCherry linker histone. Experimental measurements performed in HEK293T cells coexpressing mEGFP WT and H1.1-mCherry revealed a linear correlation between the mean nuclear electrostatic potential and the average fluorescence intensity of H1.1-mCherry in cell nuclei. This result suggests that supercharged nuclear / chromatin protein complexes, such as H1.1 histones (*q* = +53.0*e*), can significantly influence nuclear electrostatic potential. In total, *N* = 36 cells were measured in the experiment. **(b)** Relative expression levels of endogenous H1 linker histones in HEK293T cells grown on fibronectin-coated glass slides and 1 kPa gel as revealed by Western blotting. The image shows that HEK293T cells grown on 1 kPa gel have approximately half the amount of H1 linker histones as HEK293T cells grown on glass slides. In the experiments, an antibody that binds to all isoforms of H1 linker histone was used. Fitting parameter values in panel (a) are presented as mean *±* 95% CI.

**FIG. S8.**
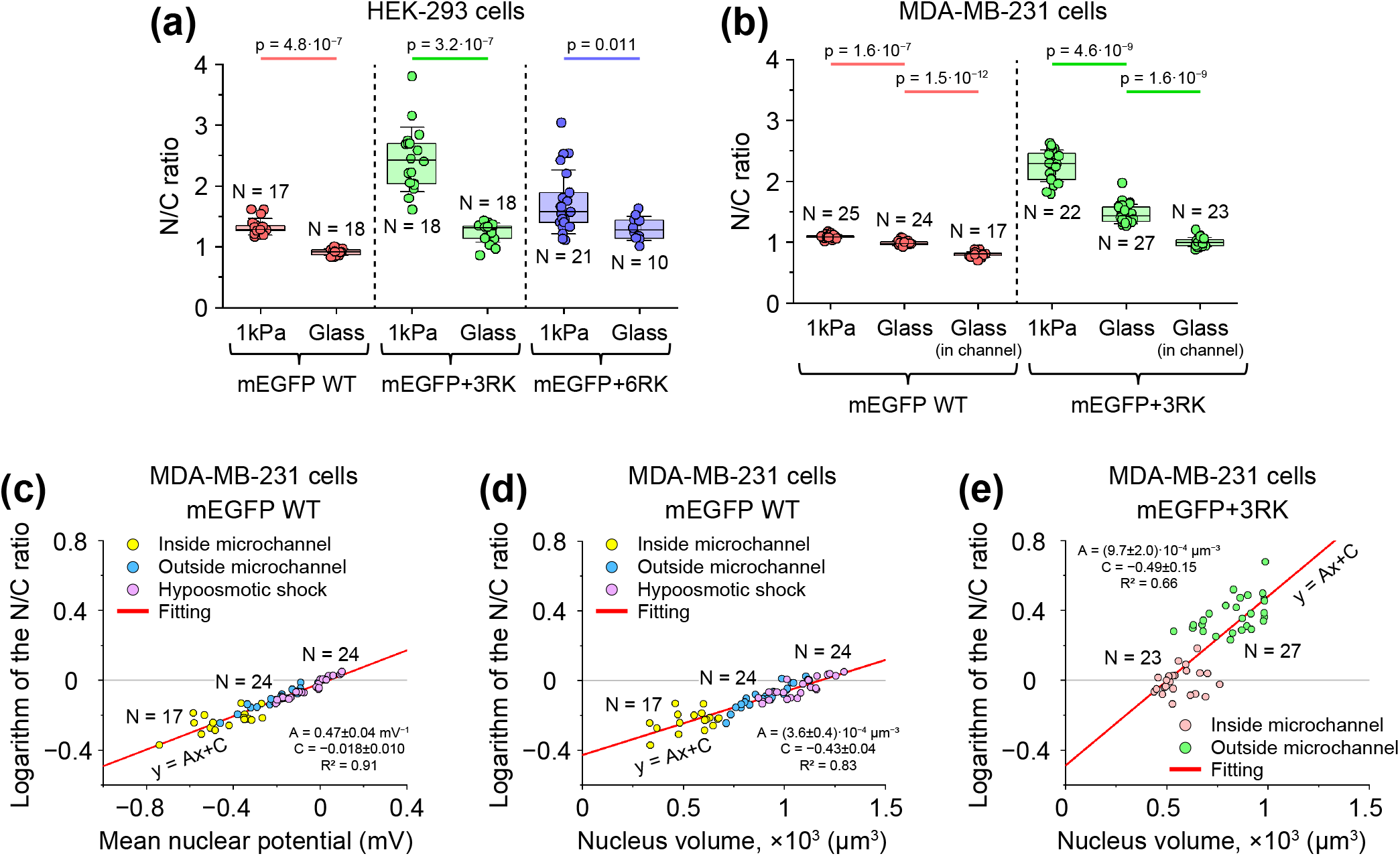
Mechano-dependent behaviour of mEGFP WT and mEGFP+3RK probes. **(a)** Nuclear-to-cytoplasmic (N/C) ratio of mEGFP WT, mEGFP+3RK and mEGFP+6RK probes measured in HEK293T cells grown on fibronectin-coated glass slides and 1 kPa gel. **(b)** N/C ratio of mEGFP WT and mEGFP+3RK probes measured in unconfined MDA-MB-231 cells and MDA-MB-231 cells migrating through 5 *µ*m wide microchannels. For comparison, the N/C ratios of mEGFP WT and mEGFP+3RK probes measured in MDA-MB-231 cells grown on 1 kPa gel are also shown. **(c)** Logarithm of the N/C ratio of the mEGFP WT probe in MDA-MB-231 cells measured in the microchannel cell migration assay and hypoosmotic shock experiment as a function of nuclear electrostatic potential. **(d, e)** Logarithms of the N/C ratios of the mEGFP WT (d) and mEGFP+3RK probes (c) in MDA-MB-231 cells measured in the microchannel cell migration assay and hypoosmotic shock experiment as a function of nuclear volume. In the graphs, *N* indicates the number of cells processed under the corresponding experimental conditions. Boxes represent the interquartile range, error bars indicate SD. Bars above the graph show pairwise statistical differences between corresponding experimental data, calculated using Mann-Whitney test. Fitting parameter values in panels (c-e) are presented as mean *±* 95% CI.

**FIG. S9.**
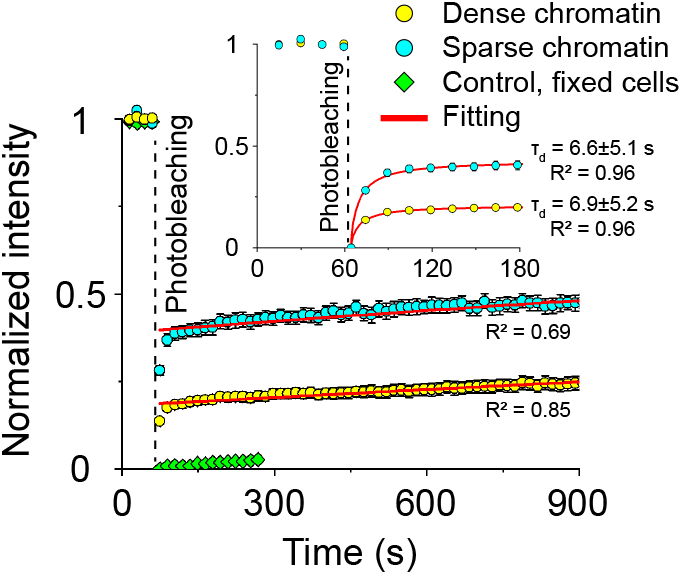
Measurements of the H3.1-mCherry FRAP signal in dense (yellow data points) and sparse (cyan data points) chromatin regions of HEK293T cells. The graph shows the average over *N* = 23 live cells (cyan and yellow data points) and, for comparison, *N* = 8 fixed HEK293T cells (green data points). Cells were grown on fibronectin-coated glass slides. Error bars indicate SEM. The inset displays an enlarged region of the graph before and immediately after the photobleaching. Data fitting was performed as described in Appendix E. Fitting parameter values are presented as mean *±* 95% CI.

**FIG. S10.**
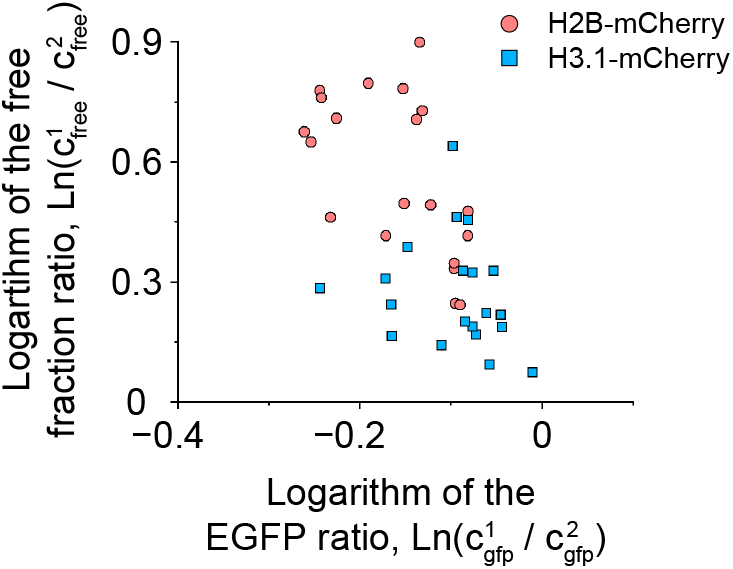
Correlation between the logarithm of the mEGFP signal ratio 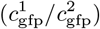 measured for dense (1) and sparse (2) chromatin regions and the logarithm of the density ratio of freely diffusing histone-chaperone complexes 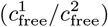 in the same regions. The data points were obtained based on the fitting of the H2B-mCherry and H3.1-mCherry FRAP signals measured in HEK293T cells grown on fibronectin-coated 1 kPa gel, as described in Appendix E. Notably, for cells grown on 1 kPa gel, this correlation is negative for both H2B-mCherry and H3.1-mCherry constructs, whereas for cells grown on glass slides, this correlation is positive, see Figure 4(d). This result could potentially indicate a change in the sign of the net electrical charge of histone-chaperone complexes.

**FIG. S11.**
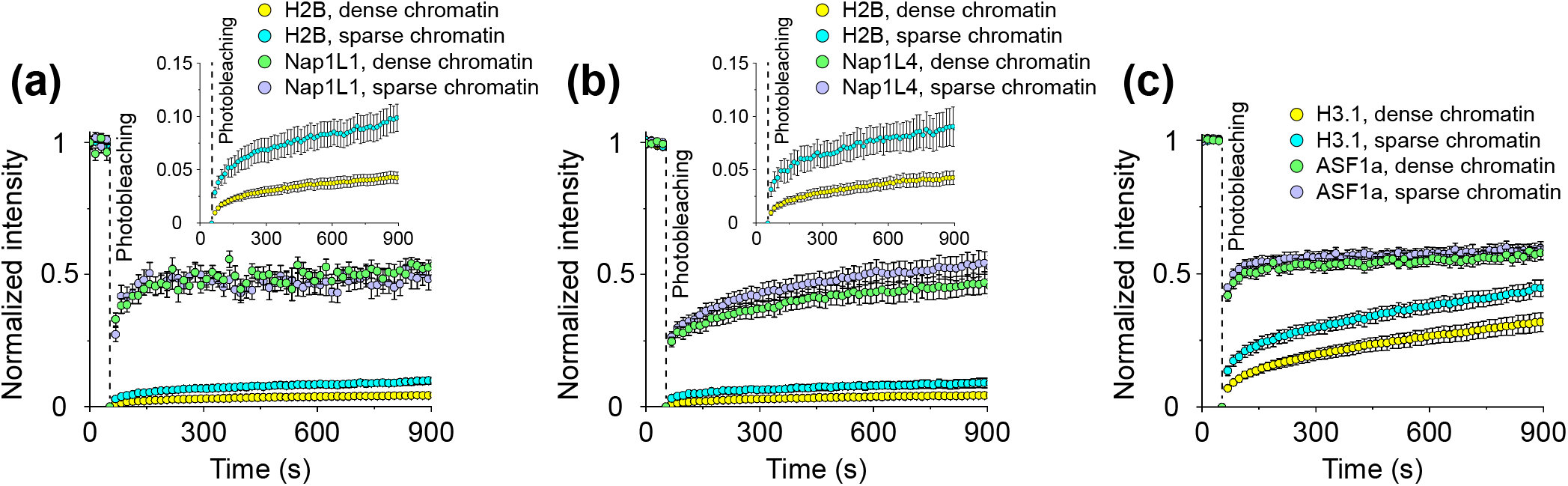
Co-FRAP signals of H2B and H3.1 histones and their chaperones in dense and sparse chromatin regions of HEK293T cells. **(a-c)** FRAP signals of H2B-mCherry and Nap1L1-EGFP (a), H2B-mCherry and Nap1L4-EGFP (b), and H3.1-mCherry and ASF1a-EGFP (c) coexpressed in HEK293T cells. The insets on panels (a, b) show enlarged FRAP H2B-mCherry signals. Comparison of the fluorescent signals of H2B and H3.1 histones and their chaperones shows that the initial rapid recovery of the H2B and H3.1 signals (within the first *∼* 60 s) correlates with the rapid recovery of the fluorescent signals of their chaperones, suggesting that it is mainly due to the diffusion of histone-chaperone complexes into the photobleached region, see also Figures 4(b), S9 and S12. The subsequent, slower recovery of H2B and H3.1 signals over a longer time course is likely due in part to the exchange of fluorescently-labelled histones between histone-chaperone complexes and nucleoprotein complexes formed on DNA. In the experiment, cells were grown on fibronectin-coated glass slides. In total, measurements were performed on *N* = 15 (H2B-Nap1L1), *N* = 12 (H2B-Nap1L4) and *N* = 11 (H3.1-ASF1a) living cells. Error bars indicate SEM.

**FIG. S12.**
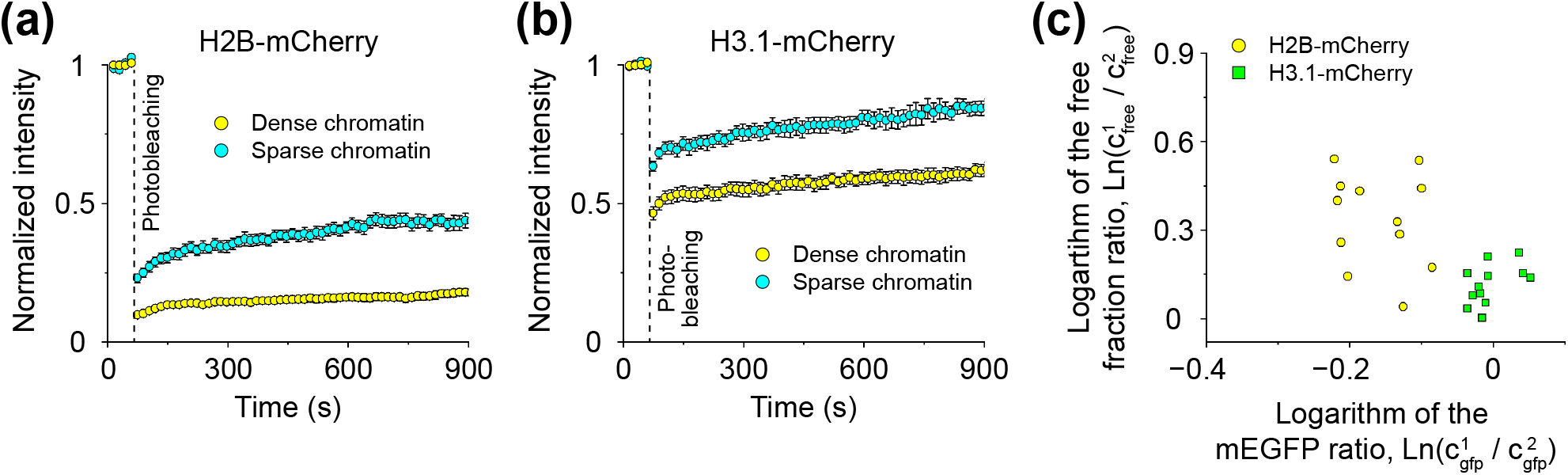
Measurements of FRAP signals of H2B-mCherry and H3.1-mCherry in HEK293T cells arrested in the S-phase of the cell cycle. **(a, b)** H2B-mCherry (a) and H3.1-mCherry (b) FRAP signals measured in dense (yellow data points) and sparse (cyan data points) chromatin regions of HEK293T cells. Cells grown on fibronectin-coated glass slides were arrested in the S-phase by treatment with hydroxyurea for 24 h, see Methods. In total, *N* = 13 cells were measured in both cases. Error bars in the graphs indicate SEM. Comparison of the graphs shown in panels (a, b) to Figures 4(b) and S9 shows that cells arrested in the S-phase have a higher fraction of freely diffusing histones compared to untreated cells (i.e., a stronger initial rapid recovery of histone signals is observed after photobleaching), although the level of freely diffusing H2B histones increases to a lesser extent compared to H3.1 histones. Indeed, in cells arrested in the S-phase, the fraction of freely diffusing H3.1 histones [*∼* 0.5 *−* 0.7, panel (b)] is significantly increased compared to untreated cells [*∼* 0.2 *−* 0.4, Figure S9]. This leads to an increase in the H3/H2B fluorescent signal ratio between sparse and dense chromatin regions by *∼* 14%, as shown in Figure 6(a). Overall, these results are in good agreement with previous experimental studies showing that histone expression levels increase in the S-phase [64]. **(c)** Correlation between the logarithm of the mEGFP signal ratio 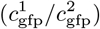 measured for dense (1) and sparse (2) chromatin regions and the logarithm of the density ratio of freely diffusing histone-chaperone complexes 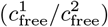 measured in the same regions. The difference between the graphs shown in panel (c) and Figure 4(d) suggests that cell cycle arrest in the S-phase may interfere with the function of histone-binding chaperones.

**FIG. S13.**
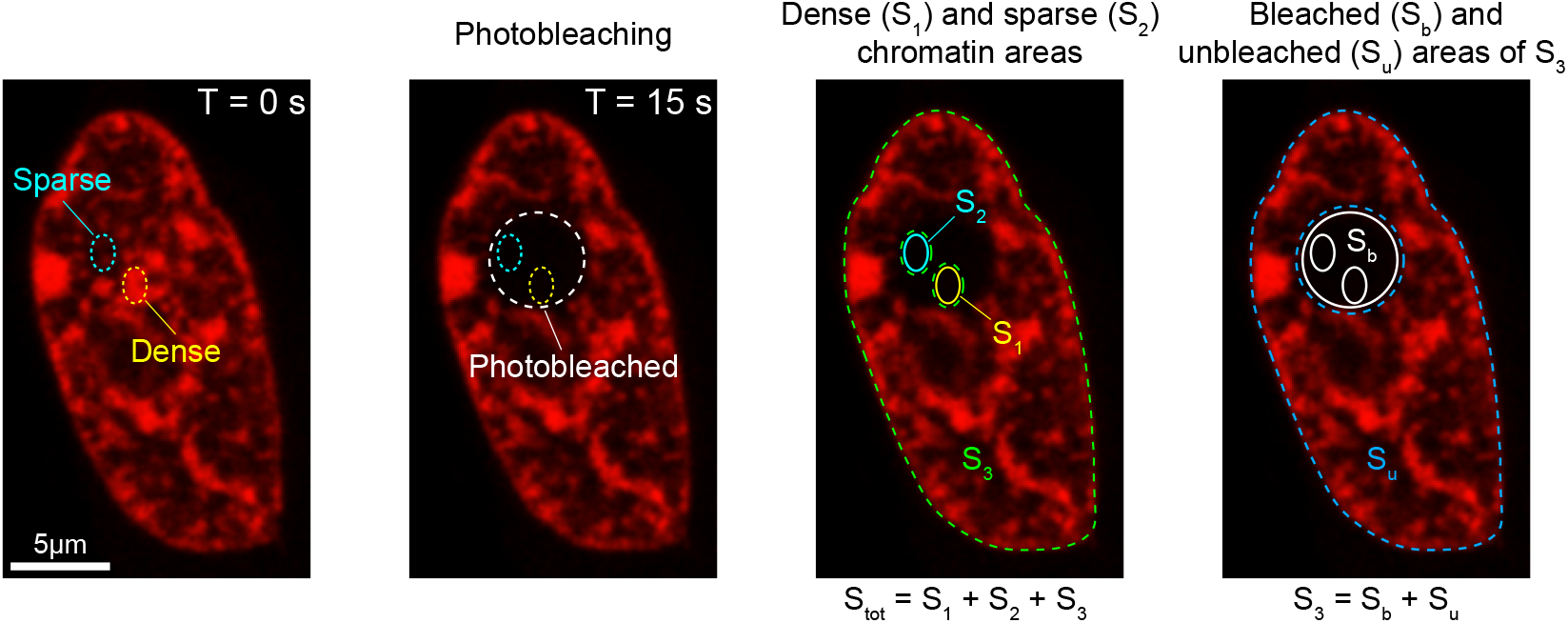
Different regions of the cell nucleus that were used to fit the measured FRAP signals. The fluorescent signal is measured in three distinct regions of the nucleus: dense (*S*_1_) and sparse (*S*_2_) chromatin regions selected within the photobleached nucleus area, and the rest of the nucleus (*S*_3_). For the sake of data processing, the latter is further subdivided into photobleached (*S*_b_) and unbleached (*S*_u_) areas.

**FIG. S14.**
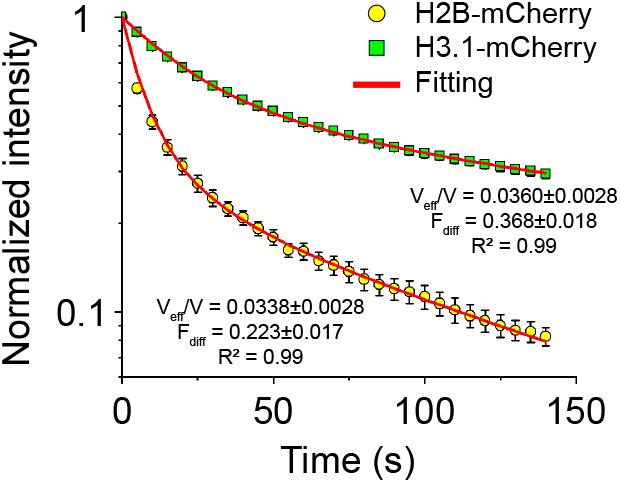
Measurement of the freely diffusing histone fraction in the nuclei of HEK293T cells by continuous fluorescence photobleaching. The graph shows the experimentally measured decrease in fluorescence intensity of H2B-mCherry and H3.1-mCherry constructs expressed in HEK293T cells that were continuously photobleached over a 2 *µ*m-sized nuclear region using a low-intensity laser. The average fractions of freely diffusing H2A *·* H2B histones (0.223 *±* 0.017, *N* = 7 cells) and H3 *·* H4 histones (0.368 *±* 0.018, *N* = 11 cells) were estimated by fitting the collected experimental data to Eq. (9) from ref. [191]. In the experiment, cells were grown on fibronectin-coated glass slides. Error bars indicate SEM. Fitting parameter values are presented as mean *±* 95% CI.

**FIG. S15.**
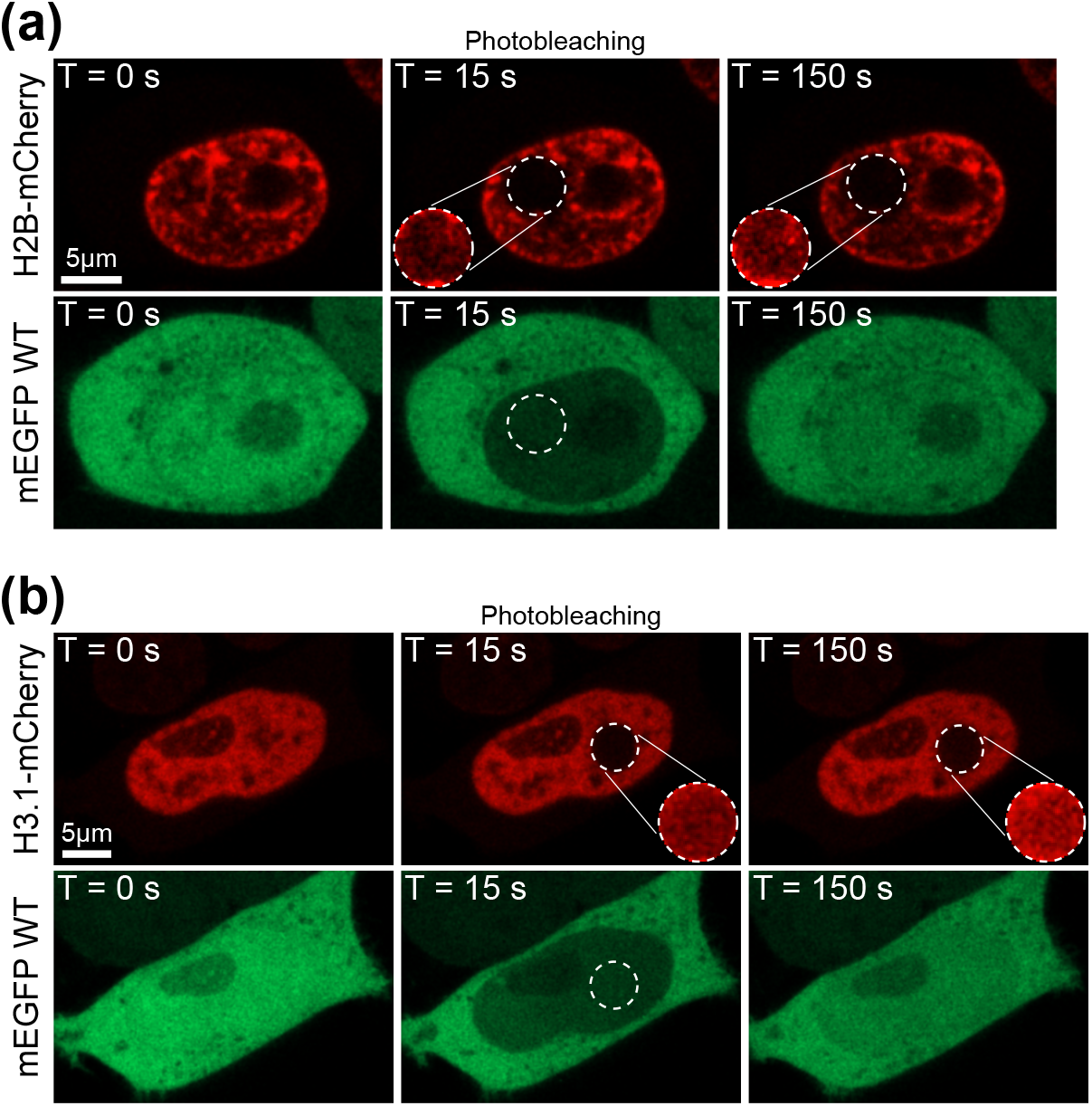
Differences in fluorescence recovery of mEGFP WT, H2B-mCherry and H3.1-mCherry constructs after photobleaching. **(a, b)** FRAP experiments performed on HEK293T cells co-transfected with plasmids encoding mEGFP WT and H2B-mCherry or H3.1-mCherry constructs. The figure shows that mEGFP WT rapidly recovers the fluorescent signal in the photobleached area by diffusion, as after 15 s no significant difference in mean intensity is observed between this region and the rest of the nucleus. In contrast, recovery of the H2B-mCherry and H3.1-mCherry fluorescent signals takes much longer due to the slower diffusion of histone-chaperone complexes and the slow exchange between DNA-bound and chaperone-transported histones, see also Figures 4(b) and S9. In the figure, the dashed circles indicate the photobleached area. In the case of H2B-mCherry and H3.1-mCherry, slow recovery of fluorescence can be observed in the photobleached area, see the enlarged and contrasted inset images of this region.

**FIG. S16.**
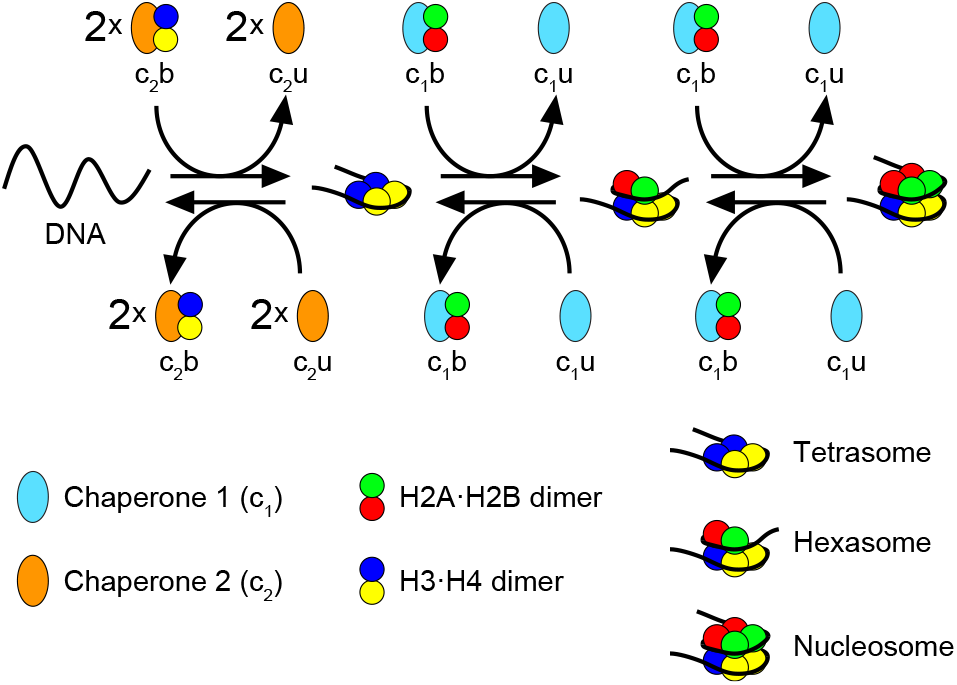
Chemical equation describing nucleosome assembly. Nucleosomes assemble on DNA through several intermediate steps, shown schematically in the figure, during which histone dimers transported by chaperones *c*_1_ and *c*_2_ are deposited onto DNA.

**FIG. S17.**
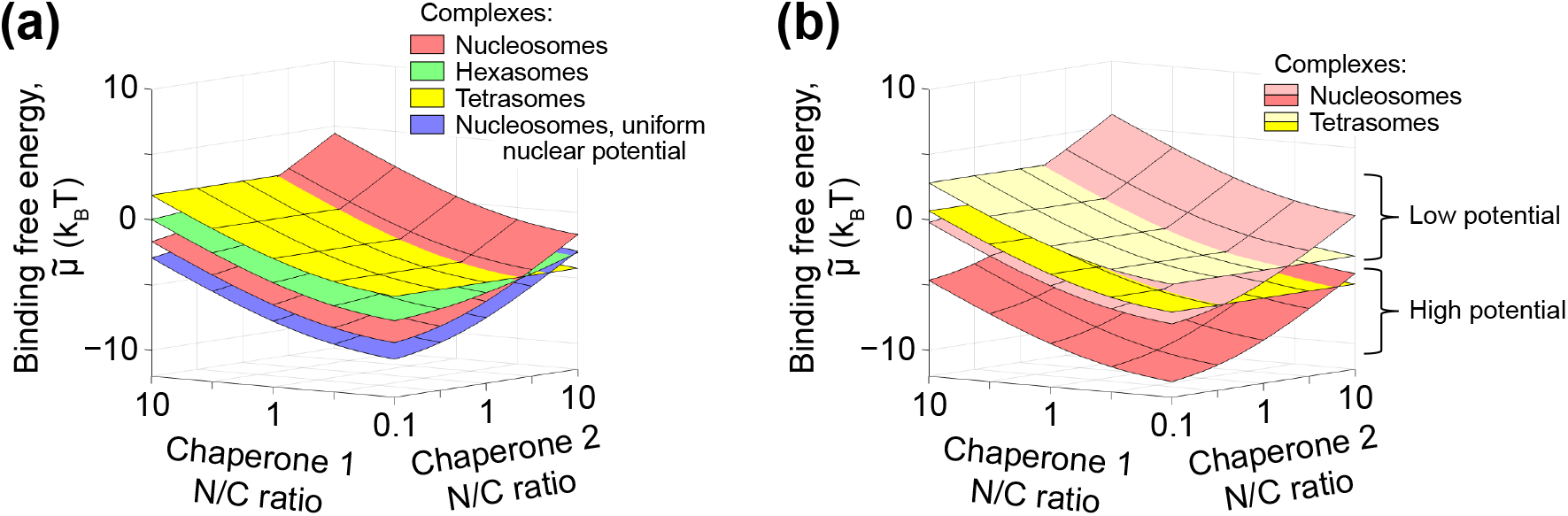
Effect of nuclear electrostatic potential on nucleosome stability in the case of HEK293T cells grown on 1 kPa gel. **(a)** Combined contribution of the nuclear electrostatic potential, weak transient interactions and active nuclear transport to the binding free energy of histone octamers, hexamers and tetramers to DNA as a function of the nuclear-to-cytoplasmic (N/C) ratio of chaperones involved in the transportation of H2A *·* H2B and H3 *·* H4 histone dimers (chaperones 1 and 2, respectively), see Appendix G for details. **(b)** Sensitivity of the binding free energy of histone octamers and tetramers to DNA to heterogeneity of nuclear electrostatic potential. Comparing panels (a, b) and Figure 5(a,b), it can be seen that for HEK293T cells grown on 1 kPa gel, the binding free energy of nucleosomes to DNA in regions with low nuclear electrostatic potential is *∼* 3 k_B_T lower than for HEK293T cells grown on glass. This result suggests that chromatin stability may be somewhat sensitive to the elastic properties of the substrate on which cells are grown, which is in good agreement with previous theoretical predictions [25]. Importantly, in contrast to the case of HEK293T cells grown on fibronectin-coated glass slides, the regions of lower and higher electrostatic potential no longer correlate with heterochromatic and euchromatic domains when the cells are grown on 1 kPa gel, see Figure 2 and main text for details.

**FIG. S18.**
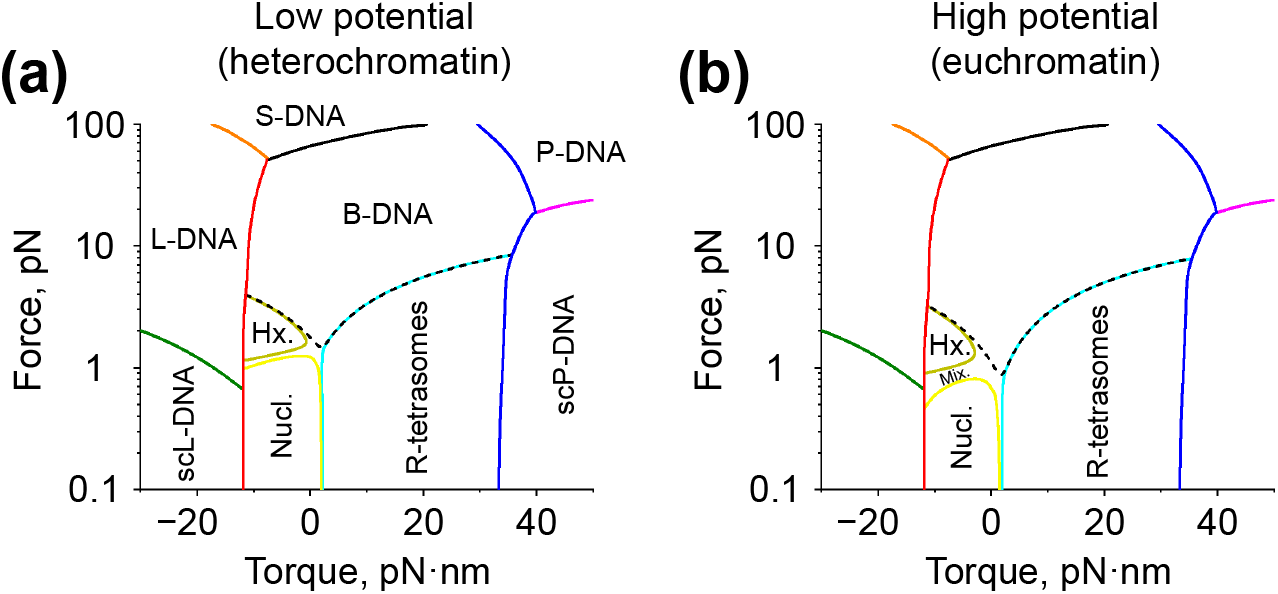
DNA phase diagrams. **(a, b)** Phase diagrams of DNA located in regions of low (a) and high (b) electrostatic potential in the presence of histone-based nucleoprotein complexes (tetrasomes, hexasomes and nucleosomes). The diagrams were calculated based on experimental measurements performed in HEK293T cells grown on fibronectin-coated glass slides, in which regions of low and high electrostatic potential are often associated with heterochromatic and euchromatic domains, respectively (see Figure 2). In the graphs, the solid curves predicted by transfer-matrix calculations represent the transition boundaries between extended (B, L and P) and supercoiled (scL and scP) states of DNA, as well as between different DNA-protein conformations, such as: right-handed tetrasomes (R-tetrasomes), hexasomes (Hx.), nucleosomes (Nucl.) or a mixture of the last two with L-tetrasomes (Mix.), see Appendix H for details. The dashed curve represents the boundary of DNA state in which *>* 50% of DNA is occupied by histone-based nucleoprotein complexes. The figure shows that the difference in stability between euchromatin and heterochromatin, which is mainly due to the difference in the nuclear electrostatic potential of the respective regions, is most pronounced at low torques: approximately twice the stretching force (*∼* 1.5 *−* 2 pN) is required to dissociate most histone-based nucleoprotein complexes from DNA in a heterochromatic region, compared to the lower force (*∼* 0.8 *−* 1.2 pN) required to achieve the same effect in a euchromatic region. This indicates a significant reduction in the stability of histone-based nucleoprotein complexes in euchromatic regions compared to heterochromatic regions. The predicted magnitudes of the forces leading to destabilization of histone-based nucleoprotein complexes are similar to those measured in single-molecule experiments performed in *Xenopus* egg extracts (*∼* 1.2 *−* 1.7 pN in the presence of ATP [48]), indicating good agreement between transfer-matrix calculations and experiment. The values of the model parameters used in the calculations are given in Table T1-T3.

